# Dissecting regulatory syntax in human development with scalable multiomics and deep learning

**DOI:** 10.1101/2025.04.30.651381

**Authors:** Betty B. Liu, Selin Jessa, Samuel H. Kim, Yan Ting Ng, Soon Il Higashino, Georgi K. Marinov, Derek C. Chen, Benjamin E. Parks, Li Li, Tri C. Nguyen, Austin T. Wang, Sean K. Wang, Serena Y. Tan, Michael Kosicki, Len A. Pennacchio, Eyal Ben-David, Anca M. Pasca, Anshul Kundaje, Kyle K. H. Farh, William J. Greenleaf

## Abstract

Transcription factors (TFs) establish cell identity during development by binding regulatory DNA in a sequence-specific manner, often promoting local chromatin accessibility, and regulating gene expression. Mapping accessible chromatin offers critical insights into transcriptional control, but available datasets for human development are restricted to bulk tissue, single organs, or single modalities. Here, we present the Human Development Multiomic Atlas (HDMA), a single-cell atlas of chromatin accessibility and gene expression from 817,740 fetal cells across 12 organs, spanning 203 cell types and over 1 million candidate cis-regulatory elements, many of which exhibit organ-specific *in vivo* enhancer activity. Deep learning models trained to predict accessibility from local DNA sequence unravel a comprehensive lexicon of motifs which influence accessibility, including composite motifs exhibiting distinct syntactic constraints predicted to mediate TF cooperativity. We identify “hard” syntactic rules requiring precise motif spacing and orientation, “soft” rules allowing flexible motif arrangements, and ubiquitous motifs that inhibit accessibility. Model-based interpretation of genetic variants reveals that disruption of motifs with positive and negative effects is associated with concordant effects on gene expression. Our work delineates how motif syntax governs cell type-specific chromatin accessibility and provides a foundational resource for decoding cis-regulatory logic and interpreting genetic variation during human development.

## Introduction

During human development, a single zygote gives rise to the full diversity of human cell types. Although somatic cells share a near-identical genome, cell identity emerges through differential expression and activity of transcription factors (TFs), which integrate cell-intrinsic and -extrinsic signals to direct gene regulatory programs^1^. Once expressed and appropriately localized intracellularly, TFs bind specific sequences of DNA in cis*-*regulatory elements, often inducing regions of local chromatin accessibility, and altering the expression of downstream genes^2^. However, a comprehensive view of the TF motifs capable of driving chromatin state changes during human development remains incomplete, as does our understanding of how the organization—or syntax—of TF binding sites contributes to cell type-specific regulatory activity^3^.

Mapping chromatin accessibility using DNase-seq and ATAC-seq^2,4^ has enabled the inference of regulatory activity of TFs via sequence motifs in human tissues^5,6^. However, bulk assays are confounded by cellular heterogeneity, and recent single-cell atlases have largely focused on individual organs^7,8^ or single “omic” modalities^9,10^. Given the pleiotropic roles of many TFs across tissues, a multi-organ, multi-modal view is needed to fully capture the cell context-specificity of cis-regulation. In particular, simultaneous measurement of chromatin accessibility and gene expression within the same cell is critical to directly link regulatory element activity to transcriptional programs.

Chromatin accessibility at regulatory elements often arises from cooperative TF binding, either through direct TF–TF interactions^11,12^ or competition with nucleosomes^13–15^. These mechanisms respectively impose either “hard” syntactic constraints—fixed spacing and orientation of motifs— or “soft” constraints that allow flexible motif organization. Yet the extent and generality of such syntactic rules across human developmental contexts remain poorly understood. Furthermore, genetic variants associated with traits and disease risk are highly enriched in the non-coding genome and likely disrupt regulatory sequence syntax^16^. However, our ability to predict which non-coding variants disrupt regulatory activity, and in which cell types, remains limited.

Recent deep learning models trained to predict base-resolution chromatin accessibility profiles from local DNA sequence have demonstrated the ability to learn causal sequence features that influence accessibility^17–21^. Beyond *de novo* discovery of predictive motifs and TF footprints, these models also enable systematic interrogation of regulatory sequence syntax by predicting the quantitative effects of modifying motif composition, spacing, and orientation on accessibility *in silico*. Similarly, these models can predict and interpret the effects of non-coding genetic variants on accessibility^17,22^. Thus, deep learning models provide a powerful framework for decoding the combinatorial logic of how TF binding influences chromatin accessibility and for linking sequence variation to disruption of cis-regulation.

Here, we present the Human Development Multiomic Atlas (HDMA), a true multiomic, multi-organ single-cell atlas profiling chromatin accessibility and gene expression in 817,740 primary human fetal cells across 12 organs. We mapped over one million accessible regulatory elements across 203 annotated cell types, and demonstrated their ability to resolve organ- and cell type-specific enhancer activity *in vivo*. We trained and interpreted deep learning models of cell type-resolved accessibility profiles from HDMA, to define a comprehensive lexicon of regulatory sequence motifs driving accessibility—including ubiquitous, cell type-specific, and novel elements—and inferred on average over eight hundred thousand predictive motif instances in the genome per cell type. Systematic interrogation of motif syntax uncovered both “hard” and “soft” syntactic constraints. Interpreting genetic variation through these models, we found that disruption of positive or negative motifs predicted concordant effects on gene expression, and prioritized disease-associated variants that likely perturb TF binding sites and downstream regulatory function in developmental contexts. Altogether, HDMA provides a foundational resource for decoding cis-regulatory syntax, linking sequence variation to gene regulation, and understanding how chromatin accessibility patterns drive human cell type diversity.

## Results

### A multiomic, multi-organ atlas of human development

We simultaneously profiled chromatin accessibility and gene expression from 817,740 fetal cells spanning 12 organs between post-conception weeks 10 and 23 using SHARE-seq^23^, a scalable split-and-pool combinatorial barcoding platform (**Fig. 1a, Fig. S1, Fig. S2a**). The resulting Human Development Multiomic Atlas (HDMA) captures true multiomic measurements from the same cells, with data quality for both modalities exceeding that of previous multi-organ fetal “single-ome” atlases (**Fig. 1b**, **Fig. S2b-c, Table S1**)^9,10^.

**Figure 1.**
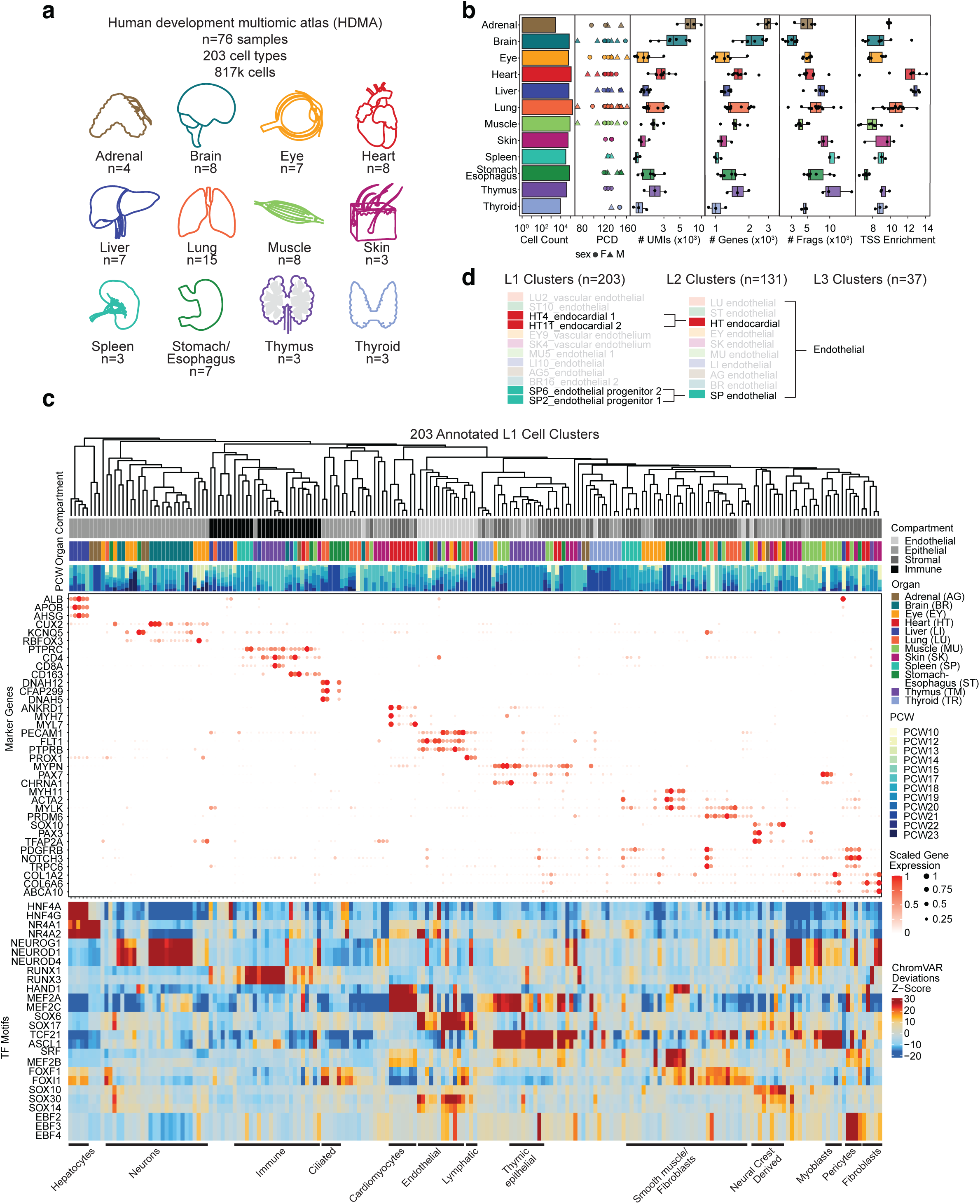
A multiomic, multi-organ atlas of human development. **a)** HDMA dataset overview and sample size distribution by organ. **b)** Sample meta data and quality control metrics for both RNA and ATAC modalities, with samples ranging from PCW 10-23. **c)** Key marker gene expression and TF motif chromVAR deviation z-score for 203 hierarchically clustered cell types. **d)** An example of different levels of cell annotation for endothelial cells. PCD=post-conception days, UMI=unique molecular identifiers, Frags=ATAC fragments. PCW=post-conception weeks.

We annotated 203 distinct cell clusters (L1 clusters) through an iterative approach combining canonical compartment markers, known cell type markers, and *de novo* marker genes from our single-cell transcriptomes (**Methods**). Cell identity was further corroborated by significant TF motif enrichments within cluster-specific accessible chromatin peaks (**Note S2**).

Hierarchical clustering of pseudobulk expression profiles for the top marker genes revealed that cell types common to different organs—such as endothelial cells, fibroblasts, and immune cells— often clustered together (**Fig. 1c**). Based on shared marker gene expression, we manually grouped related L1 clusters into 134 collapsed cell clusters (L2 clusters) and further into 37 broad cell classes (L3 clusters) (**Fig. 1d, Table S2, Table S3**).

Global chromVAR^24^ deviation analysis identified TF motifs with elevated accessibility in each L3 cell cluster (**Fig. 1c**). Compared to marker genes, TF motif accessibility variations were generally less specific, with many motifs exhibiting pleiotropic accessibility patterns across cell types. For example, while expression of the *CUX2* marker gene was restricted to neurons, the top motif in neurons, NEUROD1, showed increased accessibility not only in neurons but also in neural crest-derived and fibroblast cell types, reflecting motif degeneracy and the context-dependent activity of TFs during development.

Together, these results define a high-quality multiomic, multi-organ single-cell atlas of human development, establishing a foundation for dissecting cis-regulatory logic across diverse developmental contexts.

### Accessible DNA elements reveal the regulatory landscape of human development

We cataloged the accessible regulatory landscape of human development by aggregating peaks independently called across the 203 L1 cell types. Iterative overlap of peaks defined a global set of 1,032,273 <u>a</u>ccessible <u>c</u>andidate <u>c</u>is-regulatory <u>e</u>lements (acCREs), each spanning 500 bp, collectively covering ∼17% of the genome. Most acCREs had their highest normalized ATAC signal in liver, eye, heart, and stomach/esophagus cell types (**Fig. S2d**).

Comparison with the ENCODE v4 database^25^ revealed that 85.1% of HDMA acCREs overlapped previously characterized cCREs (**Fig. S2e**). Notably, we recovered 48.7% of all ENCODE v4 sites, including 56.2% of CTCF-bound elements. Among the 14.9% of acCREs not overlapping ENCODE cCREs, brain- and eye-derived elements were disproportionately represented, suggesting that single-cell profiling uncovers cell type-specific regulatory elements absent from ENCODE bulk-derived datasets (**Fig. S2d**).

We linked acCREs to genes using the Activity-By-Contact (ABC) model^26^ applied within each L1 cell cluster. For each broad L3 cluster, we then aggregated ABC links from all its constituent L1 clusters and filtered for elements targeting genes with accessible promoters. In each L3 cluster, we defined highly regulated genes (HRGs) as the top 1% of genes with the most ABC-linked acCREs (**Table S4**). Gene Ontology (GO) analysis revealed that HRGs were enriched for specialized cellular processes consistent with the function of the cell types, such as “B cell proliferation” in immune cells, “keratinocyte differentiation” in keratinocytes, and “camera-type eye morphogenesis” in pigmented epithelial cells (**Fig. 2a**). Transcriptional regulation-related terms were consistently enriched across 25 of 37 L3 clusters. Further GO molecular function analysis confirmed that HRGs were highly enriched for transcription factor binding activity across 33 of 37 L3 clusters (**Fig. S2f**). For example, endothelial HRGs included canonical regulators such as *NR2F2*, *ELF4*, *SOX18*, *NFIC*, and *FOXC1*, transcriptional coregulator *BCL9L*, as well as non-TF genes related to endothelial functions such as *EGFL7*, *PLXND1* and *PLEC*. The top 10 GO terms enriched in endothelial HRGs included both general TF activity terms and endothelial-specific terms such as PDGFR binding (**Fig. 2b-c**).

**Figure 2.**
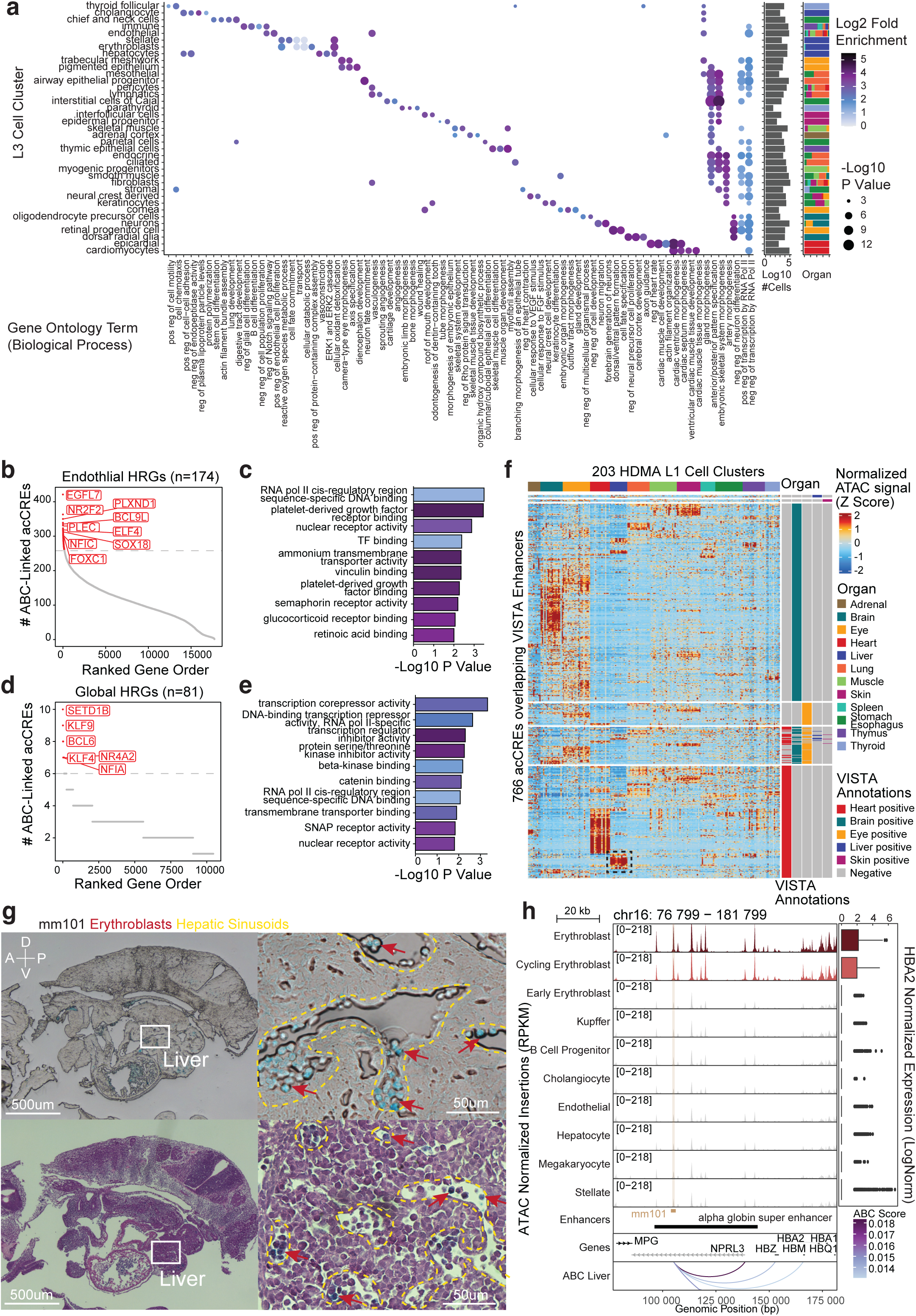
Connecting DNA regulatory elements to genes and resolving organ- and cell type-specificity in experimentally-validated enhancers. **a)** Top 5 Gene Ontology biological process terms enriched in the highly regulated genes (HRG) in each broad class cell cluster (L3 cell cluster). pos=positive, neg=negative, reg=regulation. **b)** Endothelial and **d)** global HRGs with the most ABC-linked acCREs. Gene Ontology molecular function term enrichment in **c)** endothelial HRGs and **e)** global HRGs. **f)** Normalized and Z-scored ATAC signal in the HDMA acCREs overlapping VISTA enhancers. **g)** Bright field (top) and H&E (bottom) staining images of VISTA embryo mm101 sections. Blue color is from X-Gal staining indicating where the enhancer is active. A=anterior, P=posterior, D=dorsal, V=ventral. **h)** Accessibility at VISTA enhancer mm101 and ABC-linked alpha-globin gene *HBA2* expression in liver cell types.

To identify conserved regulation by acCREs across cell types, we filtered for acCRE-gene links consistently identified in all L3 clusters. The resulting set of global highly regulated genes (gHRGs), corresponding to the top 1% of genes with the most consistent ABC-linked acCREs, was also significantly enriched for transcription factor activity (**Fig. 2d-e**), and included ubiquitous regulators such as *KLF9, KLF4, and BCL6*. These 81 gHRGs constitute a set of ubiquitously highly regulated TFs and chromatin factors across cell type development (**Table S4**).

Our results support the hypothesis that TFs are among the most highly regulated genes within each cell type, consistent with our previous observations in the developing human brain^7^. While many TFs exhibit dynamic, cell type-specific regulation, we also identified a subset of TFs that are consistently highly regulated across diverse developmental contexts, suggesting ubiquitous roles in maintaining cellular state.

### Resolving organ- and cell type-specificity in experimentally-validated enhancers

To assess whether HDMA recapitulates known enhancer activity patterns, we examined chromatin accessibility at acCREs overlapping experimentally validated enhancers from the VISTA database, assayed by X-gal reporter staining at mouse embryonic day 11.5^27–29^. Among VISTA enhancers annotated as active in brain and heart, we observed strong enrichment of both accessibility (Wilcoxon rank-sum test, p<2.2×10^-16^, AUROC probability=0.72 for brain and 0.75 for heart) (**Fig. 2f**) and gene expression (Wilcoxon rank-sum test, p<2.2×10^-16^, AUROC probability=0.57 for brain and 0.67 for heart) (**Fig. S3a**) in the HDMA cell type clusters from the corresponding organs.

Surprisingly, we also detected high chromatin accessibility in organs where VISTA had not originally annotated enhancer activity. For example, several enhancers previously annotated as heart-specific exhibited strong accessibility exclusively in liver cell types (dashed black box, **Fig. 2f**). Given that the mouse liver lies directly beneath the heart and is often naturally colored, these cases may have been missed or mislabeled by visual inspection alone. VISTA annotations of our top candidate enhancers postulated as active in the liver were expertly re-reviewed, confirming six enhancers (mm2143, mm69, mm291, mm101, mm257, mm18) as active in both liver and heart^27^. Sectioning and imaging of X-gal stained embryos validated enhancer activity within the liver for all six candidates (**Fig. 2g, Fig. S3b-f**).

One such enhancer, mm101, overlapped the known alpha-globin super-enhancer locus^30–33^ and was predicted by ABC analysis to regulate the alpha-globin gene cluster (*HBZ, HBM, HBA2, HBA1, HBQ1*) (**Fig. 2h**). Alpha-globin is an important subunit of fetal hemoglobin present exclusively in erythroblasts^34^. Given the role of the fetal liver as a site of erythropoiesis during development, we hypothesized that mm101 enhancer activity would be restricted to erythroblasts. Indeed, accessibility at the mm101 locus and expression of HBA2 were both elevated specifically in liver erythroblasts and cycling erythroblasts (**Fig. 2h**). Histological analysis confirmed X-gal positivity in liver erythroblasts characterized by a globular morphology and characteristic darkened purple color from H&E staining, which reside within hepatic sinusoids (blood vessel spaces) (**Fig. 2g**).

These results demonstrate that many of our organ- and cell type-specific enhancers are capable of driving chromatin accessibility and gene expression *in vivo* in relevant tissues during development.

### *De novo* discovery of the lexicon of motifs influencing chromatin accessibility

To identify cis-regulatory sequence features predictive of chromatin accessibility, we trained deep convolutional neural networks (ChromBPNet) to model the shape and magnitude of ATAC-seq profiles from local DNA sequence^17,18^. For each cell type, we trained models to predict both total read counts and base-resolution distribution of reads in 1,000 bp windows centered on relaxed pseudobulk ATAC peaks and GC content-matched background regions, using 2,114 bp of local sequence context as input (**Fig. 3a**, **Methods**). We trained five-fold cross-validated models for 203 cell types and retained models for 189 cell types (N=945 total models) that achieved a median Pearson correlation of 0.78 between predicted and observed log-read counts in held-out peaks in test chromosomes (**Fig. 3b-c, Fig. S4a-b, Table S5, Note S2**).

**Figure 3.**
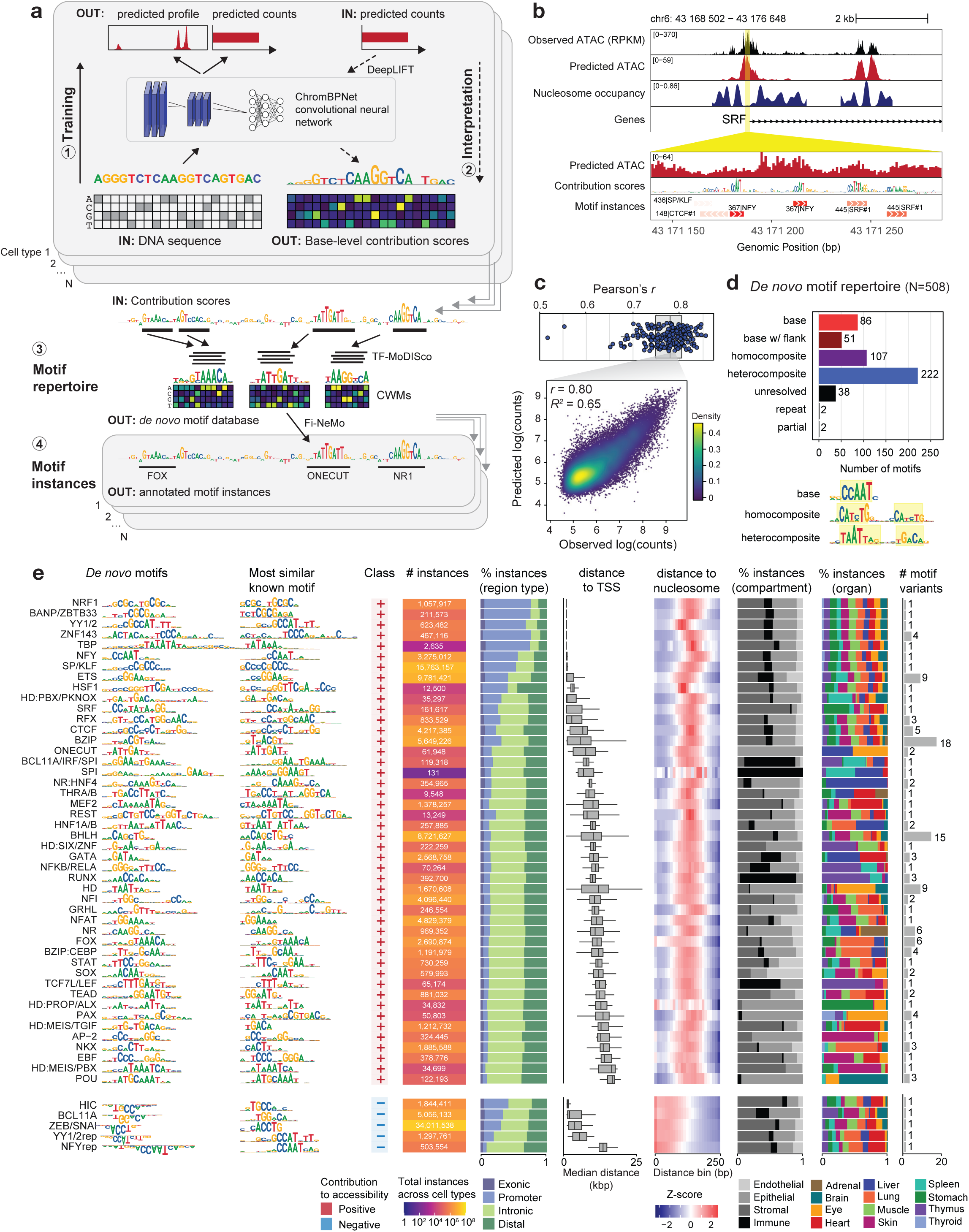
*De novo* discovery of the repertoire of motifs in human development. **a)** Overview of workflow. For each cell type, a ChromBPNet model is trained to predict accessibility from local DNA sequence. Models are interpreted to derive per-base scores representing contribution to accessibility. High-contribution regions are clustered within and across cell types into motifs, and motifs are used to annotate high-contribution regions in each cell type. **b)** Example of *SRF* locus in cardiomyocytes showing observed and predicted chromatin accessibility, inferred nucleosome occupancy, per-base contribution scores, and annotated motif instances. **c)** Top: Distribution of Pearson correlation between predicted and observed log counts in peak regions per model (mean across five folds). Bottom: predicted vs observed log counts in peak regions for one cell type, one model fold. **d)** Summary and categorization of unique *de novo* motifs, and example motifs from select categories. **e)** Summary of base motifs, one row per broad group of base motifs. From left to right: *de novo* motif representation as a contribution weight matrix (CWM), most similar known motif (PWM), direction of contribution to accessibility, total number of genomic instances across cell types, proportion of instances overlapping various genomic features, distribution of median distance of instances (per cell type) to nearest transcription start sites (TSS), proportion of instances from cell types in each tissue compartment, proportion of instances from cell types in each organ, and number of motif variants within each broad motif group.

We asked which *cis*-regulatory sequence features influence accessibility in each cell type. We expect these sequences to correspond to sites bound by TFs, which protect nucleotides they occlude from Tn5 cleavage, but promote local accessibility in their vicinity^2^. To interpret predictive sequence features in each peak from each cell type, we interrogated the corresponding model with the DeepLIFT algorithm^35^ to estimate the contribution of each base-pair in the peak sequence to the predicted accessibility (**Fig. 3a**). Next, in each cell type, we used TF-MoDISco^36^ to identify subsequences with high contribution scores within peaks, and then align and cluster similar subsequences into a set of motifs. Motifs are represented by contribution weight matrices (CWMs) derived from the average contribution scores of all constituent predictive subsequences. Further aggregation of motifs across all cell types yielded a comprehensive, non-redundant lexicon of 508 *de novo* sequence motifs predicted to regulate chromatin accessibility across 189 cell types (**Fig. 3a,d,e**, **Methods**).

We annotated these *de novo* motifs based on the TFs likely to bind them, typically at the TF-family level given that many TFs within families recognize similar sites (**Methods, Table S6**). Several motifs matched known TF binding motifs in existing databases (“base” motifs) or matched known motifs with additional predictive flanking nucleotides (“base with flanks”) (**Fig. 3d**). The motif lexicon also included several variants of canonical motifs; for example, the models recovered multiple CTCF motifs, including the core motif and an upstream variant, reflecting DNA contacts by distinct CTCF zinc fingers^5,37^ (**Fig. S4c**). More than half of the *de novo* motifs were composite motifs, comprising either two homotypic sites (“homocomposites”) or heterotypic combinations of different sites (“heterocomposites”). A subset of 38 motifs showed low similarity to known databases and were classified as “unresolved.” Most motifs (N=493, 97%) had positive contribution scores, indicating they promote chromatin accessibility, while a small subset had negative contribution scores, suggesting they reduce accessibility (**Fig. 3e, Table S6**). We refer to these as “positive” and “negative” motifs, respectively, based on their predicted directional influence on accessibility.

Next, we used Fi-NeMo^38^ to identify predictive instances of each motif (putative TF binding events) in accessible peaks of each cell type (**Fig. 3a**, **Methods**). On average, each cell type harbored 839,544 motif instances, with 2–8 instances per peak and 65% of instances falling within 150 bp of peak summits (**Fig. S4e-i**). The number of motif instances scaled with the total number of peaks in each cell type, suggesting potential for further sensitivity with deeper sequencing for cell types with less coverage (**Fig. S4j**).

As an illustrative example, we revisited the mm101 enhancer active in fetal liver erythroblasts. Model interpretation revealed two high-scoring GATA motif instances within the enhancer (**Fig. S4k**), consistent with the essential role of GATA1 in erythropoiesis^39^ and supporting its contribution to alpha-globin gene activation in this context^40^.

We next stratified predictive motif instances by genomic context and cell type specificity. Each instance was annotated as promoter-proximal, intronic, exonic, or distal (all others), and assigned distances to the nearest transcription start site (TSS) and nucleosome dyad inferred from the ATAC modality using NucleoATAC^41^ (**Fig. 3e**). We also analyzed the distribution of motif instances across tissue compartments and organs. This analysis revealed a set of promoter-dominant, TSS-proximal motifs used ubiquitously across organs, including NRF1, NFY, YY1/YY2, TBP, SP/KLF, ETS, and the recently characterized BANP. These motifs are generally CG-rich, consistent with the CpG-rich nature of promoters, and NRF1 and BANP are known to be DNA methylation-sensitive TFs^42,43^. Many of these same motifs have been implicated in regulating transcription initiation^44–46^. Other motifs that were also broadly used across organs occurred in both promoter-proximal and distal or intronic regions. These included CTCF, which binds at promoters and distal elements to mediate 3D chromatin contacts^37^, as well as BZIP, RFX, and HSF1 motifs. In contrast, organ- and cell type-specific motifs were predominantly located in distal or intronic regions. These included SPI and RUNX (immune cells), POU (eye and brain), and HNF4 (liver) (**Fig. 3e**). In cell types such as endothelial cells which arise in several different organs, independently trained models learned consistent motifs (**Fig. S5a**), suggesting robust discovery of regulatory lexicons across related cell contexts. Among unresolved motifs lacking matches to known databases, we also identified two palindromic elements with restricted usage in stomach, lung, and liver (**Fig. S5b**).

Together, this analysis defines a comprehensive motif lexicon across fetal cell types learned directly from sequence-to-accessibility models, revealing ubiquitous, cell type-specific, and previously unannotated motifs with distinct positional preferences relative to gene promoters and nucleosomes.

### Systematic inference of transcription factor synergy reveals hard and soft motif syntax

Combinatorial binding of transcription factors at regulatory elements can enhance specificity and function, often through cooperative interactions. In DNA-mediated cooperativity, TFs with direct protein-protein interactions bind a specific composite recognition site, or alternatively, DNA at such sites stabilizes weak interactions between TFs^11,12^. In contrast, nucleosome-mediated cooperativity could arise from active or passive competition between TFs and nucleosomes^13,14^. We reasoned that DNA-mediated cooperativity imposes fixed spacing and orientation constraints on binding sites (defined here as hard syntax) within protein-protein interaction length-scales (<20 bp), whereas nucleosome-mediated cooperativity is compatible with flexible binding site organization (soft syntax) and longer distances between sites (20-150 bp)^47^.

To systematically identify synergistic TF interactions and infer syntactic constraints across cell contexts, we implemented an *in silico* marginalization framework using our trained ChromBPNet models and the tangermeme package^48^, which exhaustively evaluates the joint effects of two motifs across various spacing or orientation arrangements, compared to their independent effects^17,19,49^ (**Fig. 4a-b, Methods**). For each composite motif identified *de novo* (**Fig. 3d**), we inserted its constituent motifs into 100 neutral inaccessible background sequences at all possible pairwise distances (0–200 bp) and orientations, and predicted accessibility using the trained model for the cell type with the most predictive instances of the composite motif (**Fig. 4a-b, Methods**). We quantified joint effects as the increase in predicted log counts for the edited sequences relative to background sequences, and defined synergy as a significant deviation from a log-additive model of independent motif effects. We classified composite motifs as exhibiting hard syntax if the maximal joint effect of the constituent motifs at any specific spacing and orientation was at least 4 standard deviations above the mean across all arrangements. Soft syntax was assigned when joint effects exceeded additive expectations within a range of relative distances between 20–150 bp (**Methods**).

**Figure 4.**
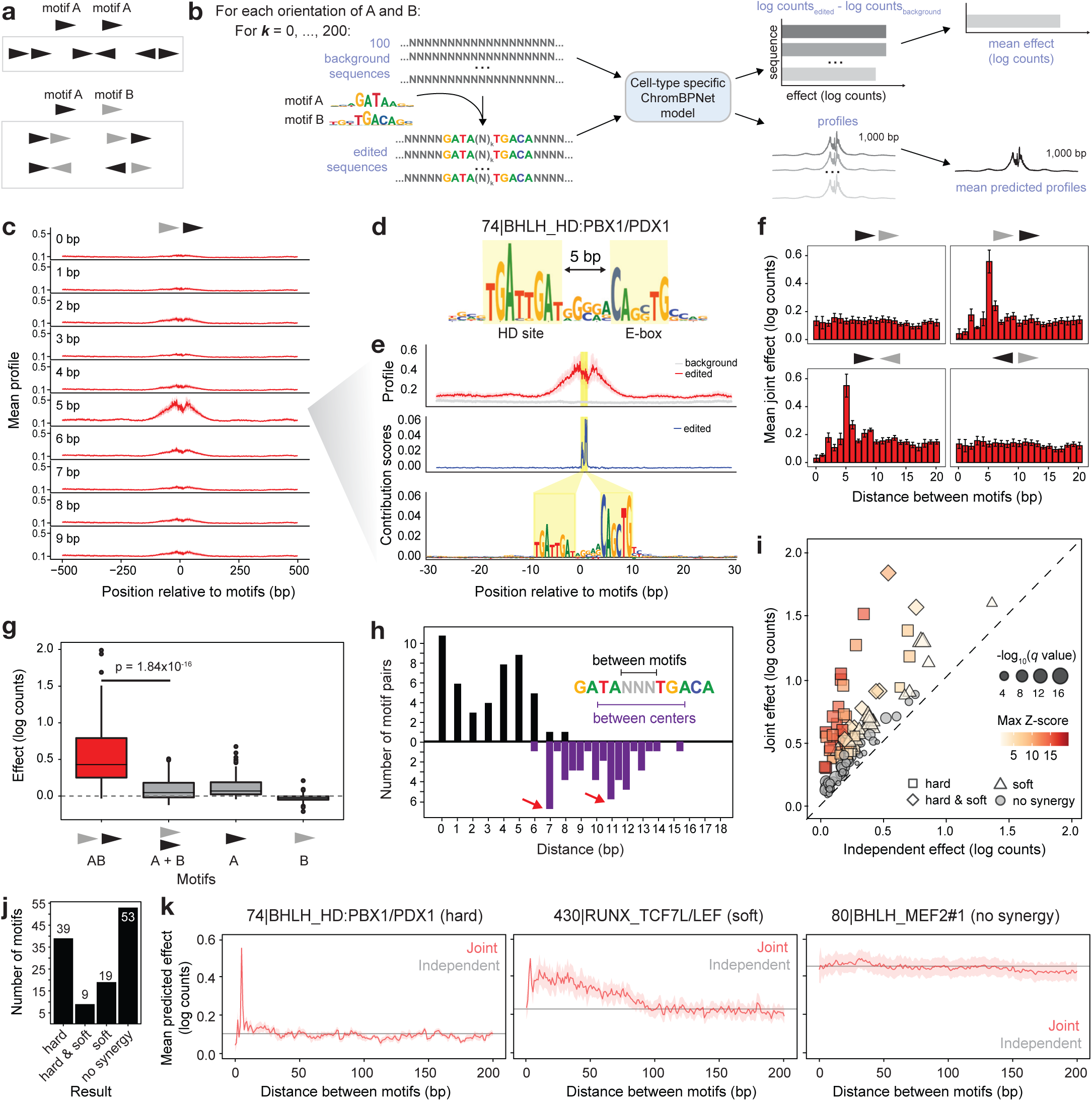
Systematic inference of TF cooperativity. **a)** Depiction of unique orientations of one or two distinct motifs. **b)** Workflow for *in silico* marginalization to predict causal effects of sequence motifs on accessibility. **c-g)** Synergy analysis for a composite motif with an HD site and BHLH site (E-box). **c)** Predicted profile when motifs are inserted at 0-9 bp distance, for one arrangement. **d)** CWM for the composite motif. **e)** Predicted profile at 5 bp, and mean contribution scores after model interpretation of edited sequences with motifs inserted. Yellow highlights indicated inserted nucleotides. **f)** Quantified predicted joint effects of two motifs in log counts, per orientation (error bars: standard deviation across folds). **g)** Distribution of predicted effects across 100 sequences, for both motifs in their optimal arrangement, the sum of motifs A and B inserted independently, and motifs A and B alone. P value: Wilcoxon signed-rank test. **h)** Distribution of distance between motifs (top) and between motifs centers (bottom) for all composite motifs inferred to have hard syntax preferences. **i)** Predicted joint effects (at optimal motif arrangement) and sum of independent effects for all motifs pairs tested. **j)** Barplot indicating number of composite motifs with hard, hard & soft, or soft synergy, or no predicted synergy. **k)** Predicted joint effects for motifs inserted at 0-200 bp distance between motifs, compared with sum of independent effects, for three representative composite motifs.

We recovered several known syntax-constrained composite elements. A *de novo* motif matching the recently described Coordinator element^50,51^—a composite of E-box and TAAT homeodomain motifs—showed strong synergy at a 5 bp head-to-tail arrangement (**Fig. S6a-d**). Remarkably, this arrangement precisely matched the experimentally validated binding geometry of the Coordinator complex, in which TWIST1–TCF4 heterodimers and ALX4 bind cooperatively. X-ray crystallography showed that this strict motif spacing facilitates stabilizing contacts between the TWIST1 backbone and an ALX4 side chain^50^, and the cooperative binding kinetics of these proteins were dependent on the strict 5 bp spacing between motifs. In our analysis, this motif was predominantly identified in skin cell types, consistent with its reported role in neural crest-derived mesenchyme. Model interpretation confirmed that the inserted motifs were responsible for driving predicted accessibility (**Fig. S6c**), and the joint effect significantly exceeded the sum of individual motif effects (*p* = 1.84×10⁻¹⁶, Wilcoxon signed-rank test) (**Fig. S6e**). We observed a similar syntax-constrained synergistic effect for a distinct composite motif composed of an E-box and an alternative homeodomain site (TGATTGAT), predominantly found in muscle and thymus cell types (**Fig. 4c-g**).

Across all 130 de novo composite motifs tested, we identified dozens with significant synergy, including 39 with hard syntax, 27 with soft syntax, and 9 exhibiting both (**Fig. 4h-j, Table S7**). Hard syntax motifs showed preferred spacings between motif centers around 7 bp and 11 bp— consistent with protein binding in adjacent or helical-turn offset DNA grooves—and exhibited sharp spikes in predicted joint effects at specific arrangements (**Fig. 4h,k, Fig. S6f-g**). In contrast, soft syntax motifs displayed broader distance preferences with more modest joint effects that decayed gradually beyond ∼100 bp (**Fig. S6f-g**). Outside of hard or soft synergistic motif arrangements, the joint effects of motifs were typically comparable to independent effects (**Fig. 4k, Fig S6f-g**). Among hard syntax motifs, we recovered composite elements consistent with known dimeric TF interactions, including SOX homodimers^52^, p53 homodimers^53,54^, and GATA– TAL heterodimers^55,56^ (**Fig. 5a**). Notably, the optimal spacing and orientation of the predicted p53 homocomposite motif matched the canonical 4 bp separation of its two NCATGN binding sites oriented head-to-tail^53,54^.

**Figure 5.**
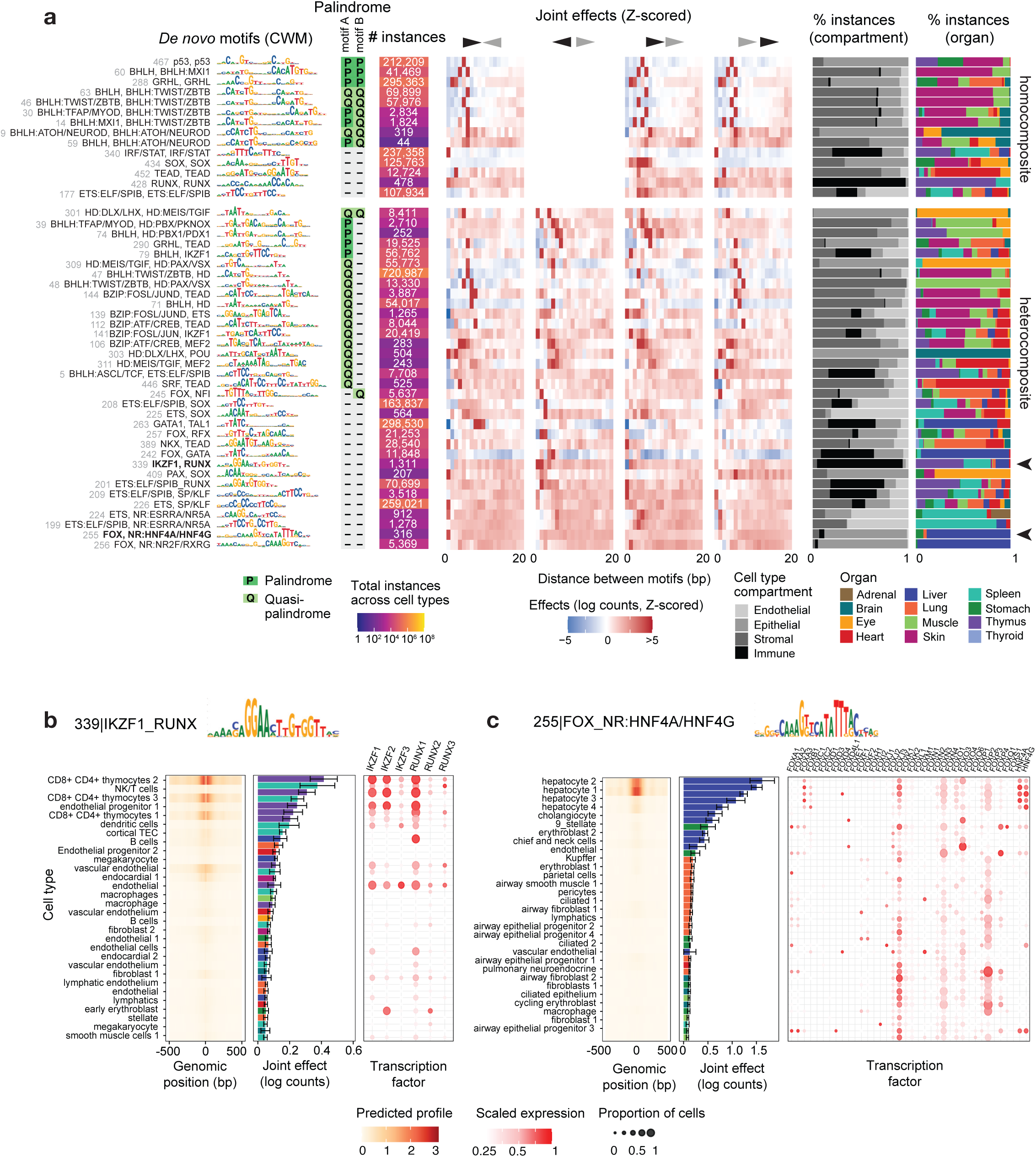
Composite motifs defined by hard syntax preferences. **a)** Summary of N=48 composite motifs inferred to be synergistic with hard syntax preferences. Left to right: *de novo* CWM-representation, annotation of palindromic sequences tested, total number of motif instances across cell types, predicted joint effects of inserted motifs at all arrangements at up to 20 bp (Z-scores were computed across all 200 bp), and distribution of motif instances across tissue compartments and organs. Arrowheads and boldface indicate motifs shown in (b) and (c). **b)** Predicted mean accessibility profile (left) and mean effect (middle) for *in silico* marginalization of IKZF-RUNX composite motif in each cell type, and aggregated expression of transcription factors which could bind constituent motifs in each cell type (right). Error bars: standard deviation across model folds. 40 cell types with the highest predicted accessibility are shown. **c)** Same as (b), for the FOX-HNF4 composite motif.

Synergistic motif syntax was detected across most cell lineages. Approximately 60% of cell types revealed at least one hard syntax and one soft syntax composite motif pattern. Within each cell type, hard syntax motifs were significantly more prevalent, based on the number of predictive instances, than motifs with soft syntax or no synergy (**Fig. S6h**). Hard and soft syntax composite motifs also exhibited higher total contribution scores (across positions of the motif CWM) compared to motifs without synergy and non-composite motifs, indicating greater predictive influence on chromatin accessibility (**Fig. S6h**).

Several synergistic composites with constrained syntax involved cell type-specific motifs, prompting us to ask whether their cooperative effects were also cell type-specific (**Fig. 5a**). To test this, we inserted motif pairs at their optimal arrangement into background sequences and predicted accessibility across all cell types using their respective ChromBPNet models. For example, an IKZF–RUNX composite motif drove accessibility only in a subset of thymus- and spleen-derived immune cell types (**Fig. 5b**). This pattern was consistent with the expression of IKZF and RUNX family TFs in those same cells, suggesting that cooperative effects are constrained not only by motif syntax but also by the availability of constituent factors. Similarly, a FOX–HNF4 composite motif exhibited predicted accessibility specifically in hepatocytes (**Fig. 5c**), correlating with restricted expression of FOX and HNF4 family members such as *FOXA3* and *FOXB1* in these cells (**Fig. 5c**). Although limited detection of lowly expressed TFs in single-cell RNA-seq precludes comprehensive analysis, these examples support a model in which motif cooperativity depends jointly on precise binding site syntax and cell type-specific TF expression.

These findings reveal that a substantial fraction of motifs exhibit cooperative effects on chromatin accessibility governed by distinct syntactic constraints. Hard syntax motifs suggest direct protein– protein interactions at fixed geometries, while soft syntax motifs likely reflect indirect cooperativity mediated by nucleosomes or chromatin remodelers. These *de novo* predictions from the models nominate candidate TF pairs and motif syntaxes for further mechanistic investigation in developmental gene regulation.

### Ubiquitous motifs associated with reduced accessibility and concordant eQTL effects

While the majority of *de novo* motifs had positive contribution scores—suggesting a role in promoting chromatin accessibility—a small subset (15 motifs; 3% of the lexicon) were predicted to have negative effects (**Fig. 6a-b**). Despite being a minority in the total set of motifs, these negative motifs were widespread in accessible genomic regions, with 2–5 predictive instances per peak and comprising over one-third of all predictive motif instances in several organs (**Fig. S4e, Fig. 6b**). Many matched known TF families with repressive activity, including ZEB/SNAIL, HIC, BCL11A, NFY, and YY1/2, although their ubiquitous negative effects on chromatin accessibility have not been previously reported (**Fig. 6c**). Other negative motifs were unresolved and did not match known databases (**Fig. S5b**).

**Figure 6.**
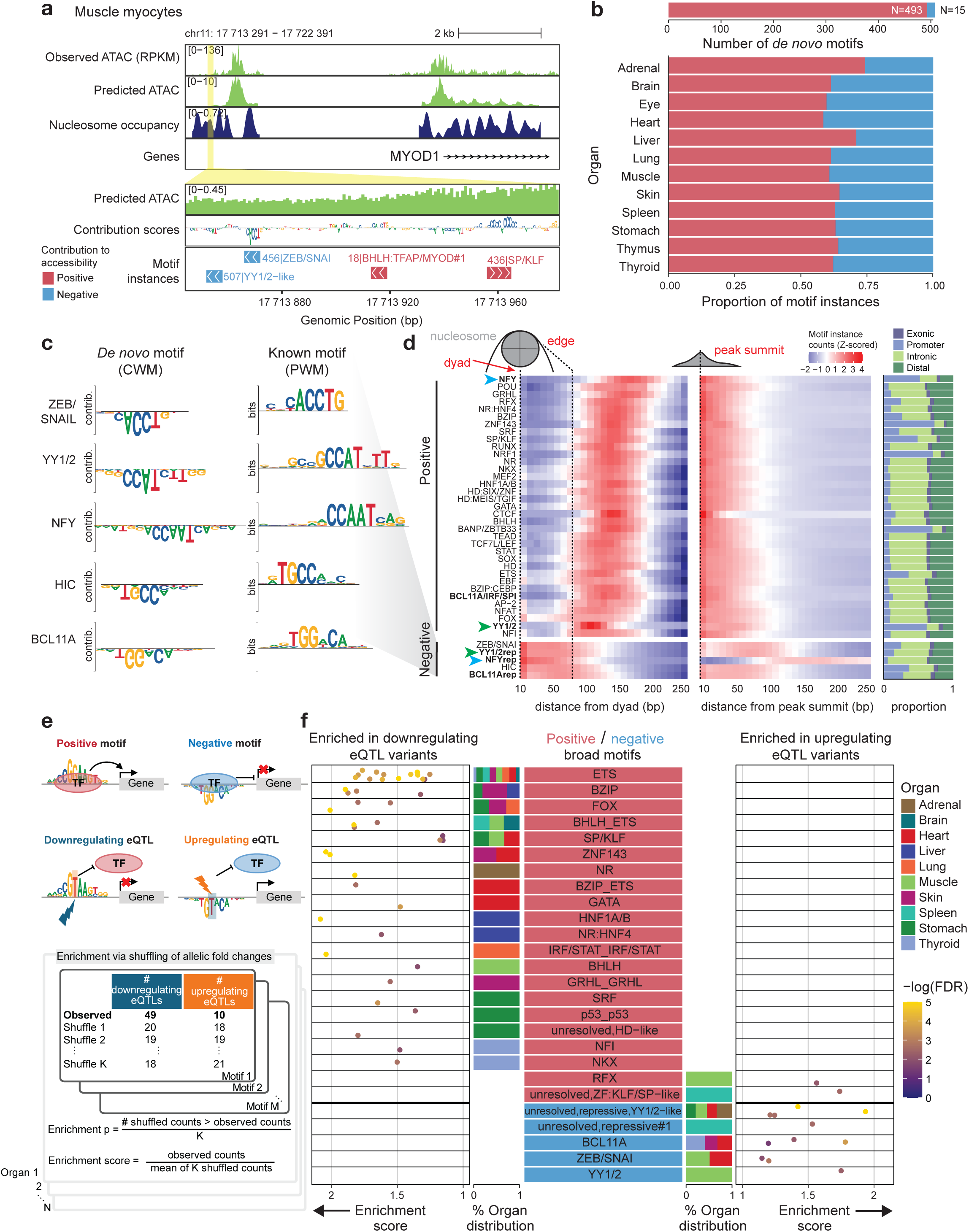
Ubiquitous motifs associated with reduced accessibility and directional eQTL effects. **a)** *MYOD1* locus in myocytes with observed and predicted chromatin accessibility, inferred nucleosome occupancy, contribution scores, and motif instance annotations showing an example of a negatively-contributing ZEB/SNAIL motif. **b)** Breakdown of *de novo* motifs, and breakdown of motif instances for each organ, by positive and negative motifs. **c)** *De novo* motifs (as CWMs) for each negative motif category, and most similar known PWM in external databases**. d)** Left and middle: Heatmaps indicating counts of motif instances in 10 bp bins from inferred nucleosome dyad positions (left) and peak summits (middle), Z-scored across distance bins per motif in each heatmap. Right: Proportion of genomic instances overlapping various genomic features. Each row represents a broad group of base motifs, and only groups with at least 50,000 instances across bins are shown. **e)** Workflow for determining tissue-specific enrichment of different classes of fetal motifs with upregulating or downregulating eQTL variants. **f)** Results of eQTL variant analysis. For each broad motif category, points indicate enrichment scores of each unique motif from each organ, colored by -log_10_ FDR values for downregulating eQTL variants (left) and upregulating eQTL variants (right). Stacked bar plots display the proportion of organs in which motif instances with enrichment originate. Only statistically significant enrichments are shown.

Negative motifs exhibited distinct positional preferences in peaks compared to positive motifs. They were enriched near nucleosome dyads and depleted near peak summits, in contrast to activating motifs, which tended to avoid dyads and cluster near summits (**Fig. 6c-d**). Intriguingly, NFY and YY1/2 motifs were discovered to exhibit both positive and negative effects on accessibility with distinct positional preferences, with separate positive and negative motifs discovered for each factor (**Fig. 6c**). While positive YY1/2 motifs were enriched adjacent to the nucleosomes, consistent with previous observations^4^, their negative counterparts were enriched near inferred dyads. For NFY, positive motifs were among the most nucleosome-avoidant, and concentrated near peak summits, but negative motifs were enriched in peak flanks, at 100-200 bp from peak summits (**Fig. 6d, Fig. S4i**).

To assess whether positive and negative motifs also differ in their effects on downstream gene expression, we analyzed the enrichment of directional effects of fine mapped variants in expression quantitative trait loci (eQTLs) from the Genotype-Tissue Expression (GTEx) dataset^57^ that overlapped predictive instances of positive and negative motifs from related fetal organs (**Fig. 6e, Methods**). We reasoned that variants disrupting positive motifs should predominantly reduce gene expression, whereas variants disrupting negative motifs should increase expression. For each fetal organ, we quantified enrichment per motif of upregulating and downregulating eQTL gene-variant pairs from the relevant GTEx tissues, compared to a background set derived by shuffling effects. Although only a small fraction of eQTL variants overlapped our predictive motif instances (**Fig. S7**), we observed enrichments generally consistent with our hypothesis. Across multiple organs, eQTLs overlapping positive motifs were significantly enriched for downregulating variants (Fisher’s exact test, *p* = 9.91×10⁻¹¹, OR = 0.047), while eQTLs overlapping negative motifs were enriched for upregulating eQTL variants (*p* = 8.01×10⁻⁴, OR = ∞) (**Fig. 6f, Table S8**).

Notably, we found tissue-specific enrichments of eQTL variants in instances of motifs known to have important tissue-specific roles in development, such as GATA motifs in heart^58^, HNF4 in liver^59^, and NKX motifs in thyroid^60,61^ (**Fig. 6f**).

These results show that deep learning models distinguish between noncoding DNA regions with identical sequence features but distinct chromatin localization and opposite contributions to accessibility. Positive and negative motifs are differentially enriched for variants that decrease or increase gene expression, respectively, highlighting their distinct regulatory roles. These findings suggest that the balance of motifs with opposing effects is potentially important to fine-tune chromatin accessibility and gene regulation during development.

### Accessible regions in fetal cell types are enriched with genetic variants associated with disease

Thousands of genome-wide association studies (GWAS) performed on myriad complex traits and diseases have linked genomic loci to complex phenotypes^62^, but it remains challenging to pinpoint causal variants and the relevant cell types they impact. We thus asked whether accessible chromatin landscapes in fetal cell types are enriched for genetic variants associated with human disease (**Fig. 7a**). We intersected candidate cis-regulatory elements from HDMA with fine-mapped variants from CAUSALdb^63^, which aggregates credible sets across 1,483 genome-wide association studies spanning 349 distinct traits.

**Figure 7.**
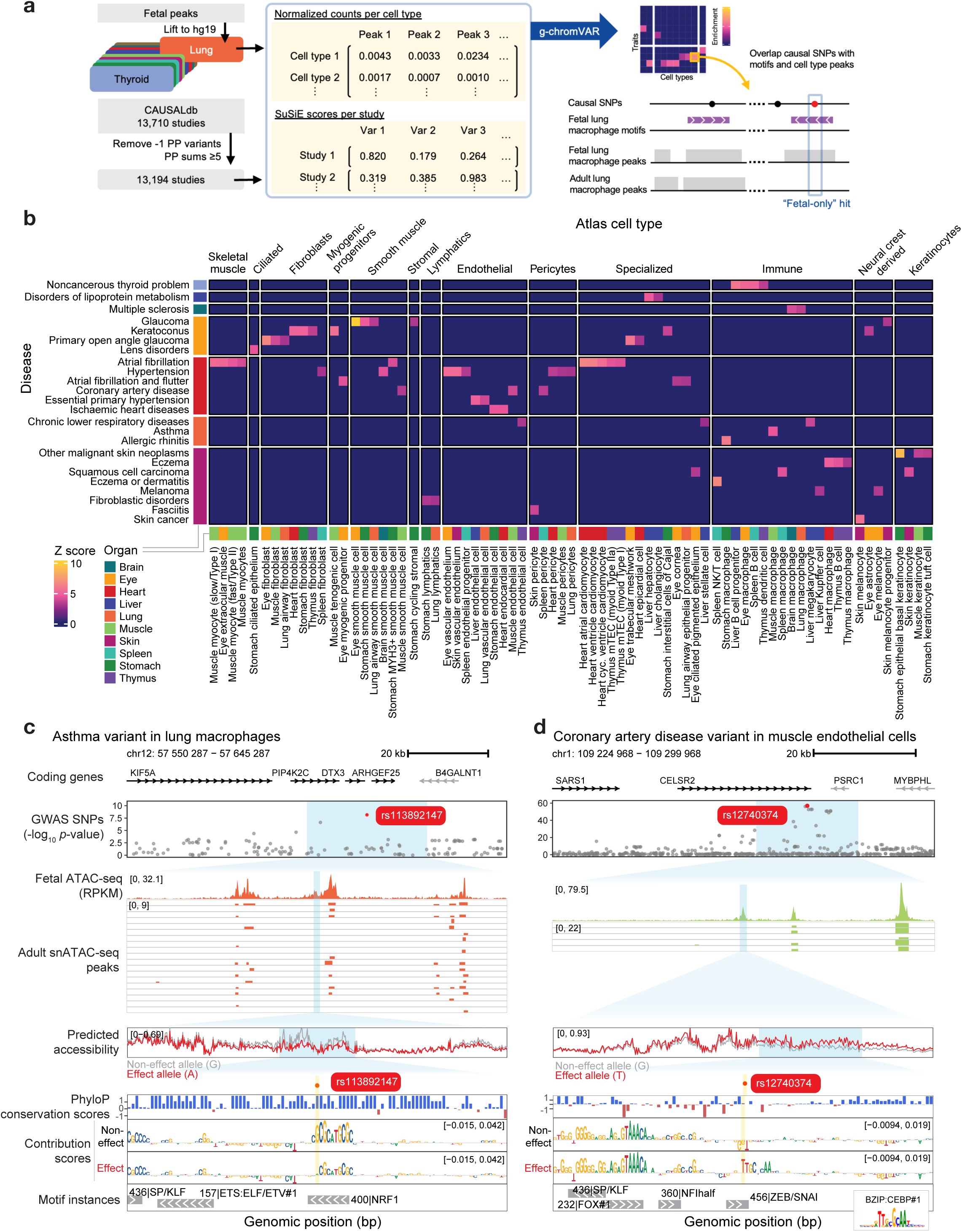
Disease causal variants overlap motifs in fetal-only peaks. **a)** Workflow for the identification of disease-relevant causal variants in motifs that are found in fetal but not adult peak sets. **b)** Top hits from g-chromVAR enrichment results showing only disease traits with the highest average Z-scores among similar MeSH terms, and L2 fetal cell types with the highest total Z-scores further grouped by L3 cell type annotations. **c)** rs113892147, an asthma variant, is a fetal-only hit in a positive NRF1 motif in fetal lung macrophages, predicted to reduce accessibility. From top to bottom, tracks are the coding genes, p-values for GWAS SNPs, observed pseudobulk accessibility for fetal lung macrophage cluster, peaks from snATAC-seq in adult lung macrophages, predicted accessibility for the non-effect and effect alleles, PhyloP conservation scores, per-basepair scores for contribution to predicted accessibility for non-effect and effect alleles, and the motif instances for the non-effect allele. **d)** rs12740374, a coronary artery disease variant, is a fetal-only hit overlapping a ZEB/SNAIL negative motif in muscle endothelial cells and predicted to increase accessibility through creation of a C/EBP site (the corresponding motif is shown as an inset at bottom right). Tracks are as in (c).

Using g-chromVAR^64^, we quantified the enrichment of trait-associated variants across all fetal L2 cell types. This analysis identified 131 cell types with significant enrichment (FDR < 0.05) for credible set variants (**Table S9**). To focus on disease-relevant signals, we curated a subset of 68 cell types enriched for variants from 79 studies across 36 MeSH disease terms (**Table S10**).

We observed that variant enrichments were predominantly cell type-specific but organ-agnostic: similar cell types from different organs tended to be enriched for the same disease traits (**Fig. 7b**). For example, immune cell types were enriched for variants associated with thyroid disease^65,66^, dermatitis and eczema^67,68^, rhinitis^69,70^, and cancer^71^. Myocytes were linked to atrial fibrillation risk^72,73^, while endothelial cells were enriched for variants associated with hypertensive diseases^74–76^. Specialized liver parenchymal cells, such as hepatocytes and cholangiocytes, were enriched for variants linked to lipid metabolism disorders^77^.

Overall, these results suggest that cis-regulatory elements of developmental cell types may harbor genetic variants that may predispose individuals to a wide range of adult-onset diseases.

### Interpreting effects of putative causal variants on chromatin accessibility in fetal regulatory elements

To gain mechanistic insight into how disease-associated genetic variants might perturb regulatory activity during development, we prioritized putative causal variants overlapping fetal-specific accessible elements. From disease-enriched fetal cell types, we identified high-confidence causal variants (SuSiE^78^ posterior inclusion probability (PIP) ≥ 0.8) overlapping predictive motif instances (**Table S11**). To distinguish fetal-specific effects from shared effects with adult cell types, we compared variant overlaps with HDMA acCREs to overlaps with ATAC-seq peaks from matched adult cell types from ENCODE^79^ (**Fig. 7a, Table S12**). We identified 28 hits as fetal-only and 80 hits present in both fetal and adult peak sets. The remaining 74 hits did not have matching adult cell types.

Using our trained cell type-specific ChromBPNet models, we predicted the effect of each of the 28 fetal-specific variants on local chromatin accessibility and computed sequence contribution scores in the relevant cell types (**Table S13**). Many fetal-only variants were predicted to either reduce accessibility by ablating positive motifs or increase accessibility by disrupting negative motifs (**Fig. S8**). We highlight two such fetal-specific variants (**Fig. 7c-d**) which involve a positive NRF1 motif (Z-score = 3.27, FDR = 2.09×10^-2^) associated with asthma^80^ in lung macrophages, and a negative ZEB/SNAIL motif (Z-score = 3.45, FDR = 1.42×10^-2^) found in muscle endothelial cells associated with coronary artery disease^81^ (CAD).

The asthma-associated fine mapped variant (rs113892147, SuSiE PIP = 0.819) overlapped a conserved NRF1 binding motif^82^ within an accessible region in fetal lung macrophages (**Fig. 7c**). This site lacked an accessibility peak in all 20 adult lung macrophage samples, suggesting fetal-specific regulatory activity. Installation of the effect A allele was predicted to reduce accessibility relative to the reference G allele by the fetal lung macrophage ChromBPNet model, consistent with eQTL data showing reduced expression of the nearby gene ARHGEF25 in adult lung tissue (beta = -0.295, p = 6.9×10^-7^, Open Targets Platform^83^). Although the direct connection to asthma pathology remains unclear, this result highlights how fetal macrophage enhancers may shape genetic risk factors of asthma.

The CAD-associated variant (rs12740374, SuSiE PIP score = 0.855) overlapped a negative ZEB/SNAIL motif in fetal muscle endothelial cells (**Fig. 7d**). Installation of the effect G allele disrupted the weak negative ZEB/SNAIL motif and created a C/EBP binding site relative to the reference T allele, leading to a predicted increase in accessibility from the corresponding ChromBPNet model. This locus (CELSR2–PSRC1–SORT1) is well-established in CAD risk^84–87^, and prior studies have shown that in hepatocytes, this variant enhances C/EBP binding and SORT1 expression^87,88^. Our findings extend its potential regulatory role to endothelial cells, consistent with emerging models implicating vascular dysfunction in CAD pathogenesis. Interestingly, outside of the liver, rs12740374 is associated with increased expression of the nearby *CELSR2* (beta = 0.338, *p* = 1.2×10^-36^) and *PSRC1* (beta = 0.238, *p* = 4.6×10^-8^) genes in muscle but not in heart ventricle or atrial tissues (Open Targets Platform), further highlighting the potential roles for non-cardiac cells in CAD.

Overall, these results demonstrate that deep learning models trained on fetal regulatory landscapes can predict the consequences of disease-associated variants on cell-type specific chromatin accessibility, revealing how noncoding variation potentially perturbs gene regulation and contributes to disease susceptibility later in life.

## Discussion

Here, we defined the chromatin accessibility and transcriptomic landscapes of 203 primary human fetal cell types across 12 organs, creating a comprehensive multiomic atlas of human development. We mapped over one million accessible regulatory elements and demonstrated their ability to resolve organ- and cell type-specific enhancer activity *in vivo*. By training deep learning models to predict accessibility directly from DNA sequence, we identified a predictive lexicon of regulatory motifs, uncovered distinct modes of TF cooperativity governed by motif syntax, and systematically interpreted the impact of disease-associated non-coding genetic variants on chromatin landscapes in fetal cell types. Together, these findings reveal how DNA sequence encodes the cis-regulatory syntax that drives cellular diversity during development and suggest fetal cell types and TFs that might influence genetic risk of adult on-set diseases.

Compared to previous efforts, our work substantially extends the resolution and scope of regulatory mapping in human development. Prior TF binding and chromatin profiling experiments in cell lines and bulk tissue samples^5,6,25,89^, single organs^7,8^, and single modality fetal atlases^9,10^ have provided valuable resources, but could not resolve cell type-specific regulatory logic across multiple organs. Our atlas includes over one million regulatory elements active in primary fetal cell types, 14% of which were not captured in previous datasets. Novel elements were particularly enriched in brain and eye cell types, likely reflecting the increased resolution afforded by single-cell profiling. Our data also refined the annotation of experimentally validated VISTA enhancers by resolving ambiguous organ activity. We found that tissue-specific motifs were primarily located in distal and intronic elements, while promoter-dominant motifs included ubiquitous regulators^90^ and CpG-rich, methylation-sensitive TFs such as NRF1^42^ and BANP^91^. These observations are consistent with prior findings that a limited set of core motifs regulate the majority of transcriptional initiation events^44–46,90,92^.

Our deep learning framework is complementary to classical motif discovery approaches such as motif enrichment from DNase footprinting^5,93^ or *in vitro* TF-DNA binding assays^11,12,94^. First, unlike footprinting methods that require deep sequencing, ChromBPNet models can be trained and interpreted on pseudobulk data from single-cell clusters with modest coverage, enabling robust, cell context-specific motif discovery across a broad range of primary tissues (**Note S2**). Further, unlike motif enrichment-based methods, our models learn likely causal sequence features directly predictive of chromatin accessibility, allowing us to distinguish regulatory motifs that actively shape chromatin landscapes^17^. Moreover, deep learning models offer a unique capability of predicting the impact of *in silico* perturbations of DNA sequence, enabling systematic interrogation of motif synergy, spacing, and orientation. This approach not only enables recovery of known motifs, but also empowers discovery of novel motifs, motif variants, and higher-order syntax, thereby providing mechanistic insight into the regulatory syntax encoded in regulatory DNA.

We discovered 67 motif pairs exhibiting synergistic effects on chromatin accessibility, with 48 showing strong preference for specific spacing and orientation—indicative of “hard” motif syntax. This mode of cooperativity recalls the classic “enhanceosome” model^95^, exemplified by the IFN-β enhancer^96^, in which precise spatial organization of TF binding sites is essential for assembly and function of the TF complex. Additional examples from immune regulation, including the AP-1– IRF4 (“AICE”) and ETS–IRF (“EICE”) composite elements, also demonstrate rigid binding architectures that promote lineage-specific gene activation during T and B cell differentiation^97,98^. Prior systematic surveys of TF cooperativity using *in vitro* systems have identified similar spacing and orientation preferences^11,12^, but lacked the *in vivo* resolution needed to assess genome-wide regulatory syntax during human development. Notably, those studies often focused on composite motifs with fused and overlapping constituent binding sites, which we did not specifically prioritize in our *in silico* experiments. Furthermore, because our motif lexicon is built from clustered *de novo* motifs, additional diversity—including fused, partially overlapping or low affinity non-canonical configurations—may emerge from finer-grained dissection of motif subclusters^18^. The widespread presence of predicted synergistic, tissue-specific composite motifs in our analysis supports the view that densely packed, spatially constrained TF interactions are a common regulatory mechanism underlying developmental gene programs.

In parallel, we identified 27 synergistic motif pairs exhibiting “soft” syntax, in which synergy persisted across a broader range of distance between motifs (typically <150 bp). These effects are consistent with indirect cooperativity models, including mass-action models of TF competition with nucleosomes, or recruitment of chromatin remodelers which evict nucleosomes from chromatin^13,14^. Such syntactic flexibility may confer greater evolutionary robustness, allowing regulatory elements to maintain function while tolerating mutations in motif spacing or orientation. These findings suggest that both rigid and flexible modes of motif cooperativity are employed across human development, expanding the motif syntax which mediates the contribution of TFs to chromatin landscapes.

Interpretation of deep learning models is a powerful technique that enables direct interrogation of the quantitative, directional effects of sequence features on chromatin accessibility, thereby identifying sequence motifs that negatively or positively influence accessibility. Although negative motifs represented a small fraction of the *de novo* motifs recovered, they were highly abundant across the genome, with one negative motif (matching ZEB/SNAIL recognition sites) emerging as the most frequent predictive motif instance. These negative motifs exhibited distinct positional biases—enriched near nucleosome dyads and distal regions rather than peak summits—and were significantly enriched for eQTL variants associated with increased gene expression when disrupted. Several lines of evidence independently support the regulatory role of negative motifs: in *Drosophila*, a ZEB2 repressor motif—similar to our ZEB/SNAIL motif—was shown to decrease enhancer activity^99^; in a neurodevelopmental disorder case, a *de novo* variant creating a repressive NR2F1 motif significantly reduced enhancer-driven reporter expression in mouse brain ^17^; and in the developing liver, abundant regions enriched for repressive motifs such as ZEB1, TCF4, and SNAI1 were predictive of lower enhancer activity and gene expression and suggested to finetune regulation^100^. In our dataset, negative motifs included known repressors and previously uncharacterized sequence elements, suggesting a broader repertoire of short repressive signals modulating accessibility in human development.

Interestingly, the models also revealed that certain TF motifs, such as YY1/2 and NFY, could function as either positive or negative regulators depending on their genomic context. For example, negative YY1/2 and NFY motifs were preferentially enriched at distal regions and near nucleosome dyads, in contrast to their positive variants near promoters. This is consistent with the known dual functions of YY1 as both an activator and repressor depending on context^101^ and the role of NFY as a histone-fold protein that maintains nucleosome-depleted regions at active regulatory elements^102,103^.

Although we uncover widespread evidence for short repressive motifs, they are distinct from classical silencer elements^104^, and their precise mechanisms of action remain to be fully elucidated. Future work will be needed on the collection of negative motifs to identify the proteins binding these motifs, define their interactions with chromatin remodelers or nucleosomes, and understand how repression is contextually integrated with activating signals to modulate accessibility and gene expression during human development.

Our eQTL analysis further highlighted that disruption of positive motifs typically decreases gene expression, consistent with a loss of activating inputs. However, two positive motifs—an RFX motif and an unresolved motif resembling a KLF/SP site—showed the unexpected enrichment of upregulating eQTLs. This could reflect variants that improve motif affinity, leading to enhanced TF binding and increased target gene expression, a phenomenon observed for gain-of-function mutations impacting limb and heart development^105,106^. Alternatively, TFs may increase chromatin accessibility while repressing gene expression, depending on the regulatory context. Indeed, RFX family members regulate distinct gene sets across tissues^107^ and can act as repressors when bound to specific regulatory elements^107,108^. REST provides a well-established example of a TF that binds neuron-restrictive silencer elements, and is positively associated with local accessibility (**Fig. 3e**), but functions as a transcriptional repressor^109^. These findings illustrate the power of deep learning models to decouple the influence of DNA-binding proteins on chromatin accessibility from their downstream effects on gene regulation.

Finally, by intersecting fetal regulatory maps with fine-mapped genetic association studies, we linked disease-associated variants to specific developmental cell types and predicted their effects on chromatin accessibility. Many of these variants were located in fetal-specific accessible regions absent from adult tissues. These predictions nominate specific regulatory elements, transcription factors, and cell types for mechanistic investigation across a wide range of conditions, from asthma to coronary artery disease. The identification of causal variants associated with adult-onset diseases within fetal-specific regulatory elements raises fundamental questions about how genetic risk is encoded during development. For example, our finding that asthma-associated variants affect fetal macrophage enhancers suggests that early-acting variants could compromise the development of embryonically derived, long-lived cell types. Alveolar macrophages, which originate from fetal liver monocytes, establish residence in the lung before birth and persist into adulthood as self-renewing, tissue-resident cells^110–113^. Thus, genetic variants may exert lasting effects by perturbing the developmental specification of disease-relevant cell lineages, even if the affected chromatin elements become inaccessible in adult tissues. Alternatively, some variants may increase disease predisposition by enabling aberrant reactivation of fetal regulatory elements later in life, suggesting a latent regulatory potential encoded in early development. These findings underscore the importance of studying gene regulation in a developmental context to fully understand the molecular origins of human disease.

Our study has several limitations. Although our atlas captures a broad range of fetal organs and cell types, deeper sampling will be needed to fully resolve the diversity of cell types in certain tissues, rare cell populations, and cell types that arise in later developmental stages. Our data does not recapitulate organ-specific enhancer activity at all experimentally validated VISTA enhancers, perhaps because these enhancers were only assayed at mouse embryonic day E11.5, which is transcriptionally most similar to human post-conception week 5^114^, substantially earlier than the samples we profiled. Deep learning models trained on chromatin accessibility primarily capture the influence of sequence-specific DNA-binding proteins that modulate accessibility and may miss regulators that act downstream or independently of accessibility^17^. Furthermore, motif degeneracy within TF families precludes definitive assignment of individual TFs to *de novo* predictive motifs without orthogonal data. Finally, while our variant effect predictions offer mechanistic hypotheses, experimental validation in the same cell contexts will be required to confirm the predicted impacts on chromatin state and gene regulation.

In conclusion, we present a comprehensive, multiomic single-cell atlas of human fetal development together with a predictive framework for decoding the cis-regulatory syntax that governs chromatin accessibility across cell types. By identifying predictive sequence motifs, syntactic rules of motif cooperativity, and variants that perturb regulatory logic, we uncover how non-coding DNA encodes cell type-specific regulatory programs and how their disruption contributes to disease risk. To support future investigations, we provide an extensive resource of processed single-cell data, accessible chromatin elements, nucleosome occupancy profiles, contribution scores, predictive motif instances, and cell type-specific deep learning models. These datasets and models—accompanied by genome browser compatible tracks and an efficient preprocessing pipeline—enable detailed exploration of regulatory mechanisms across human development and offer a foundation for studying the genetic and molecular basis of disease in a developmental context.

## Acknowledgements

We thank Zhainib A. Amir-Ugokwe and Kristy Red-Horse for their advice on embryo embedding and sectioning, Jackie Tang for advice on histology staining, Steve Avolicino (Histo-Tec Laboratories) for optimizing paraffin embedding protocols, Nick Addleman, John Coller and the Stanford Genome Technology Center for their help with sequencing libraries, Soon Il Higashino for keeping our laboratory running, Anna Scherbina for maintaining our lab compute cluster, and members of the Greenleaf and Kundaje labs as well as Chew Guo-Liang for helpful discussions. VISTA enhancer work was supported by a U.S. National Institutes of Health (NIH) grant to L.A.P. (R01HG003988) and conducted at the E.O. Lawrence Berkeley National Laboratory and performed under U.S. Department of Energy Contract DE-AC02-05CH11231, University of California. This work was supported by a Wu Tsai Neurosciences Institute Postdoctoral Fellowship (S.J.), Stanford Graduate Fellowship (B.L.), NSERC Postgraduate Scholarship (B.L.), and funding from the Singapore Economic Development Board Industrial Postgraduate Programme (Y.T.N.). W.J.G Acknowledges support from the Arc Institute and the Chan-Zuckerberg Biohub.

## Author Contributions

B.B.L., S.J., S.H.K., and Y.T.N. contributed equally as co-first authors, and have agreed that any author can be listed first in reporting this study. S.H.K. and W.J.G. conceived of the study. S.H.K., S.H., L. L. and A.M.P. performed tissue collection. S.H.K. optimized the SHARE-seq protocol. B.B.L., S.H.K., S.H., G.K.M., D.C.C. and T.C.N. performed SHARE-seq experiments. B.B.L. and B.E.P. wrote the snakemake preprocessing pipeline with input from G.K.M.. B.B.L., S.H.K. and S.J. programmed and performed data preprocessing. B.B.L., S.H.K., S.J. and S.K.W. annotated all cell types. B.B.L. performed ABC and VISTA analysis. L.A.P. and M.K. provided the VISTA embryos and contributed to VISTA analysis discussions. S.T. helped annotate mouse embryo sections. S.J. trained ChromBPNet models, and performed analyses related to the motif lexicon and TF cooperativity with input from A.K.. A.T.W. developed the algorithm to call predictive motif instances with input from A.K.. Y.T.N. performed eQTL and causal variant analysis with input from E.B.D.. B.B.L., S.J., S.H.K., Y.T.N., A.K. and W.J.G. wrote the manuscript, with input from all authors. A.K., K.K.H.F. and W.J.G. jointly supervised the work.

## Supplementary Figures

**Figure S1, related to Figure 1.**
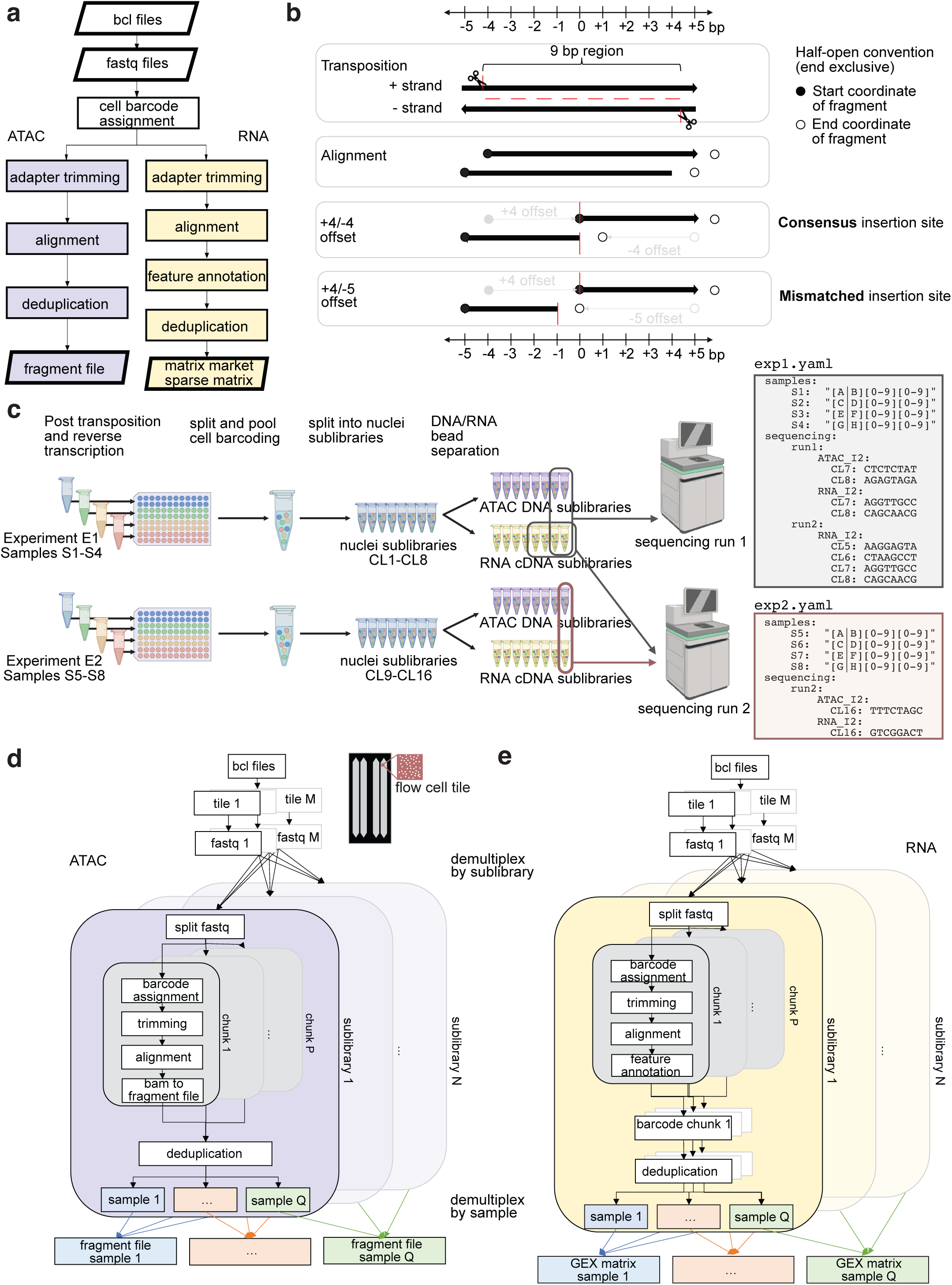
A highly parallelized, rapid and storage-efficient pre-processing snakemake pipeline to convert BCL files to ATAC fragment files and RNA sparse matrices. **a)** Overview of pre-processing workflow. Files are bolded in parallelograms while processes are in rectangles. **b)** Schematic explaining the +4/-4 offset adopted by this pipeline to reach a consensus Tn5 insertion site. **c)** Definition of a SHARE-seq experiment, samples, nuclei sublibraries, ATAC and RNA sublibraries. Example snakemake configuration file snippets to process sequencing data from mixed experiments. Each experiment should be run in a separate snakemake directory. Detailed pre-processing workflow showing the parallelization behind the scenes for **d)** ATAC and e) RNA sequencing data.

**Figure S2, related to Figure 1.**
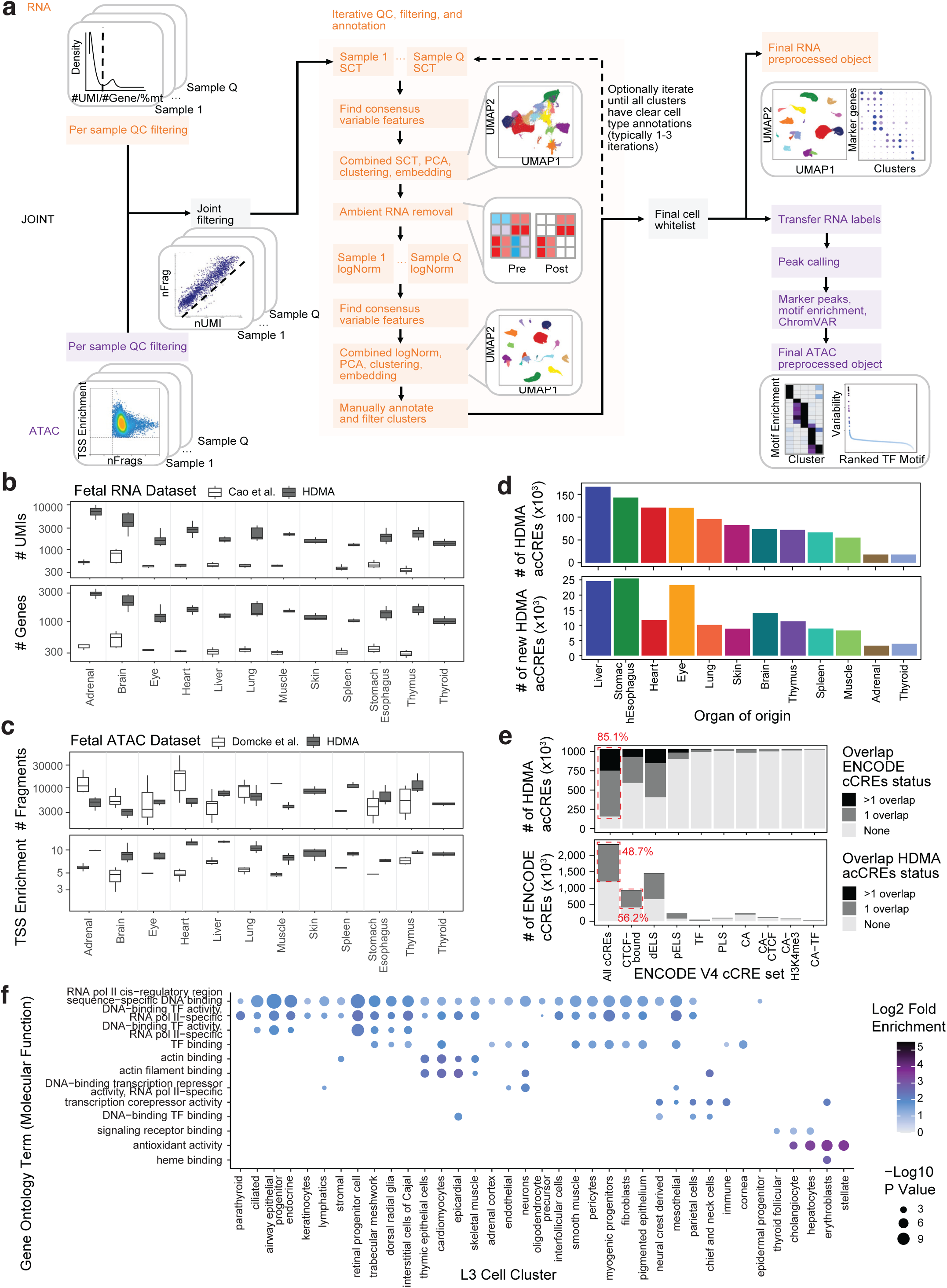
Cell type annotation workflow and regulatory elements landscape overview. **a)** Overview of the iterative cell type annotation pipeline converting fragment files to annotated ArchR project for ATAC data and sparse matrices to annotated Seurat project for RNA data. **b)** Comparison of the number of UMIs and number of genes detected per cell in RNA modality with Cao et al and **c)** comparison of the number of fragments and TSS enrichment ratio per cell in ATAC modality with Domcke et al. **d)** Organ origin of all HDMA acCREs and *de novo* HDMA acCREs. **e)** Overlap between HDMA acCREs and all or subsets of ENCODE v4 cCREs. dELS=distal enhancer-like signature. pELS=proximal enhancer-like signature. PLS=promoter-like signature. CA=chromatin accessible. **f)** Top Gene Ontology molecular function terms enriched in the highly regulated genes (HRG) in each broad class cell cluster (L3 cell cluster).

**Figure S3, related to Figure 2.**
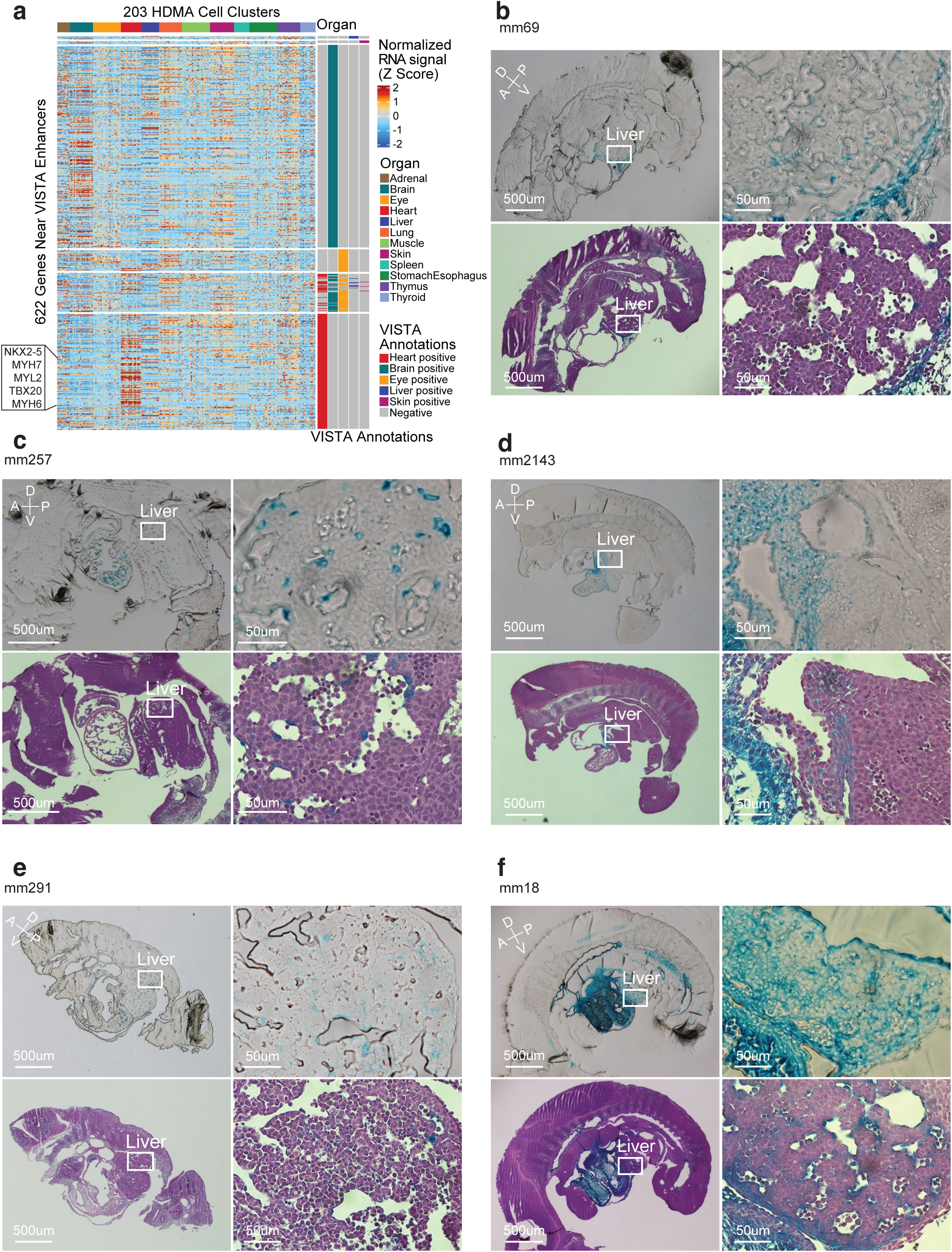
VISTA enhancer gene expression and organ specificity validation. **a)** Normalized and Z-scored expression of the nearest gene in HDMA for each VISTA enhancer, grouped by VISTA-annotated organ positivity. **b-f)** Bright field (top) and H&E images (bottom) for 5 VISTA enhancers re-annotated as positive in the liver based on HDMA data. Blue color is from X-Gal staining indicating where the enhancer is active. A=anterior, P=posterior, D=dorsal, V=ventral.

**Figure S4, related to Figure 3.**
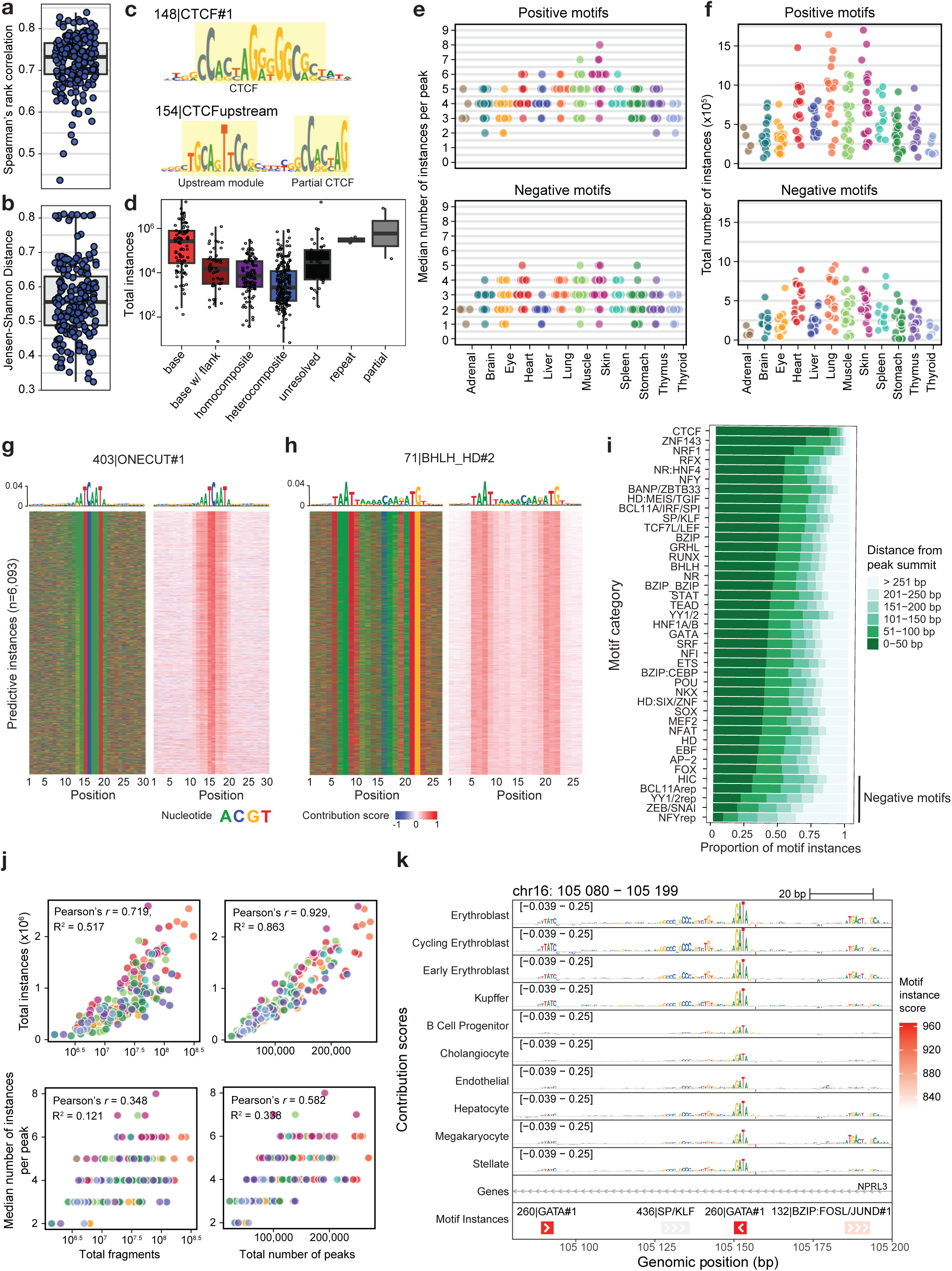
Modeling of sequence determinants of chromatin accessibility. **a)** Distribution of Spearman correlation between predicted and observed log counts in peak regions per model (mean across five folds). **b)** Distribution of Jensen-Shannon Distance between predicted and observed accessibility profiles in peak regions per model (mean across five folds). **c)** Example contribution weight matrix (CWM, neural network-derived motif representation) for distinct CTCF upstream and CTCF motifs. **d)** Total number of motif instances across all cell types for the 508 lexicon motifs, grouped by motif category. Each point is one motif. **e)** Median number of motif instances per peak per cell type, for all peaks with instances called. **f)** Total number of motif instances per cell type. **g)** Illustrative representations of all motif instances for ONECUT motif in liver hepatocytes. Heatmaps indicating nucleotide composition (left) and contribution scores (right) for 30 bp centered at all 6,093 motif instances. **h)** Same as (g), for a BHLH, HD composite motif. **i)** Proportion of total motif instances across cell types at each distance from peak summits, for each motif category, sorted by proportion of instances within 50 bp of peak summits. **j)** Relationship between total number of motif instances and median positive motif instances, and total number of fragments or peaks for a given cluster. **k)** Contribution scores of GATA motif instances within mm101 enhancer in liver cell types.

**Figure S5, related to Figure 3.**
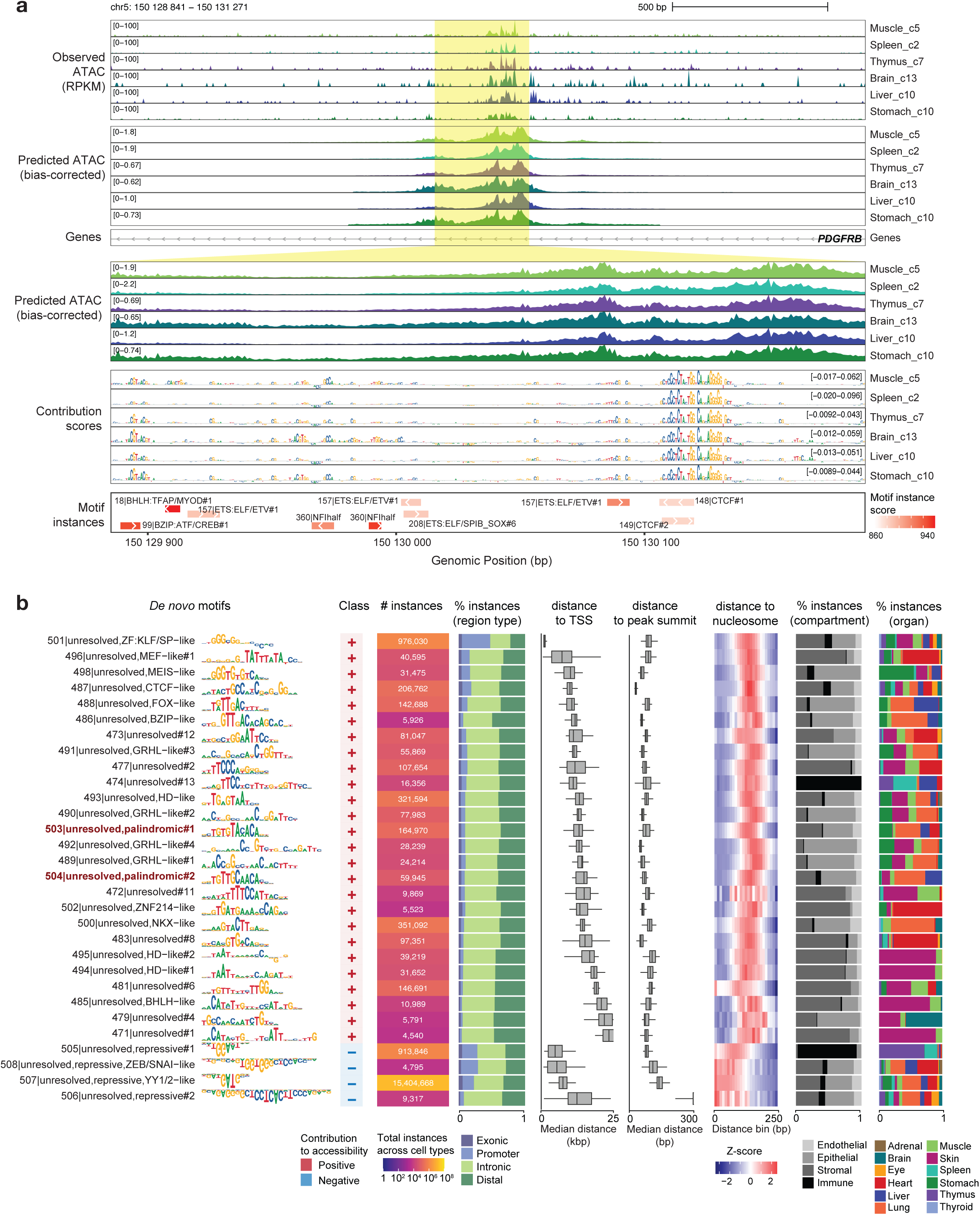
Modeling of sequence determinants of chromatin accessibility. **a)** Visualization of *PDGFRB* locus with observed and predicted accessibility and basepair-resolution contribution scores for several endothelial cell types from different tissues. **b)** Summary of unresolved *de novo* motifs not matching known motifs. From left: CWM representation, motif class (based on contribution to accessibility), total number of instances across cell types, and distribution of motif instances by organ. Two palindromic unresolved motifs are indicated in red text.

**Figure S6, related to Figure 4.**
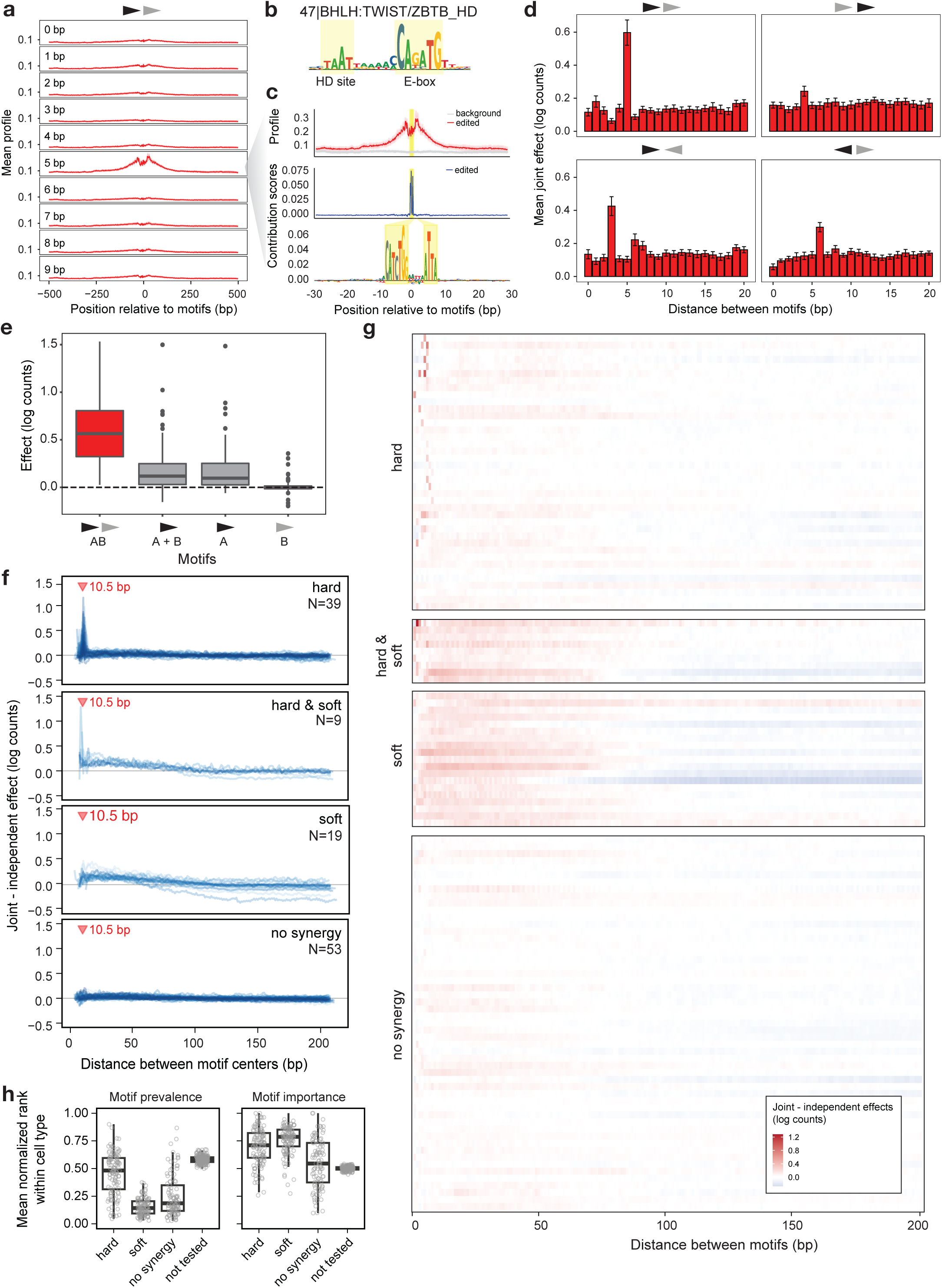
Motif synergy and syntax preferences. **a-e)** Synergy analysis for Coordinator-like composite motif. **a)** Predicted profile for Coordinator constituent motifs (BHLH, HD) inserted at 0-9 bp distance between motifs, for one arrangement. **b)** CWM for composite motif. **c)** Predicted profile at 5 bp distance between Coordinator motif constituents, and mean contribution scores after model interpretation of edited sequences with motifs inserted. **d)** Quantified effects in log counts, per orientation (error bars: standard deviation across folds). **e)** Distribution of predicted effects across 100 sequences, for both motifs in their optimal arrangement, the sum of motifs A and B inserted independently, and motifs A and B alone. **f)** Difference between predicted joint effects for all composite motifs tested, and sum of independent effects. The best orientation for each motif is shown. **g)** Heatmap representation of joint minus independent effects as in (f). **h)** Distribution of mean normalized rank across motifs in each synergy group within each cell type, for motifs ranked based on prevalence defined by number of predictive instances (left) or importance defined by summed contribution scores across the motif CWM (right) (see Methods). Two-sided Wilcoxon rank-sum test p values for prevalence: hard vs soft (1.07×10^-26^), hard vs no synergy (7.15×10^-13^), hard vs not tested (1.33×10^-20^); and for importance: hard vs no synergy (5.34×10^-7^), hard vs not tested (1.33×10^-20^), soft vs no synergy (8.00×10^-13^), soft vs not tested (2.83×10^-20^). Only positive motifs were considered.

**Figure S7, related to Figure 6.**
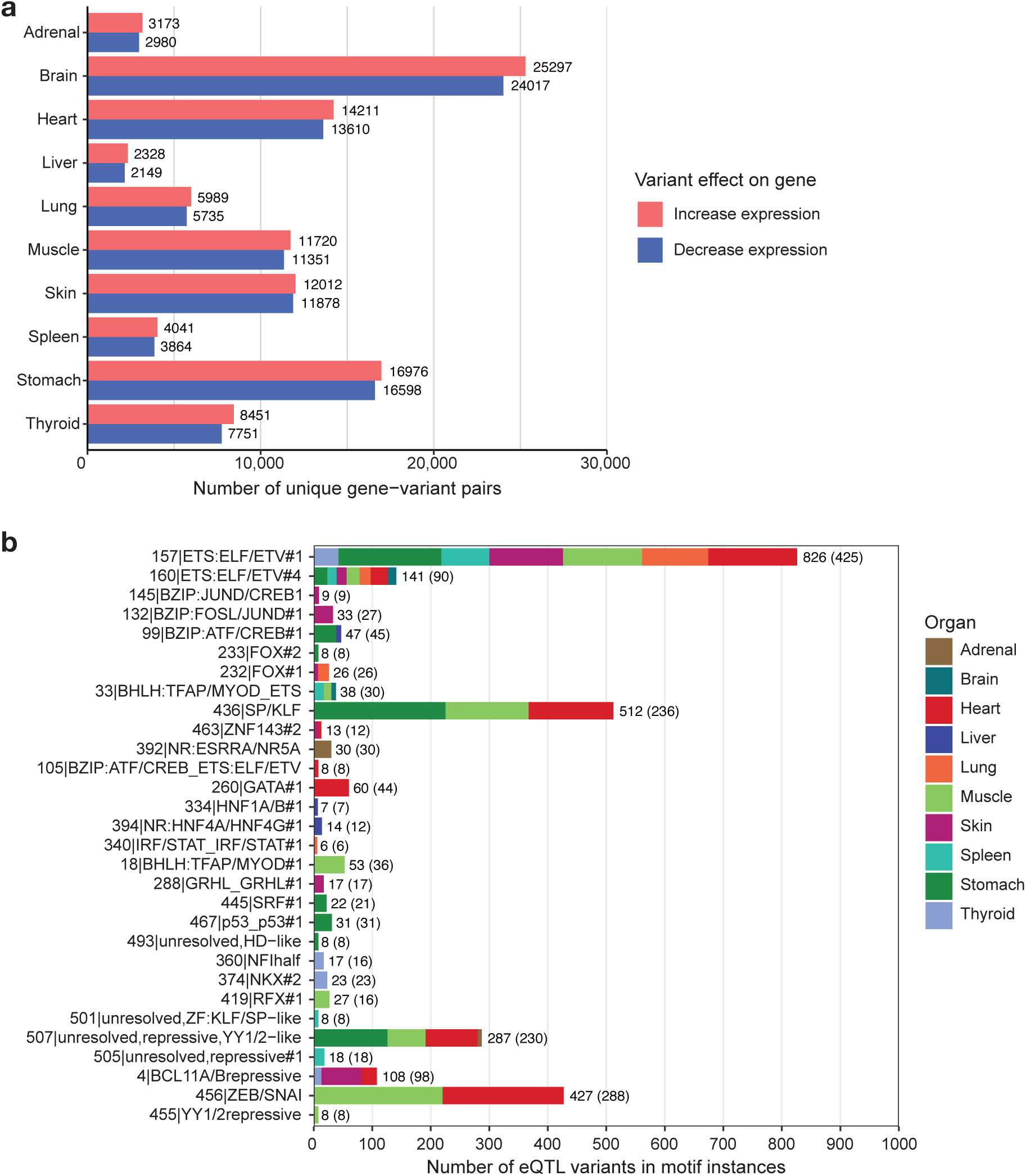
Tissue-specific overlaps between eQTL variants and HDMA motifs. **a)** Number of unique eQTL variant-gene pairs examined per organ. **b)** Motifs significantly enriched in eQTL variants. Numbers state the total number of variants within all instances of the respective motif across all organs, with the total number of unique gene-variant pairs in parentheses.

**Figure S8, related to Figure 7.**
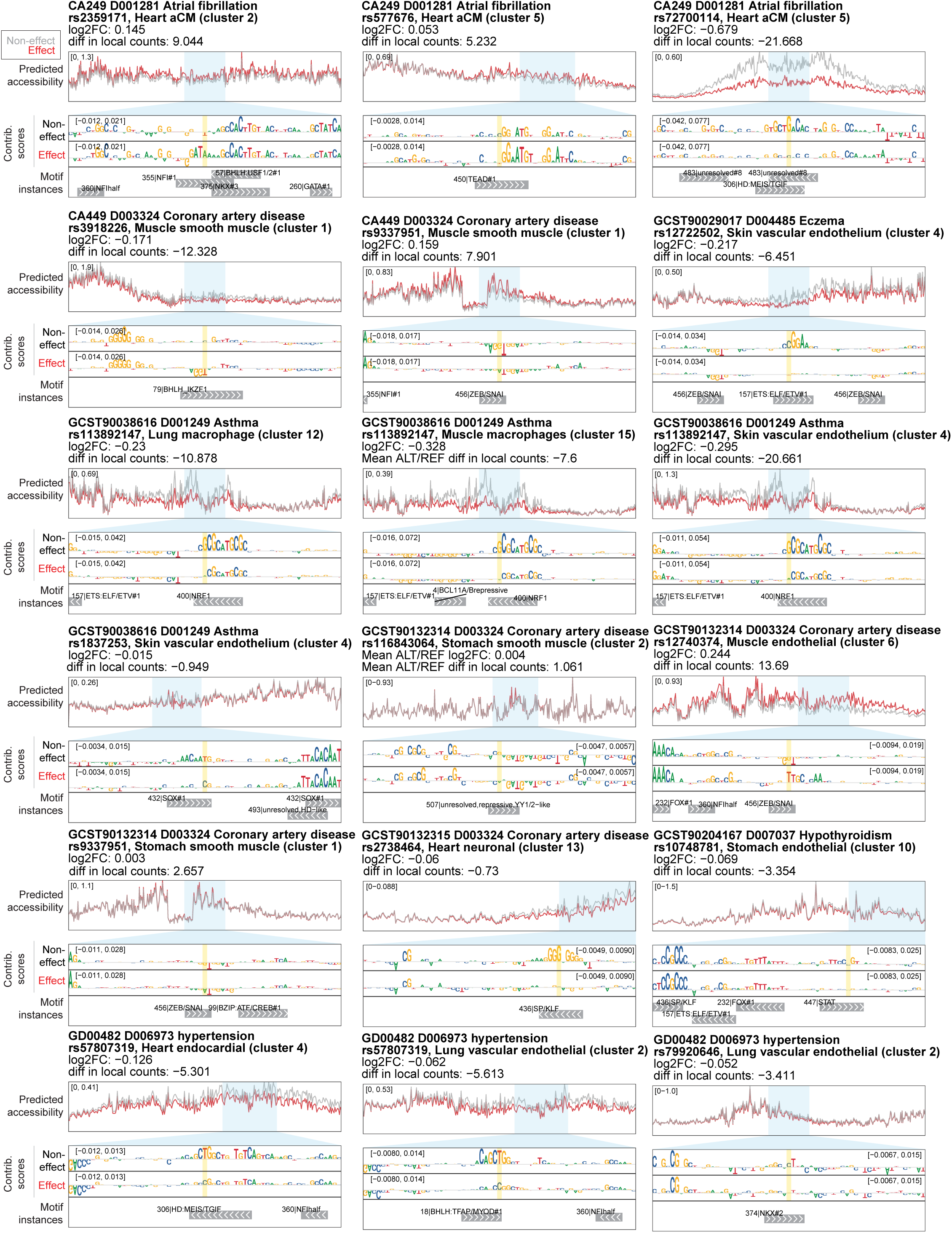
Prediction of variant effects in selected fetal-only hits. For each hit, predicted accessibility for the effect and non-effect alleles are shown for approximately +/- 200 bp around the variant site. Blue highlights indicate the region +/- 30 bp around the variant site displayed in the tracks below. Tracks, in order from top to bottom, represent per-basepair contribution scores for predicted accessibility for non-effect and effect alleles, and motifs called in the respective cell types (with the non-effect allele). Yellow highlights indicate the affected nucleotide. “log2FC” indicates the mean log2 fold change between predicted counts for the effect allele vs the non-effect allele. “Diff in local counts” indicates the difference in predicted counts between the effect allele and non-effect allele in the 100 bp surrounding the variant.

## Supplementary Tables

Table S1. Sample meta data + QC thresholds per sample, related to Fig. 1

Table S2. Cluster annotations and QC metrics, related to Fig. 1

Table S3. Cluster marker genes, related to Fig. 1

Table S4. L3 cell cluster highly regulated genes (HRGs) and global HRGs, related to Fig. 2

Table S5. ChromBPNet model QC, related to Fig. 3

Table S6. Motif lexicon summary, related to Fig. 4-5

Taable S7. Results of in silico tests for synergy, related to Fig. 4-5

Table S8. eQTL tables (S8.1) and fisher exact test statistic (S8.2), related to Fig. 6

Table S9. g-chromVAR results, related to Fig. 7

Table S10. Causal variants linked to relevant traits, related to Fig. 7

Table S11. List of motifs with disease causal variants, related to Fig. 7

Table S12. List of ENCODE accessions of the adult snATAC-seq peak sets, related to Fig. 7

Table S13. Fetal-only hits for variant scoring, related to Fig. 7

Table S14. List of uploaded files, file types and Zenodo depositions they can be found in

## Supplementary Notes

Note S1. Greenleaf Lab SHARE-seq protocol.

Note S2. Quality control per organ.

## Methods

### Sample collection

De-identified tissue samples were obtained at Stanford University School of Medicine from elective pregnancy terminations under a protocol approved by the Research Compliance Office at Stanford University. Tissue samples were delivered on ice and immediately stored in liquid nitrogen prior to processing.

### Nuclei isolation

A multi-tissue compatible nuclei isolation protocol was developed to efficiently isolate stable nuclei for further library preparation. Briefly, for a given sample, 100-200 mg of tissue was added directly into 1mL of Nuclei Extraction Buffer (250mM Sucrose, 25mM KCl, 5mM MgCl2, 20mM HEPES-KOH, 65mM beta-glycerol, 0.5% IGEPAL CA-630, 1x protease inhibitor, 1mM DTT, 0.2mM Spermine, 0.5mM Spermidine, 60U/mL RNasin Plus, 2-5% normal goat serum) in a chilled 2mL dounce homogenizer (Kimble #885300-0002) on ice. The sample was incubated for 10 minutes on ice. The sample was dounced 20 times each with pestle A then with pestle B. Sample was transferred to a DNA low binding tube. 300µL additional Nuclei Extraction Buffer was used to rinse any remaining nuclei from dounce homogenizer. Sample was incubated with vertical rotation for 5 minutes at 4°C. Sample was filtered using a 70µm Flowmi strainer. Volume was adjusted with additional Nuclei Extraction Buffer to 1.2mL total volume. 37% formaldehyde was added to the sample for a 0.2% final formaldehyde concentration and incubated for 4 minutes at room temperature with vertical rotation. Fixation was quenched with 125mM glycine for 8 minutes at room temperature with vertical rotation. Nuclei Extraction Buffer was added to the sample for a final volume of 1.4mL. An iodixanol gradient was prepared to enrich nuclei from homogenate. Briefly, 50% iodixanol solution was prepared from 60% iodixanol with the addition of 1mM DTT, 60U/mL RNasin Plus, and 2-5% normal goat serum. The sample was mixed with an appropriate amount of iodixanol for a final 22% iodixanol concentration. 44% iodixanol solution was layered below the sample. Then a 22% iodixanol solution was gently added between the sample and the 44% iodixanol solution layer. The sample was centrifuged at 3500xg for 30 minutes at 4°C with brakes off. The nuclei layer was separated with gentle pipetting for further processing.

### SHARE-seq library preparation

See full protocol in **Note S1**, adapted from published SHARE-seq protocols^23,115^.

### Library sequencing

All DNA libraries were sequenced on a NovaSeq 6000 using 300-cycle S4 v1.5 reagent kits with XP workflow. Paired-end sequencing was run with a 96-99-8-96 configuration (Read1-Index1-Index2-Read2). We quantified DNA libraries using Qubit and tapestation, then prepared library pools at 1.5nM concentration for a final loading concentration of 300pM. Sequencing was performed at the Stanford Genome Technology Center.

### VISTA embryo histology and immunochemistry

We received X-Gal-stained and fixed whole mouse embryos in PBS from Dr. Len Pennachio^27,28^ and transferred them to 70% EtOH for storage. Paraffin embedding was performed by Histo-Tec Laboratory (Hayward, CA) using a xylene-free dehydration protocol as xylene could dissolve the X-Gal stain. Briefly, the embryos were sequentially dehydrated with 80%, 95%, 100%, 100%, 100% EtOH for 20 minutes each, followed by washes with 50:50, 80:20, 90:10, 100:0 paraffin:alcohol mix for 20 minutes each to remove the EtOH. Subsequent embedding and Hematoxylin & Eosin staining was performed with standard protocols on 5um sections.

### SHARE-seq data preprocessing

We developed a highly parallelized, rapid, and storage-efficient pre-processing Snakemake (v7.15.1)^116^ pipeline to convert BCL files from sequencers to ATAC fragment files and RNA sparse matrices (**Fig. S1**). Briefly, raw BCL files were first converted to FASTQ files using a custom script that parallelizes the *bcl2fastq* (v2.20.0.422, Illumina) conversion by flow cell tiles, parses the read cycles, and demultiplexes the raw FASTQ files into sublibraries based on sublibrary barcodes in the Index2 reads. For each sublibrary, we further split the FASTQ file into random chunks of 20 million reads.

Within each chunk of an ATAC sublibrary, we performed barcode matching against the SHARE-seq barcode whitelist, allowing for 1 bp mismatch for each of the three rounds of 8 bp barcodes that make up a single-cell barcode, followed by Nextera adapter trimming with *fastp* (v0.23.2)^117^, genome alignment with *Bowtie2* (v2.5.0)^118^, and conversion of the output BAM file to a more storage-efficient fragment file. We then merged the fragment files from all chunks of a sublibrary, deduplicated fragments per cell based on start and end coordinates, and demultiplexed the fragments into samples based on round 1 cell barcodes. Finally, for each sample, we merged the demultiplexed fragment files for that sample across all sublibraries to generate the final ATAC fragment files (*.fragments.tsv.gz, *.fragments.tsv.gz.tbi).

Within each chunk of an RNA sublibrary, we performed barcode matching, 10 bp UMI parsing from Read2, and adapter trimming for Read1 only, followed by genome alignment with *STAR* (v2.5.4b)^119^, gene annotation with *featureCounts* (v2.0.1)^120^, and conversion of the output BAM file to a more storage-efficient TSV format. We then merged the annotated TSV files from all chunks of a sublibrary, split into 12 barcode chunks based on round 3 barcodes, deduplicated UMIs per cell per annotated gene per barcode chunk using *UMI-tools* (v1.1.2)^121^, demultiplexed the deduplicated TSV files into samples based on round 1 cell barcodes, and converted the TSV files into the Matrix Market Exchange format. Finally, for each sample, we merged the demultiplexed Matrix Market Exchange files for that sample across all sublibraries to generate the final RNA sparse matrix files (*.matrix.mtx.gz, *.features.tsv.gz, *.barcodes.tsv.gz).

On average, we can process a 10B-read NovaSeq run in under 4 hours using an academic high performance computing cluster. This pipeline can be easily adapted to process other split-and-pool-based single-cell multiomic data. All libraries were aligned to the hg38 reference genome.

The pipeline is available at https://github.com/GreenleafLab/shareseq-pipeline (stable release v1.0.0).

It is well known that in ATAC-seq experiments, Tn5 transposase forms a homodimer with a 9-bp gap between the two Tn5 molecules, resulting in two insertions 9-bp apart on different DNA strands per accessible site^4,122^. When sequencing the DNA fragments using paired-end sequencing, the start and end positions need to be adjusted based on the insertion offset of Tn5 to reflect the true center of the accessible site at the midpoint of the Tn5 dimer. To account for the Tn5 offset, previous ATAC-seq studies used a +4/-5 offset approach where plus-stranded insertions are adjusted by +4 bp, and minus-stranded insertions by -5 bp. However, this in fact results in a 1 bp mismatch of the adjusted insertion sites between the two fragments sharing a single transposition event (**Fig. S1b**). The discrepancy may stem from the end-exclusive coordinate system used by BAM and BED files, as the original +4/-5 convention is only correct if the output file is interpreted in a non-standard 0-based, end-inclusive genomic coordinate system. This mismatch does not affect most downstream ATAC analysis that bins insertions on the scale of hundreds of basepairs, but it does affect basepair-sensitive analysis such as TF footprinting and motif analysis. In this SHARE-seq preprocessing pipeline, we have adopted the +4/-4 offset instead, which results in a consensus insertion site. See example motifs generated from reads corrected by either of these offset schemes in Figure S3a of Pampari et al^17^.

### SHARE-seq data QC and filtering

We performed per-sample QC filtering by manually inspecting and thresholding the following metrics: 1) TSS enrichment ratio and number of fragments for ATAC fragment files, 2) number of UMIs, number of genes and percentage of mitochondrial reads for RNA sparse matrices, 3) ratio of RNA UMIs vs ATAC fragments to remove cells with low quality in one modality (**Fig. S2a**). All sample filtering thresholds are summarized in **Table S1**.

### RNA normalization, ambient RNA removal, dimensionality reduction, and clustering

We used Seurat (v4.3.0)^123^ in R (v4.1.2) to process filtered RNA sparse matrices into Seurat objects per organ^123^. We adopted an iterative dimensionality reduction and clustering workflow to sequentially annotate cell types and filter out additional low quality clusters (**Fig. S2a**). For each iteration, we first performed SCTransform v2 and variable feature selection on RNA raw counts of each sample, then selected the top 3,000 consensus variable features across samples using the *SelectIntegrationFeatures* function from Seurat, excluding mitochondrial genes, sex chromosomes genes, and cell cycle genes to minimize batch effects. We merged the raw RNA counts from per-sample objects into a single matrix, performed SCTransform v2 using consensus features, and used the DecontX function from the celda (v1.6.1) package ^124^ on SCT-corrected counts to remove ambient RNA contamination per cell. The decontaminated counts were then split by sample, scaled to 10,000 UMIs per cell and log normalized. Similar to the process mentioned above, we selected a list of top 3,000 consensus variable features from the per-sample variable features. PCA was performed on the merged object with the consensus features, followed by cell clustering using the Louvain algorithm at a resolution of 0.3 with 50 PCs and UMAP embedding. We then inspected each cluster and removed any low quality clusters with significantly lower UMIs than other clusters, high levels of co-expression for different tissue compartment markers that are biologically impossible and suggestive of doublets (e.g. high expression for both epithelial and endothelial compartment markers), or no clear cell type-defining marker genes. After removing cells in the low quality clusters, we repeated the processing steps starting from RNA raw counts for each sample. This process was repeated until no more low quality clusters were identified, which usually required 1-3 iterations. Cells in the final set of clusters passing this iterative QC were considered “whitelisted”. For each cell type cluster, marker genes were identified in a one-vs-all Wilcoxon rank-sum test versus all other clusters from the same organ, and filtered to genes with a log2 fold change greater than 1.

All final cluster annotations are included as **Table S2**. All cluster markers are summarized in **Table S3** and a subset is visualized in marker gene dot plots in **Note S2**. All UMAP embeddings are included in **Note S2**.

### ATAC peak calling, motif enrichment, and chromVAR

We used ArchR (v1.0.2)^125^ to process filtered ATAC fragment files into ArchR projects per organ. After filtering to the final whitelisted cell barcodes from the iterative RNA processing workflow and transferring the clustering and cell type annotations, we called peaks per cluster using Macs2 (v2.2.7.1)^126^, merged peaks into a single reproducible peak set per organ using ArchR’s iterative overlap strategy, and created a cell-by-peak matrix of fragment counts. We identified marker peaks per cluster using a Wilcoxon rank-sum test and performed TF motif enrichment within the marker peaks with a cutoff of FDR <= 0.1 and log2FC >= 0.5. We calculated chromVAR motif deviations across all clusters within each organ^24^. For both of these analyses, we used a curated cisBP motif set of 1,141 unique human TF motifs described in Kartha *et al* 2022^127,128^. We created a global ArchR project by merging all 12 per-organ ArchR projects and an HDMA global accessible candidate cis-regulatory element (acCRE) set by iteratively overlapping peak sets called from individual clusters across all organs^125,129^.

### Linkage of regulatory elements to genes with modified Activity-By-Contact (ABC) model

We used the ABC approach to link acCREs to gene promoters. To ensure consistency of ABC enhancer regions as those in our HDMA global acCREs set, we adapted ABC^26^ to enable custom regions as inputs to the model. To create the custom region set, we used bedtools to merge the HDMA global acCREs set with the hg38 genome transcription start site set. We used the pseudoubulk ATAC-seq signal as enhancer activity and estimated 3D contact frequency between enhancers and promoters using a power law function of genomic distance. ABC was run on each L1 cell cluster. The results were filtered for enhancer-promoter links with an ABC score greater than 0.013, which corresponds to a 70% recall rate from the benchmark CRISPR dataset. Our modified ABC workflow is available at: https://github.com/GreenleafLab/ABC-Enhancer-Gene-Prediction-CustomRegions (commit b3d2156).

### VISTA enhancer analysis

We filtered for VISTA-validated enhancers that originated from humans or have a human sequence homolog and annotated as X-Gal positive in any organs present in HDMA (accessed 2024-01-24), overlapped with HDMA global acCREs, and retained VISTA enhancers with a minimum of 75% (375 bp) overlap. If multiple acCREs overlapped the same VISTA enhancer, we chose the acCRE with the highest ATAC signal, based on previous observations that VISTA enhancers often have a much smaller core element and enhancer activity does not depend on all regulatory elements within an enhancer^130^. ATAC signal was scaled and log normalized per L1 cluster then Z-scored across clusters per enhancer. For each organ, a one-sided Wilcoxon rank-sum test was performed to calculate the statistical significance of the HDMA ATAC signal enrichment in acCREs overlapping VISTA enhancers annotated as positive in that organ. For example, to test the significance of brain ATAC signal enrichment, we first subsetted to HDMA brain clusters only, then compared ATAC signal in acCREs overlapping VISTA brain-positive enhancers and acCREs overlapping VISTA brain-negative enhancers using a one-sided Wilcoxon rank-sum test. The effect size was calculated as the W statistic / (n1 * n2), where n1 is the number of acCREs in the brain-positive group and n2 is the number of acCREs in the brain-negative group. This effect size represents the AUROC probability that a given acCRE in the brain-positive group will have higher ATAC signal than an acCRE in the brain-negative group. Similarly, for the RNA data, we first identified the nearest gene for each acCRE overlapping a VISTA enhancer, scaled and log normalized the raw RNA expression counts per L1 cluster, and then Z-scored expression values across clusters per enhancer. An analogous Wilcoxon rank-sum test was performed for each organ to assess the statistical significance of the HDMA RNA signal enrichment in VISTA positive enhancers.

### Preparation of input regions for ChromBPNet models

To define genomic regions for training ChromBPNet models, we performed a second round of peak calling to obtain a lenient set of accessible regions. First, pseudobulk fragment files for each of the 203 L1 cell type clusters were generated by concatenating fragments from the SHARE-seq ATAC modality for all cells in that cluster, from all samples. For each pseudobulk, we then derived pseudoreplicates. For each fragment, starts and ends (corresponding to Tn5 insertion sites) were randomly allocated to each of two pseudoreplicate files, and pseudoreplicate files were also concatenated into a total-pseudoreplicate file. Macs2 (v2.2.9.1) was used to call peaks on all three pseudoreplicate files with parameters: *-p 0.01 --shift -75 --extsize 150 --nomodel -B --SPMR -- keep-dup all --call-summits*. Only peaks called on the total-pseudoreplicate which overlapped peaks called in both pseudoreplicates were retained. Peaks overlapping the GRCh38 ENCODE blacklist (ENCODE accession ENCFF356LFX) were excluded. Peak coordinates were adjusted to 1,000 bp centered at the Macs2 peak summit. Pseudoreplicates were only used for peak calling, and pseudobulk fragment files were used for downstream model training.

We used the ChromBPNet package (https://github.com/kundajelab/chrombpnet, commit a5c231) and followed the workflow described by Pampari et al^17^. We used the command *chrombpnet prep nonpeaks* to define background regions which match the GC content of peak regions. For each cell type, we used a five-fold cross-validation scheme, where each fold (designated 0 to 4) comprised a different set of training, validation, and test chromosomes, with each chromosome in the test set of at least one fold. We used the default human chromosome folds provided with ChromBPNet^131^.

### ChromBPNet model training and interpretation

ChromBPNet models are supervised convolutional neural networks trained to use 2,114 bp one-hot-encoded DNA sequence in peaks and background regions to predict the accessibility profile (as a probability distribution) and total natural log counts (as a scalar value) in the central 1,000 bp window of input regions. ChromBPNet models use a pre-trained bias model and explain the residual accessibility not captured by Tn5 enzyme bias. We trained a bias model to learn the enzymatic bias in the SHARE-seq setting using fold 0 for the heart L1 cluster 0 pseudobulk, with bias threshold factor -b 0.4 using the *chrombpnet bias pipeline* which also performs model interpretation using DeepLIFT. We confirmed that the bias model learned the Tn5 motifs but not transcription factor motifs, and used this bias model to subsequently train ChromBPNet models for five folds for each of the 203 L1 cell types (1,015 models total) using the *chrombpnet pipeline* command with the GRCh38 reference genome from ENCODE (fasta: https://www.encodeproject.org/files/GRCh38_no_alt_analysis_set_GCA_000001405.15/, with chromosome sizes from ENCODE accession ENCFF667IGK). Models were evaluated based on the Pearson and Spearman correlations between predicted and observed log counts in peaks and the Jensen-Shannon Distance between predicted and observed profiles in peaks, for peaks on held-out test-set chromosomes (**Table S5, Note S2**). Models for four cell types where the Spearman correlation for any fold was less than 0.5, generally corresponding to pseudobulks with low coverage, were excluded. To generate the average predicted accessibility tracks across folds for peak regions (representing counts per base), for each region, the mean predicted profile logits across folds were softmaxed to convert them to probabilities, then scaled by the exponentiated mean predicted log counts across folds.

We performed model interpretation to determine the extent to which each nucleotide was predictive for accessibility. We ran the *chrombpnet interpret* command which uses the DeepLIFT^35^ algorithm to compute contribution scores for each nucleotide in the 2,114 bp input windows with respect to the predicted counts. Contribution scores were derived for each model fold for all peak regions, and the mean computed across folds. The averaged predicted accessibility profiles and contribution scores were converted to bigWig files, as well as used for all analyses and figures.

### *De novo* motif discovery and assembly of the motif lexicon

Assembly of the *de novo* motif lexicon required three steps: 1) *de novo* motif discovery per cell type, 2) collapsing similar motifs across cell types, and 3) motif annotation.

1. First, for *de novo* motif discovery on sequences driving chromatin accessibility, we used TF-MoDISco^36^ which briefly, identifies seqlets, corresponding to short spans of contiguous high positive-importance or high negative-importance nucleotides, and clusters them into recurrent 30 bp patterns. We used the implementation in the tfmodisco-lite package, https://github.com/jmschrei/tfmodisco-lite, v2.0.7) on the mean contribution scores for each cell type, sampling 1,000,000 seqlets for both positively- and negatively-contributing seqlets (parameter *-n 1,000,000*), and using the default behaviour to search for seqlets in the central 400 bp of input regions. Each *de novo* motif is represented by a 4x30 contribution weight matrix (CWM), computed as the mean contribution score per position per nucleotide across its seqlets, and a 4x30 position probability matrix (PPM), computed by normalizing the nucleotide frequencies per position across its seqlets. We manually inspected CWMs learned in each cell type, and the ChromBPNet models and motifs for ten cell types which predominantly learned low-complexity motifs were excluded from downstream analysis. This resulted in 189 cell types (945 models total) passing quality control and used throughout our study, which collectively learned 6,362 motifs including both positively-contributing and negatively-contributing motifs.
2. Next, we automatically consolidated the 6,362 motifs into a non-redundant set. We first derived clusters of motifs which were broadly similar. CWMs were trimmed by removing positions where the total contribution across nucleotides was less than 30% of the maximum total contribution among all positions^17^. PPMs were trimmed to the same positions as the trimmed CWMs, and converted to position frequency matrices (PFMs) by multiplying PPMs by the total number of seqlets associated with each motif. PFMs from all cell types in each organ were first leniently clustered, separately for positive and negative motifs, using the gimmemotifs package^132^ (v0.18.0, with command *gimme cluster -t 0.8*), which returns an average PFM for each motif cluster. Average PFMs from across organs were then subjected to a second round of clustering with gimmemotifs. Within each broad motif cluster, we then collapsed the constituent CWMs originating from individual cell types using the SimilarPatternCollapser functionality in TF-MoDISco, which merges together similar motifs using the same method it does for seqlets during initial motif discovery in step (1). This procedure resulted in 834 motifs.
3. Finally, we performed motif quality control, and annotated and categorized each motif. For annotation, we used TOMTOM^133^ (v4.11.2) to compute similarities between the 834 *de novo* motif CWMs and a curated set of 5,193 known TF binding site position weight matrices (PWMs) from JASPAR, CIS-BP, and other studies (obtained from https://resources.altius.org/~jvierstra/projects/motif-clustering-v2.1beta/), using the command *tomtom -no-ssc -oc . --verbosity 1 -text -min-overlap 5 -mi 1 -dist pearson -evalue-thresh 10.0*. For every motif, we manually inspected the most similar PWMs to assign a provisional label, typically at the TF family level. We further collapsed highly similar motifs missed by the clustering approach, retaining the motif with the highest number of seqlets across cell types. We flagged 107 motifs which were low-complexity, noisy, or dominated by CG dinucleotides for exclusion. We categorized motifs as “base” if they matched known PWMs; “base with flanks” if they matched known PWMs but exhibited additional high-contribution nucleotides; and “homocomposite” or “heterocomposite” if the motifs clearly matched two similar or distinct known motifs, respectively. Motifs which did not resemble known PWMs were labelled as “unresolved”. After exclusion of low-quality motifs, this resulted in a set of 508 non-redundant motifs used for downstream analysis (**Table S6**). Motif labels have the naming scheme “<UNIQUE_ID>|<FAMILY>:<SUBFAMILY>/<ALTERNATIVE_SUBFAMILY>#<INDEX>”. The unique ID is a value from 0 to 508 (one ID corresponding to the low-quality motifs was filtered out). The index is used to distinguish separate motifs which each match the same known motif, typically representing subtle variations in nucleotide preferences or flanks. Composite motifs used a similar naming scheme, with labels for constituent motifs separated by underscores. Motifs were also assigned a broad label (e.g. “CTCF” and “CTCFupstream” motifs shared a broad “CTCF” label), used throughout figures and analyses to aggregate results.

### Identification of predictive motif instances

To identify genomic instances of *de novo* motifs in each cell type, we used the Fi-NeMo package^38^ (https://github.com/austintwang/finemo_gpu, v0.23, commit b81876d), which performs motif scanning in a contribution-aware manner. Briefly, Fi-NeMo fits a sparse linear regression model for each peak region to minimize the difference between the contribution scores in each region, and a reconstruction of the scores from a weighted combination of trimmed input CWMs. In the coefficient matrix, a non-zero coefficient at a certain location indicates a motif instance in that location. For each cell type, the output of Fi-NeMo is a BED file of predictive instances across all peaks, representing short regions which both match motifs by sequence and have relatively high absolute contribution scores. We first ran Fi-NeMo with all 834 clustered CWMs, with parameters *–alpha 0.8 –trim-threshold 0.3* and defaults otherwise (parameter *alpha* is now known as *lambda*). Due to the competitive nature of motifs in the linear modelling approach, we ran Fi-NeMo a second time with a reduced set of 436 motifs after excluding composite motifs.

We performed post-hoc filtering of motifs to obtain a high-confidence annotation for downstream analysis. To evaluate the quality of instance calls for a given motif, Fi-NeMo computes the correlation between the input CWM and the CWM derived from averaging contribution scores for all Fi-NeMo-identified instances for that motif. For each cell type, instances from the two Fi-NeMo runs were concatenated, and motifs where correlation between the instance-CWM and the input-CWM was less than 0.9 were flagged. Next, all instances of these flagged motifs were filtered out. Finally, to reduce redundancy, if multiple instances with the same annotation overlapped by more than 3 bp, only the instance with the highest “hit_correlation” value was retained, representing the instance with the contribution scores having the highest similarity with the corresponding input CWM. This step resulted in final motif annotation for each cell type consisting of their predictive motif instances.

### Inference of nucleosome positions from ATAC modality

We used NucleoATAC^41^ to infer nucleosome position and occupancy from the SHARE-seq ATAC modality data. Briefly, to determine nucleosome occupancy, NucleoATAC models the observed size distribution of ATAC fragments as a mixture of nucleosomal and nucleosome-free fragments, and the maximum likelihood fraction of nucleosomal fragments at a given position is used as a continuous occupancy signal. We adapted the original code to take fragments as input (https://github.com/sjessa/NucleoATAC, v0.4.1), and ran the NucleoATAC workflow for each cluster pseudobulk fragment file in the same peak regions used to train ChromBPNet models. The outputs of the *nucleoatac occ* command were used downstream: the nucleosome dyad position calls were used for analysis of motif instances, and the per-nucleotide maximum likelihood fraction of nucleosomal fragments were used for visualization of nucleosome occupancy as genomic tracks.

### Annotation of motif instances

To annotate predictive motif instances with respect to genomic features, instances were assigned as occurring in promoters if they were within 2 kbp upstream or downstream of transcription start sites (TSS), exonic if they overlapped exons, intronic if they overlapped gene bodies but not exons, and distal otherwise. Genomic features definitions were based on the Bioconductor TxDb.Hsapiens.UCSC.hg38.knownGene (v3.14.0) annotation, corresponding to the UCSC knownGene track from GENCODE V38, and assembled using the createGeneAnnotation function in the ArchR package. Motif instance distance to TSS was computed as the distance between instance center positions and TSS defined in the same annotation. Similarly, for each cell type, motif distance to the nucleosome dyad or peak summit was computed by calculating the distance between instance center positions and the nearest dyad inferred with NucleoATAC, or the peak summit, respectively. For analysis, we counted motif instances in 10 bp bins from 0 to 250 bp from the dyad or peak summit.

### *In silico* marginalizations to assess motif synergy

We used the trained ChromBPNet models to assess motif synergy following previous approaches^19,49^. In this *in silico* marginalization strategy, the predicted effect of a short sequence on chromatin accessibility is quantified by inserting the sequence into a library of background, non-accessible genomic regions (replacing the central nucleotides); and predicting accessibility for each background and edited region with a forward pass through a trained ChromBPNet model. By averaging the difference in predicted natural log counts between the edited and background regions over many regions, we estimate the “marginal” effect of a sequence. Two or more sequences can be inserted to estimate joint effects of those sequences. Specifically, for two motifs A and B, we define the predicted log counts for a region with one motif or both inserted as *y*_*A*_, *y*_*B*_, and *y*_*B*_ respectively; and the predicted log counts for an unedited background region as *y*_0_. The marginal effects, in log counts, of motif A and B are *Δ*_*A*_ = *y*_*j*_ − *y*_0_ and *Δ*_*j*_ = *y*_*j*_ − *y*_0_ respectively, and their joint effect is *Δ*_*J*_ = *y*_*J*_ − *y*_0_ . We define the independent effects of motifs A and B as *Δ*_*S*_ = *Δ*_*A*_ + *Δ*_*j*_, corresponding to a log-additive model for independent effects, or multiplicative model in units of counts. Synergy can then be defined as a significant deviation of the joint effects *Δ*_*J*_ of two sequences from their independent effects *Δ*_*S*_.

To implement the synergy analysis, we used the tangermeme package^48^ (https://github.com/jmschrei/tangermeme, v0.4.3). We first filtered the set of *de novo* composite motifs such that each composite was composed of a unique pair of constituent motifs. For each constituent, we identified the base *de novo* motif with the highest number of motif instances, trimmed the CWM as above, and defined the consensus sequence as the nucleotide with the highest contribution score at each position in the trimmed motif. The motifs and associated sequences tested are presented in **Table S7**. Sequences were manually adjusted to further remove uninformative flanks or better match the composite motif, and deduplicated so that a pair of the same sequences was only tested once. 100 background regions were randomly selected from GC-matched background regions for each cell type. For a pair of two of the same motifs, there are three unique orientations, and for a pair of two distinct motifs, there are four unique orientations. We considered each combination of orientation and distance between motifs an “arrangement” of motifs. For each composite motif, the constituent motif sequences A and B were inserted at all possible orientations and distances (from 0 to 200 bp) in the 100 background sequences, and accessibility predicted for background and edited sequences using all five ChromBPNet model folds (for the cell type with the most predictive instances of that composite motif) to compute *Δ*_*J*_. Similarly, A and B were inserted alone to compute independent and sum of independent effects *Δ*_*A*_, *Δ*_*B*_, and *Δ*_*S*_. Effects are computed for each sequence and model fold and the mean effect is reported (**Table S7**). For joint effects, the effect of the motif pair at their optimal arrangement (*i.e.* the combination of orientation and distance with the greatest mean effect *Δ*_*J*_) is reported. We considered each motif pair at their optimal arrangement and used a Wilcoxon signed-rank test to test if the paired differences in joint and independent effects at that arrangement (*Δ*_*J*_ − *Δ*_*S*_) were significantly greater than 0. Multiple testing correction was performed using the Benjamini-Hochberg method. Composite motifs with adjusted p value < 0.001 and (*Δ*_*J*_ − *Δ*_*S*_) > 0.15 were annotated as synergistic. To confirm that inserted sequences were driving the predicted synergistic effects, we performed model interpretation using DeepLIFT as above on edited sequences, and verified that the sequences predictive of accessibility corresponded to the sequences we inserted. Composite motifs where predicted effects were driven by different nucleotides than the ones inserted were excluded from this analysis.

To define synergistic motifs with syntax preferences (*i.e.* with synergy limited to or increased at specific binding site arrangements), for each composite motif, we computed Z-scores across the joint effects *Δ*_*J*_ at all arrangements. Composite motifs which had any arrangement with an effect greater than four standard deviations from the mean (Z-score > 4) were annotated as having hard syntax. We also considered that motifs could have weaker long-range preferences, or soft syntax. Composite motifs with (*Δ*_*J*_ − *Δ*_*S*_) > 0.15 at any arrangement where constituent motifs were between 20 and 150 bp apart were annotated as having soft syntax.

To assess cell type-specificity of predicted synergistic effects, for select composite motifs, we repeated the *in silico* marginalization analysis, inserting the constituent sequences at their optimal arrangements and predicting their joint effect on accessibility using the ChromBPNet models for all 189 cell types. For each cell type, sequences were inserted into its respective background regions, and mean effects were computed across 100 sequences and five model folds.

### Ranking of motifs by prevalence and importance

To assess motif prevalence and importance, we focused on motifs learned in each cell type in the cell type-specific TF-MoDISco motif discovery step, and only considered positive motifs. To compute motif prevalence, for each cell type, we obtained the set of motifs learned in that cell type, and the number of motif instances in that cell type for the corresponding lexicon motifs. We ranked motifs within the cell type based on their number of instances, and normalized ranks by dividing each rank by the maximum rank in that cell type such that normalized ranks fell in the range [0, 1]. Similarly, to compute motif importance, for each cell type, we obtained the set of motifs learned in that cell type and summed the contribution scores across nucleotides and positions for the corresponding trimmed CWM. Finally, to compare prevalence and importance of different classes of motifs defined in the synergy analysis, we grouped motifs based on whether they had hard syntax, soft syntax, no predicted synergy, or were not tested (meaning the motif was not a composite motif, or filtered out of the synergy analysis as described above). Motifs with both hard and soft syntax were grouped with hard-syntax motifs. We then computed the mean normalized rank across motifs in each group, within each cell type.

### Enrichment of eQTL variants in motifs

We obtained tissue-specific GTEx v8 eQTL data^57^ and concatenated them by organ source to match our fetal organs, aggregating the tissues as follows: Adrenal gland, Brain (Amygdala, Anterior cingulate cortex BA24, Caudate basal ganglia, Cerebellar Hemisphere, Cerebellum, Cortex, Frontal Cortex BA9, Hippocampus, Hypothalamus, Nucleus accumbens basal ganglia, Putamen basal ganglia, Spinal cord cervical c-1, Substantia nigra), Heart (Artery Aorta, Artery Coronary, Atrial Appendage, Left Ventricle), Lung, Liver, Muscle (Artery Tibial, Skeletal), Skin (Not Sun Exposed Suprapubic, Sun Exposed Lower leg), Spleen, Stomach/Esophagus (Stomach, Esophagus Gastroesophageal Junction, Esophagus Mucosa, Esophagus Muscularis) and Thyroid. We obtained unique lists of variant-gene pairs per organ by selecting the variant with the higher posterior inclusion probability score in case of duplicates.

For each organ, the log2 allelic fold change effect sizes (aFCs) for each variant-gene pair was used to determine the direction of variant effect on gene expression (upregulating or downregulating expression). Separately for each organ, we concatenated all motif instances from each cell type, then deduplicated entries with identical motif names and genomic positions. The number of upregulating and downregulating variants that overlapped positive or negative motif instances from the matched fetal organ were counted separately to obtain observed counts. aFCs were then randomly shuffled and direction of effect reassigned before the counting was repeated. We performed 100,000 shuffles per organ. Enrichment scores were defined as observed counts divided by the mean of the 100,000 shuffled counts. Enrichment p values were calculated as the proportion of shuffles where shuffled counts were larger than observed counts. Multiple testing correction using the Benjamini-Hochberg method was applied to the p values and a motif was considered significantly enriched with upregulating or downregulating variants if the FDR was < 0.05 and the observed count was above the 95% confidence interval of shuffled counts for that type of eQTL variant. Fisher’s exact tests were performed separately for positive and negative motifs to compare the number of motifs with or without significant enrichment with upregulating or downregulating eQTL variants (**Table S8**).

### Enrichment of cell types with disease variants using g-chromVAR

We used g-chromVAR^64^ to compute cell type-specific enrichment of disease variants. Since all genetic variants in CAUSALdb^63^ are reported in hg19 coordinates, ChromBPNet model input peak sets from fetal cell types were lifted over from hg38 to hg19 coordinates, and these were used as input to g-chromVAR. All coordinate conversions were performed using liftOver^134^ as implemented in the rtracklayer (v1.54.0) R package^135^ and the *hg38ToHg19.over.chain.gz* or *hg19ToHg38.over.chain.gz* chain files obtained from the UCSC Genome Browser. SuSiE scores for each variant listed in CAUSALdb were separately collated for all 13,710 studies in the database. CAUSALdb contains variants where linkage disequilibrium information was missing in some GWASs, which affects the causal variant estimation in the process of fine-mapping. These variants were given default posterior inclusion probabilities (PIP) values of -1 in the database and were removed from our analysis. We separately ran g-chromVAR (v0.3.2) for each organ to obtain Z-scores and p values of cell type enrichments with credible variants per study. We used the PIP values generated by SuSiE^78^ and filtered out studies where the sums of the PIP values across credible variants in that study were <5 in order to remove studies wherein we would be underpowered, retaining results from 13,194 studies. Multiple testing correction using the Benjamini-Hochberg method was applied to all p values and a cell type was considered significantly enriched with variants from a particular study if the FDR was < 0.05. We next manually extracted the subset of the significant results corresponding to disease traits relevant to the organs in HDMA (see **Table S10** for the list of studies). Potentially causal variants (SuSiE PIP ≥ 0.8) were lifted over from hg19 to hg38 coordinates and overlapped with fetal motif coordinates. For each fetal cell type in our dataset, we then manually identified matching adult tissues and cell types with snATAC-seq data in ENCODE, and obtained their snATAC-seq pseudoreplicated peak sets from ENCODE^79^ (**Table S12**). Causal variants overlapping fetal motif instances were subsequently overlapped with the relevant adult peak sets (peaks with -log_10_ p value > 2) if available. Only variants found in fewer than 2 matched adult peak sets were considered as “fetal-only” hits.

### Prediction of variant effect using ChromBPNet models

We predicted and interpreted effects of specific noncoding variants on chromatin accessibility using trained ChromBPNet models, as we have done previously^7^. We used the tangermeme package for predictions and model interpretation. For each variant, we used the 1,000 bp model training peaks for the relevant cell type to extract the reference genome sequence for the peak which the variant overlapped. This sequence (extended equally on either side to 2,114 bp) was fed to all five fold trained ChromBPNet models for the relevant cell type, to obtain predicted accessibility profile and aggregate log counts in the peak. For each fold, to transform predicted profile logits into accessibility profiles, the profile logits were softmaxed and scaled by the exponentiated predicted log counts; and model interpretation with respect to the counts output was performed using DeepLIFT. Next, the effect allele was substituted into the sequence at the variant position, and predictions and contribution scores were obtained as for the reference sequence. For each model, we computed the variant effect as the sum of differences in per-base predicted read counts in the 100 bp window centered at variant, and computed the mean effect score across folds. We also computed the log2 fold change between predicted counts for the effect versus the non effect allele for the peak region, where a log2 fold change > 0 indicates the effect allele was predicted to increase accessibility. In figures, the mean predicted profiles and contribution scores across folds are shown.

### Genome browser visualizations

For all data visualization at specific genomic loci, we used the BPCells R package ^136^ (https://github.com/bnprks/BPCells) along with custom scripts included in our code repository. All genomic tracks are hosted online for interactive visualization with the WashU Genome Browser (https://epigenomegateway.wustl.edu/browser2022/?genome=hg38&hub=https://human-dev-multiome-atlas.s3.amazonaws.com/tracks/HDMA_trackhub.json).

### Statistics and reproducibility

No statistical method was used to predetermine sample size. No data were excluded from the analyses. The experiments were not randomized. The investigators were not blinded to allocation during experiments and outcome assessment. For boxplots throughout the figures, the elements represent the following: center line, median; box limits, upper and lower quartiles; whiskers, 1.5× interquartile range.

## Data availability

All data (including fragment files, counts matrices, cell annotations, global acCRE annotations, ChromBPNet models, motif lexicon, motif instances, and genomic tracks) are deposited at https://zenodo.org/communities/hdma. A description of all data types and the corresponding URLs is provided in **Table S14**.

## Code availability

All analysis code is available at https://github.com/GreenleafLab/HDMA.

