## Supplementary Note 1 for "Dissecting regulatory syntax in human development with scalable multiomics and deep learning"

### Supplementary Note 1: SHARE-seq protocol

#### General notes

- Before starting the experiment, clean materials with RNase Away and 70% ethanol, use autoclaved tips and tubes. Maintain samples and solutions ice-cold during the whole experiment if not specified otherwise.
- Default spin for cells at 500xg for 5 min w/ swing bucket centrifuge.
- Start with 100k-1M cells for fixation (more the better but ensure enough combinatorial space for final sublibraries)
  - Expect ~20% of original starting number of cells by library prep stage.
- Resuspend oligos to appropriate concentrations in IDTE buffer:
  - 1mM = linker oligos
  - 100µM = RT primer, TSO, barcoded oligos
- Add all RNase inhibitors ***fresh*** to mastermix***.***
- Enzymatics RI is the only RNase Inhibitor that seems to be compatible with Tn5 activity, so we switch cells from the PBS-2RI into PBS-RI buffer before ATAC.
- Some washing steps specifies keeping pellet/beads stable and adding buffer directly ***without*** disturbing the pellet/beads. This does affect final recovery.
- When removing supernatant, use a two-step removal method: first remove the majority of supernatant with P1000, and then remove the remaining amount with P200. It may help to tilt the 2mL tube completely horizontally as you get close to removing all the liquid to avoid removing the pellet while removing as much liquid as possible.
- 2mL lo-bind Eppendorf tubes has better cell/nuclei recovery than 1.5mL tubes only when centrifugation is performed in a swing bucket because it's easier to completely aspirate the supernatant from 2mL tubes without disturbing the pellet at the tip of the 2mL tube.
- We split the bulk ATAC and RT reaction per sample into smaller tubes because the reaction efficiency is better with a smaller volume (even under the same concentration of reaction components).
- Where possible, check cell/nuclei quality and count using Countess at every major steps (ideally: pre-fix, pre-ATAC, pre-splitpool, post-ligation. If necessary, can get away without counting at pre-fix).
- Check before every run to ensure sufficient reagents in stock.

**Stock Buffer recipes:**
These buffers can be made in big batch ahead of time and stored for a few months

**Nuclei Isolation Buffer (NIB) (Keep at 4˚C)**

| Stock solution | Final conc. | Vol to add (mL) |
|---|---|---|
| 1M Tris HCl **pH 7.5** | 10mM | 0.5 |
| 5M NaCl | 10mM | 0.1 |
| 1M MgCl2 | 3mM | 0.15 |
| 10% NP40 (IGEPAL CA-630) | 0.1% | 0.5 |
| Ultrapure water | | 48.75 |
| | Total | 50 |

**2x RCB (Keep at room temp)**

| Stock solution | Final conc. | Vol to add (mL) |
|---|---|---|
| 1M Tris HCl **pH 8.0** | 100mM | 1 |
| 5M NaCl | 100mM | 0.2 |
| 20% SDS | 0.4% | 0.2 |
| Ultrapure water | | 8.58 |
| | Total | 10 |

**2x BW (Keep at 4˚C)**

| Stock solution | Final conc. | Vol to add (mL) |
|---|---|---|
| 1M Tris HCl **pH 8.0** | 10mM | 0.5 |
| 5M NaCl | 2M | 20 |
| 0.5M EDTA | 1mM | 0.1 |
| Ultrapure water | | 29.4 |
| | Total | 50 |

**1x B&W-T (Keep at 4˚C)**

| Stock solution | Final conc. | Vol to add (mL) |
|---|---|---|
| 1M Tris HCl **pH 8.0** | 5mM | 0.25 |
| 5M NaCl | 1M | 10 |
| 0.5M EDTA | 0.4% | 0.05 |
| 10% Tween 20 | 0.05% | 0.25 |
| Ultrapure water | | 39.45 |
| | Total | 50 |

**IDTE (oligo resuspension) (Keep at room temp)**

| Stock solution | Final conc. | Vol to add (mL) |
|---|---|---|
| 1M Tris HCl **pH 8.0** | 10mM | 0.5 |
| 0.5M EDTA | 0.1mM | 0.01 |
| Ultrapure water | | 49.5 |
| | Total | 50 |

**STE (Keep at room temp)**

| Stock solution | Final conc. | Vol to add (mL) |
|---|---|---|
| 1M Tris HCl **pH 8.0** | 10mM | 0.5 |
| 5M NaCl | 50mM | 0.5 |
| 0.5M EDTA | 1mM | 0.1 |
| Ultrapure water | | 48.9 |
| | Total | 50 |

**2x TD buffer (Keep at -20C)**

| Stock solution | Final conc. | Vol to add (mL) |
|---|---|---|
| 1 M Tris-HCl pH 7.4 | 20 mM | 2 mL |
| 1 M MgCl2 | 10 mM | 1 mL |
| Before the addition of DMF, adjust pH to 7.6 with 100% acetic acid | | |
| Dimethyl Formamide (DMF) | 20% | 20 mL |
| Ultrapure water | NA | Bring up to 100 mL |
| | Total | 100 mL |

**Split pool oligo plate preparation:**

Can be made in big batch ahead of time and stored at -20C for a few months

1. Thaw Round1, 2, 3 barcode oligo plates at room temp and let completely thaw (few hours or also try water bath).
    a. Very critical step that all oligos are at room temp before use.
    b. Spin the plates down using a plate adapter on a swing bucket or plate-compatible centrifuge.

2. Prepare each linker oligo dilution in tubes as below:
    a. 120μL Round 1 linker oligo (1mM) + 11,880μL STE buffer
    b. 120μL Round 2 linker oligo (1mM) + 9,480μL STE buffer
    c. 144μL Round 3 linker oligo (1mM) + 9,360μL STE buffer

3. Dispense following amounts of linker oligos into each 96 well PCR plate:
    a. 90μL diluted Round 1 linker oligo
    b. 88μL diluted Round 2 linker oligo
    c. 86μL diluted Round 3 linker oligo

4. Use a multichannel pipette to transfer the barcode oligo plates to the appropriate Round linker plate:
    a. 10μL Round 1 barcode oligos
    b. 12μL Round 2 barcode oligos
    c. 14μL Round 3 barcode oligos

5. Seal plates with aluminum adhesive cover and ensure that each well is well protected.

    Set up thermocycler as below:

| Temp | Time |
|---|---|
| 95˚C | 2:00 min |
| -1˚C/1 min | |
| 20˚C | 2:00 min |
| 4˚C | Hold |
| Total | ~1h 26min |

6. Check each well to make sure that there was not significant evaporation. If so, add appropriate amount of water to compensate.

7. Aliquot 10μL of the annealed oligos to a new plate (should be enough for 9x plates for each round). Use a liquidator or a liquid handling platform for distribution. Seal carefully to ensure proper seal at each well with aluminum foil seal and store at -20˚C.

**Buffer preparation**

Prepare following buffers fresh on the day of experiment, keep on ice unless otherwise specified.
Recommended order to prepare buffers: PBS-2RI, 2x TB Omni, 1x TB Omni, PBS-RI, NIB-RI.

Scale according to the number of samples and number of reactions needed

| **PBS-2RI** | **Vol for 1 sample (µL)** |
|---|---|
| 1xPBS | 3000 |
| 7.5% BSA | 16.05 |
| Enzymatics RI | 7.5 |
| SUPERase RI | 3.75 |
| Total | 3027.3 |

| **PBS-RI** | **Vol for 1 rxn (µL)** |
|---|---|
| 1x PBS | 5 |
| Enzymatics RI | 0.0125 |
| Total | 5.0125 |

| **NIB-RI** | **Vol for 1 sample (µL)** |
|---|---|
| NIB | 8323 |
| Enzymatics RI | 21 |
| SUPERase RI | 21 |
| Total | 8365 |

*6.565mL fixed volume of NIB-RI required given round 1 barcoding uses a single 96 well plate, another 1.8mL per sample required

| **2x TB (Omni)** | **Vol for 1 rxn (µL)** |
|---|---|
| 0.2M Tris-acetate | 8.25 |
| 5M K-acetate | 0.66 |
| 1M Mg-acetate | 0.5 |
| 10% Tween-20 | 0.5 |
| 1% Digitonin | 0.5 |
| 100% DMF | 8 |
| Ultrapure water | 6.59 |
| Total | 25 |

**ATAC (~20min)**

* Includes time-sensitive steps, make sure the appropriate buffers are ready ahead of time.
* This protocol assumes nuclei isolation, permeabilization, and fixation has already occurred during tissue dissociation.

1. Set swing bucket centrifuge to 4C.
2. Prepare 1xTB:

| **1x TB Omni** | **Vol for 1 rxn (µL)** |
|---|---|
| 2x TB (Omni) | 25 |
| Ultrapure water | 16.45 |
| PIC | 0.2 |
| Enzymatics RI | 0.85 |
| Total | 42.5 |

3. After appropriate nuclei dissociation protocol and fixation, mix the desired amount of cells with 500µL of PBS-RI and mix thoroughly, especially if it contains iodixanol.
4. Spin down at 500xg for 5 min at 4C in swing bucket centrifuge.
5. Prepare bulk transposition as follows, each reaction is ~40k cells. Prepare just enough based on the total number of ATAC reactions needed.

| **ATAC rxn** | **Vol for 1 rxn (µL)** |
|---|---|
| Sample resuspended in PBS-RI | 5 |
| 1xTB (Omni) | 42.5 |
| SHARE-ATAC Tn5 WJG Mar 2023 | 2.5 |
| Total | 50 |

   a. Resuspend each sample pellet in PBS-RI in appropriate volumes (5 µL per ATAC reaction).
   b. Assemble the ATAC reactions per sample with appropriate volume of 1xTB and Tn5. Tn5 is viscous and should be pipette mixed in slowly.
   c. Mix well and distribute 50µL per well to a lo-bind 96 well plate on a cooling block on ice (it's ok if the last well gets <50µL). Keeping plate cold is essential to control the Tn5 activity before the reaction starts. Seal plate, and incubate at 37˚C for 30 min with shaking at 500 rpm.
   d. At this time, thaw the 5x RT buffer, RT primer (100µM), and dNTPs.
6. Pool ATAC reactions by sample in 2mL lo-bind tubes, keeping the plate and tubes on ice as much as possible.
7. Spin down at 500xg for 5 min at 4C, remove supernatant.
8. Wash pellet gently ***without*** resuspending with 1mL NIB-RI.
9. Spin down at 500xg for 5 min at 4C.
10. Remove supernatant and resuspend pellet ***thoroughly*** in appropriate volumes of EB per sample (10µL per RNA RT reaction).

**RT (~45 min)**

1. Prepare RT mix (optimized for 100k cells per 50μL rxn):

| RT mix | Vol for 1 rxn (μL) |
|---|---|
| 5x RT buffer | 10 |
| Ultrapure water | 1.56 |
| dNTPs | 2.5 |
| Enzymatics RI | 0.31 |
| SUPERase RI | 0.63 |
| RT primer (100μM) | 5 |
| 50% PEG (wide bore tips) | 15 |
| Maxima H Minus RT (add right before rxn) | 5 |
| Total | 40 |

2. Start the RT protocol on thermocycler – the protocol should have a 50˚C hold so that when the samples are put in the machine, the hold is released and the 10min at 50˚C can be started right away.
3. Add appropriate volumes of RT mix to cells in EB (40uL RT mix + 10uL sample per reaction).
4. Split RT mix into PCR strips with 50μL/tube.
5. Set up thermocycler as below:

| Temp | Time | |
|---|---|---|
| 50˚C | Hold | |
| 50˚C | 10 min | |
| 8˚C | 12s | 3 cycles |
| 15˚C | 45s | |
| 20˚C | 45s | |
| 30˚C | 30s | |
| 42˚C | 2 min | |
| 50˚C | 3 min | |
| 50˚C | 5 min | |
| Total | 37 min | |

6. All steps below are at room temperature.
7. Thaw barcode oligos plates, block 1, 2, 3 oligos and T4 DNA ligase buffer at room temp. Place NIB-RI at room temp. Set centrifuge to room temp.
8. Add 300μL NIB-RI to a new 2ml lobind tube per sample and pool RT reactions into it, pipette mix well.
9. Spin down at 500xg for 5 min at room temp in swing bucket centrifuge.
10. Wash with 500μL NIB-RI **without** disturbing the pellet.
11. Spin down at 500xg for 5 min at room temp in swing bucket centrifuge.
12. Remove supernatant and resuspend pellet ***thoroughly*** with appropriate volumes of NIB-RI. The total volume for a single 96 well plate of hybridization is 1080μL NIB-RI. Divide by the number of independent samples.
    a. Count nuclei to confirm nuclei quality and cell count (pre-splitpool)

**Hybridization-ligation (~5hrs)**

1. Make sure that all oligo plates are thawed to room temp and spun down before hybridization.
2. Prepare hybridization buffer in 5mL tube:

| Hybridization buffer | Vol for 96 wells (µL) |
|---|---|
| Ultrapure water | 2589.3 |
| T4 ligase buffer (10x) | 540 |
| SUPERase RI | 13.5 |
| Enzymatics RI | 43.2 |
| 10% NP-40 | 54 |
| Total | 3240 |

3. Add appropriate volumes of hybridization buffer to resuspended samples in NIB-RI, mix well. If there are multiple samples, then divide the hybridization buffer appropriately for number of sample.
4. Using a multichannel trough, aliquot 40µL sample mixture to Round 1 plate, mix, and seal plate.
5. Shake at 300rpm for 30min at 25ºC.

6. Prepare blocking oligo 1 mixture:

| Blocking oligo 1 | Vol (µL) |
|---|---|
| Round 1 blocking (1mM) | 25.3 |
| T4 ligase buffer (10x) | 211.2 |
| Ultrapure water | 915.5 |
| Total | 1152 |

   a. Distribute 90µL each to a PCR strip (12x) and use a P20 multichannel to distribute blocking oligo and mix.
7. Using a whole new P10 pipette tip box, add 10µL blocking oligo 1 mixture to each well, mix, and seal plate.
8. Shake at 300 rpm for 30 min at 25ºC.

9. Pool samples into a trough, and using a whole new P200 pipette tip box, aliquot 50µL of mixture to Round 2 plate.
   a. Use a P200 multichannel to aspirate and pool into a new basin and redistribute using a multichannel.
10. Mix, seal plate, and shake at 300rpm for 30min at 25ºC.

11. Prepare blocking oligo 2 mixture:

| Blocking oligo 2 | Vol (µL) |
|---|---|
| Round 2 blocking (1mM) | 30.4 |
| T4 ligase buffer (10x) | 211.2 |
| Ultrapure water | 910.4 |
| Total | 1152 |

   a. Distribute 90µL each to a PCR strip (12x) and use a P20 multichannel to distribute blocking oligo and mix.
12. Using a whole new P10 pipette tip box, add 10µL blocking oligo 2 mixture to each well, mix, and seal plate.
13. Shake at 300rpm for 30min at 25ºC.

14. Pool samples into a trough, using a whole new P200 pipette tip box, aliquot 60µL of mixture to Round 3 plate.
   a. Use a P200 multichannel to aspirate and pool into a new basin and redistribute using a multichannel.
15. Mix, seal plate, and shake at 300rpm for 30min at 25ºC.

16. Prepare blocking oligo 3 mixture:

| Blocking oligo 3 | Vol (µL) |
|---|---|
| Round 3 blocking (1mM) | 26.5 |
| 10% NP-40 (can be replaced with water) | 11.5 |
| Ultrapure water | 1114 |
| Total | 1152 |

   b. Distribute 90µL each to a PCR strip (12x) and use a P20 multichannel to distribute blocking oligo and mix.
17. Using a whole new P10 pipette tip box, add 10µL blocking oligo 3 mixture to each well. mix, and seal plate.
18. Shake at 300rpm for 30min at 25ºC.

19. Pool samples to a trough and add 70µL 7.5% BSA to the solution and mix well.
    c. Can filter while transferring using a FlowMi 40µm filter to remove multiplets.

20. Transfer to 4x lo-bind 2mL tubes, spin down at 500xg for 5 min at room temp in a swing bucket centrifuge.
21. Remove supernatant and add ***gently*** 1mL NIB-RI to pellet **without** resuspending.

22. Spin down at 500xg for 5 min at room temp, remove supernatant.
23. Resuspend with 85µL NIB-RI: start with 40µL, pool all tubes, measure volume and top up to 85 µL.
24. Prepare Ligation mix:

| **Ligation buffer** | **Vol (µL)** |
|---|---|
| Ultrapure water | 251.8 |
| T4 ligase buffer (10x) | 40 |
| SUPERase RI | 1 |
| Enzymatics RI | 3.2 |
| 10% NP-40 | 4 |
| T4 ligase | 20 |
| Total | 320 |

25. Mix 80µL sample to 320µL ligation mix.
26. Distribute 50µL to 8 PCR tubes.
27. Shake at 300rpm for 30 min at 25ºC.

28. Pool samples, spin down at 500xg for 5 min at room temp.
29. Remove supernatant and wash pellet with 1mL NIB-RI.
30. Spin down at 500xg for 5 min at room temp.
31. Resuspend ***thoroughly*** (10x pipetting) with **200µL NIB-RI and count nuclei (post-ligation).
    d. Volume of final resuspension depends on expected nuclei recovery. Generally, can assume between 100-200k but will change depending on cell type and the number of cells at the start of split and pool.
    e. **400µL assumes 200k final recovery. Adjust appropriately for desired number of cells per library.

**<u>Reverse crosslinking & pull down (2:45 hrs)</u>**

1. Based on desired sub-library size (# cells), dilute the cells appropriately for each 50µL reaction.
    a. Range: 2,000-20,000 cells / 50µL.
    b. Each sub-library is processed independently.
    c. N = number of sublibraries
2. Prepare reverse crosslinking master mix as follows.

| **RC MM** | **Vol for 1 sublib (µL)** |
|---|---|
| 2x RCB Buffer | 50 |
| proteinase K (20mg/mL) | 2 |
| SUPERase RI | 1 |
| Total | 53 |

3. For each tube of 50µL sample, add 53 µL of RC MM.

4. Incubate at 55˚C for 1h.
    a. Do not incubate longer, decreases yield

5. Recommended stopping point, store product at -80˚C for few weeks

6. Thaw libraries, if necessary, PMSF/IPA, equilibrate B&W buffers to room temp.
7. After thaw, stop proteinase K by adding 5µL 100mM PMSF/IPA (don't freeze thaw) and incubate at room temp for 10 min.
8. Prepare following buffers (N = number of sublibraries):

| **1x B&W-T/RI** | **Vol for 1 sublib (µL)** |
|---|---|
| 1x B&W-T | 400 |

| | |
|---|---|
| SUPERase RI | 4 |
| Total | 404 |

| 2x B&W/RI | Vol for 1 sublib (µL) |
|---|---|
| 2x BW | 100 |
| SUPERase RI | 2 |
| Total | 102 |

| 1x STE/RI | Vol for 1 sublib (µL) |
|---|---|
| STE | 200 |
| SUPERase RI | 1 |
| Total | 201 |

9. For n number of sublibraries, mix 11*n µL Streptavidin Dynabeads with 110*n µL 1x B&W-T and place it on a magnetic rack
10. Remove supernatant, wash ***twice*** with 110*n µL 1x B&W-T (no RNase inhibitors).
11. Remove supernatant, wash ***once*** with 110*n µL 1x B&W-T/RI (w/ RNase inhibitors).
12. Resuspend beads in 110*n µL **2x** B&W/RI.
13. Add 100µL beads to each sample and rotate over head at 10 rpm for 60 min at room temp. Ensure lids are closed.
    a. Prepare tubes for ATAC supernatant and remember that you need the SUPERNATANT.
    b. Thaw ATAC PCR reagents, thaw reagents + make TSO mix (need it immediately in Template Switching step).
14. Put on a magnetic rack, transfer supernatant containing chromatin fragments to a new tube for library preparation.
    a. Supernatant = ATAC (stable for few hours at room temp, ideally on ice, also can be processed in parallel)
        i. **DO NOT THROWAWAY SUPERNATANT**
        ii. **Refer to ATAC library preparation section for next steps with supernatant**
    b. Beads = RNA
15. Wash beads with 100µL B&W-T/RI ***three times***.
16. Put sample on the magnetic rack, add 100µL 1x STE/RI **without resuspending beads**.

**RNA library preparation – template switching (2.5hrs)**

1. Prepare template switch mix, use wide bore tips for Ficoll and 50% PEG:

| Template switch mix | Vol for 1 sublib (µL) |
|---|---|
| Ultrapure water | 1.25 |
| 5x RT buffer | 10 |
| 50% PEG 6000 | 15 |
| Ficoll PM-400 (20%) | 10 |
| 10mM dNTPs (each) | 5 |
| NxGen RNAse inhibitor | 5 |
| TSO 100µM | 1.25 |
| Maxima H Minus RT (add right before rxn) | 2.5 |
| Total | 50 |

2. Remove all supernatant (100µL 1x STE/RI) and resuspend beads in 50µL template switch mix
    a. Avoid letting beads dry. Do so by first quickly adding the template switch mix to each sample to coat the bead before going back and resuspending all samples
    b. Ensure the beads are thoroughly resuspended. Can be helpful to start with a multichannel at first, and then also pipette each sample individually. Bubbles are inevitable at this stage and during TSO, so the main focus is to get everything resuspended.
3. Rotate samples at room temp for 30 min at 10 rpm.
4. Set thermomixer to 42˚C before step 3 ends. Incubate samples at 42˚C for 90 min shaking at 700rpm.
    a. Resuspend beads every 30 min by pipetting up and down.

**RNA library preparation – on bead cDNA amplification/clean up (2.5hrs)**

1. Prepare cDNA amplification PCR mastermix

| cDNA PCR mastermix | Vol for 1 sublib (µL) |
|---|---|
| Ultrapure water | 25.74 |
| KAPA HiFi 2x MM | 27.5 |
| RNA PCR primer 25µM | 0.88 |
| P7 primer 25µM | 0.88 |
| Total | 55 |

2. Add 100µL of STE to each sample and mix to make viscous solution more pipettable.
3. Place on magnet and discard supernatant.
4. Wash bound beads with 200µL STE (**do not resuspend and do not disturb pellet**) and let sit for a bit.
5. Remove STE and add 55µL PCR mix to beads.
6. Set up thermocycler as below:

| Temp | Time | |
|---|---|---|
| 95˚C | 3 min | |
| 98˚C | 20s | 5 cycles |
| 65˚C | 45s | |
| 72˚C | 3 min | |
| 4˚C | Hold | |
| Total | 30 min | |

7. For each sublibrary, check by qPCR for additional cycles as follows:
   a. Prepare qPCR mastermix:

| qPCR mastermix for RNA | Vol for 1 sublib (µL) |
|---|---|
| KAPA HIFI 2x MM | 3.75 |
| RNA PCR primer 25µM | 0.12 |
| P7 primer 25µM | 0.12 |
| EVAgreen 20x | 0.5 |
| Ultrapure water | 3.01 |
| Total | 7.5 |

   b. Put samples on magnet, mix 2.5µL sample supernatant with 7.5µL qPCR mix.
   c. Run thermocycler as below for qPCR:

| Temp | Time | |
|---|---|---|
| 95˚C | 3 min | |
| 98˚C | 20s | 20 cycles for qPCR / 0.33Ct for add PCR |
| 65˚C | 20s | |
| 72˚C | 3 min | |
| 4˚C | Hold | |

8. Perform additional cycles based on 0.33Ct of maximum saturation.
   a. Expect 6-7 addl cycles for 2k cells, ~4 for 10k cells.
9. Magnetically separate out the Dynabeads, purify the supernatant of each sample with 40µL SPRI beads (0.8x) in new tubes, making sure no Dynabeads come through.
   a. Make fresh 80% EtOH (400µL per sublib).
   b. Add 40µL SPRI beads to each 50µL reaction.
   c. Incubate for 5min at room temp.
   d. Put on magnet and remove supernatant.
   e. Wash with 200µL 80% EtOH directly on magnet, waiting 30s each. Repeat for a total of two washes.
   f. Carefully remove all supernatant and air dry the beads until they just turn matte.

g. Remove from magnet, thoroughly resuspend in 15μL EB to elute. Let sit for 2min before briefly centrifuging and putting back on magnet. Transfer supernatant to a new tube.

10. cDNA can be stored at -20˚C indefinitely until tagmentation.

**RNA library preparation – cDNA Tagmentation**

1. Quantify cDNA concentration by Qubit using 1μL cDNA.
    a. If necessary, dilute 50ng cDNA to 5 ng/μL in water.
    b. Can get away with 20ng only if necessary.
2. Start thermocyler protocol to ensure the machine is at 55˚C hold before samples go in.
3. Mix following together:

| **cDNA tagmentation** | **Vol for 1 sublib (μL)** |
|---|---|
| 2x TD buffer | 25 |
| SHARE-cDNA Tn5 WJG Mar 2023 | 1 |
| Subtotal | 26 |
| cDNA (5ng/μL) | 10 |
| Ultrapure water | 14 |
| Total | 50 |

4. Incubate at 55˚C on thermocycler for 5 min.
5. Purify library with Zymo DNA Clean and Concentrate (250μL binding buffer). Try to get the samples to binding buffer asap to minimize over tagmentation.
    a. Elute twice with 11μL EB each time (22μL total).
6. Prepare post-tagmentation PCR mix, record the i5 index used for each sublib.

| **Post tagmentation PCR mix** | **Vol for 1 sublib (μL)** |
|---|---|
| NEBnext 2x MM | 25 |
| P7 primer 25μM | 1 |
| Ultrapure water | 3 |
| Subtotal | 29 |
| Eluted sample | 20 |
| i5 index primer (Ad 1.xx) 25μM | 1 |
| Total | 50 |

7. Set up thermocycler for the sample as below:

| **Temp** | **Time** | |
|---|---|---|
| 72˚C | 5 min | |
| 98˚C | 30s | |
| 98˚C | 10s | 10 cycles (11 cycles if cDNA input is ~20ng) |
| 65˚C | 30s | |
| 72˚C | 60s | |
| 4˚C | Hold | |
| Total | 20 min | |

8. Purify each sample with 40μL SPRI beads (0.8x) and elute library to 15μL EB.
    a. If there is a significant amount of large fragments present in tapestation after this step, can redo purification with a double sided SPRI (0.6x-0.8x) or gel extraction.
9. Libraries can be stored at -20˚C indefinitely.

**ATAC library preparation**

1. Clean the supernatant with Zymo DNA clean and concentrate kit by adding 5x DNA binding buffer (200µL rxn = 1mL of binding buffer, spin twice because the column only holds 0.7mL).
    a. Elute in 11µL EB then add 11µL EB again for a total elution volume of 22µL.
2. Prepare ATAC PCR mix, record the i5 index used for each sublib

| ATAC PCR mastermix | Vol for 1 sublib (µL) |
|---|---|
| NEBnext 2x MM | 25 |
| P7 primer 25µM | 1 |
| Ultrapure water | 3 |
| Subtotal | 29 |
| Eluted sample | 20 |
| i5 index primer (Ad 1.xx) 25µM | 1 |
| Total | 50 |

3. Mix and set up thermocycler as shown below:

| Temp | Time | |
|---|---|---|
| 72˚C | 5 min | |
| 98˚C | 30s | |
| 98˚C | 10s | 5 cycles |
| 65˚C | 30s | |
| 72˚C | 30s | |
| 4˚C | Hold | |
| Total | 15 min | |

4. Check by qPCR as usual for ATAC:
    a. 5µL sample + 10µL qPCR mix:

| qPCR mastermix for ATAC | Vol for 1 sublib (µL) |
|---|---|
| NEBnext 2x MM | 5 |
| Ad 1.1 25µM | 0.2 |
| P7 primer 25µM | 0.2 |
| 10x SYBRgreen | 0.9 |
| Ultrapure water | 3.7 |
| Total | 10 |

        i. Dilute 100x SYBR to 10x before use fresh

    b. Run qPCR as below

| Temp | Time | |
|---|---|---|
| 98˚C | 30s | |
| 98˚C | 10s | 20 cycles for qPCR / 0.33Ct for addl PCR |
| 65˚C | 30s | |
| 72˚C | 30s | |
| 4˚C | Hold | |

    c. Perform additional cycles based on 1/3 of the plateau of maximum fluorescence from qPCR (0.33Ct) (usually ~8 additional cycles for 2k cells).
5. Clean up with Zymo DNA clean and concentrate (225µL binding buffer if qPCR) and elute twice 8µL each for ~15µL total.
6. Libraries can be stored at -20˚C indefinitely.

**Library quantification and sequencing**

1. Measure concentration of libraries using Qubit
2. Run ATAC library on D5000 tapestation (10μL buffer + 1μL sample). Expected ATAC tapestation trace:

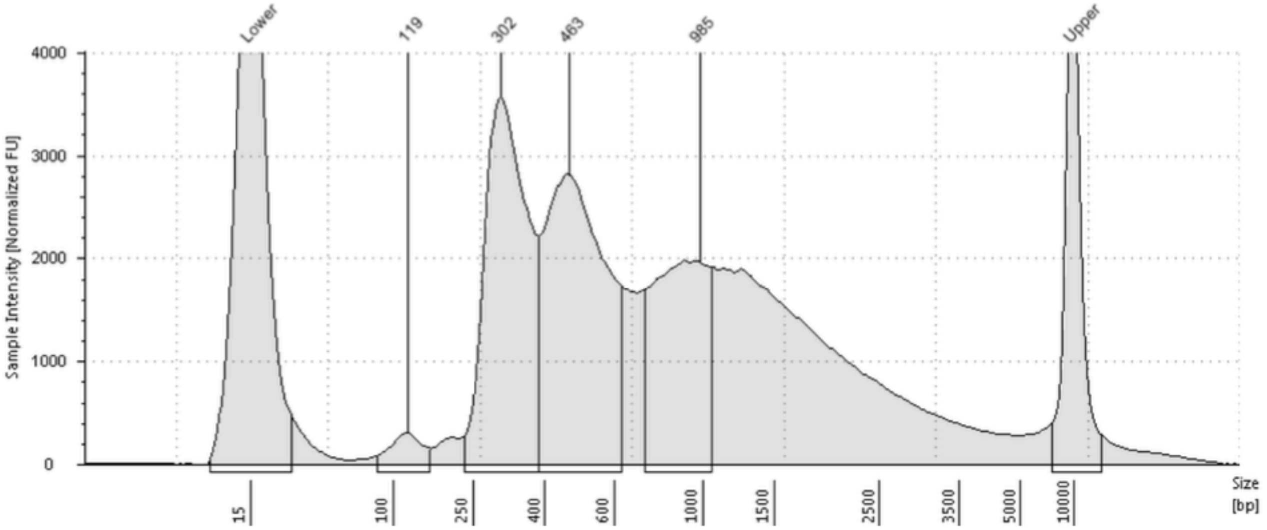


3. Run RNA library on D1000 tapestation (3μL buffer + 1μL sample). Expected RNA tapestation trace:

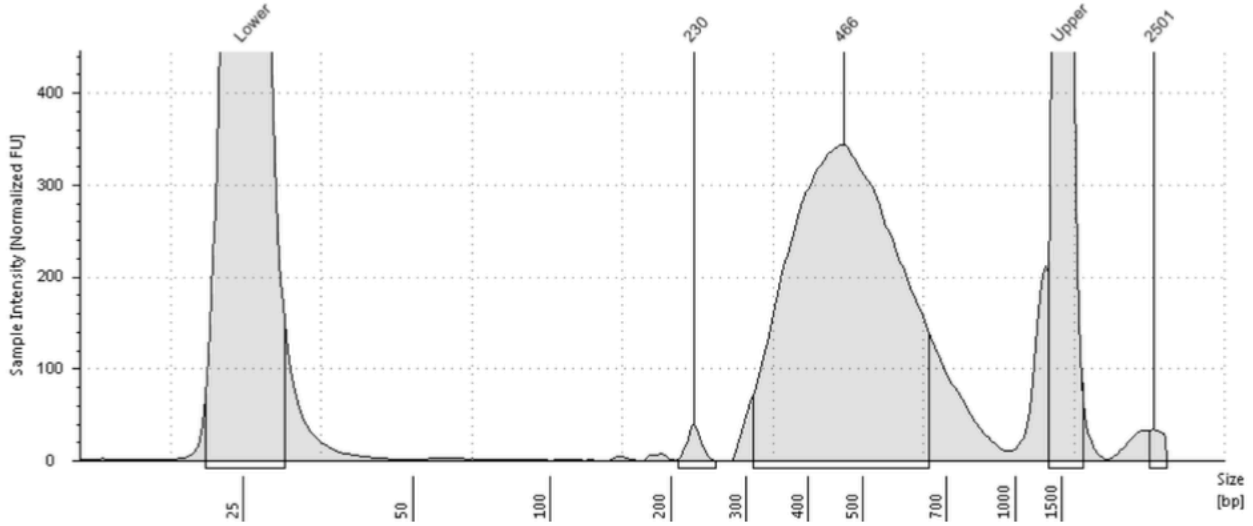


4. Sequencing configuration

If sequencing ATAC only or ATAC+RNA together:

| # cycles in seq kit | R1 | I1 | I2 | R2 |
|---|---|---|---|---|
| 150 cycle | 30 | 99 | 8 | 30 |
| 300 cycle | 96 | 99 | 8 | 96 |

If sequencing RNA only:

| # cycles in seq kit | R1 | I1 | I2 | R2 |
|---|---|---|---|---|
| 150 cycle | 50 | 99 | 8 | 10 |
| 300 cycle | 162 | 99 | 8 | 30 |

The general sequencing depth recommendation is 50,000 reads per cell per modality (ATAC or RNA). For example, for a cell sublibrary of 20,000 cells, use 2 billion reads for both ATAC and RNA sublibraries.

**Materials and equipment**

*The volumes shown in the table below does not include amount needed for pre-made buffers

| category | material | vendor | catalog # | vol needed for this exp (µL) | storage location | Notes (including vol per sample/rxn in µL) |
|---|---|---|---|---|---|---|
| Enzymes | SHARE-ATAC Tn5 | In house | WJG Mar 2023 | 312.5 | -20C | 2.5/ATAC rxn |
| | SHARE-cDNA Tn5 | In house | WJG Mar 2023 | 17.6 | -20C | 1/sublib |
| | Maxima H Minus RT and buffer | Thermo | EP0753 | 319 | -20C | 5/RT rxn+2.5/sublib |
| | T4 DNA Ligase | NEB | M0202L | 20 | -20C | 20 fixed |
| | 20mg/mL Proteinase K | NEB | P8107S | 35.2 | -20C | 2/sublib |
| | Kapa Hifi Hotstart Ready Mix | Roche | 7958935001 | 550 | -20C | 31.25/sublib |
| | NEBNext 2x MM | NEB | M0541L | 968 | -20C | 55/sublib |
| RNase inhibitors | Enzymatics RI | Enzymatics | Y9240L | 314.1758 | -20C | 17.52/sample + 0.8625/ ATAC rxn + 0.31/RT rxn + 62.9 fixed |
| | Superase RI | Thermo | AM2696 | 261.01 | -20C | 8.27/sample + 0.63/RT rxn + 8/sublib + 30.98 fixed |
| | NxGen RNAse inhibitor | Lucigen | 30281-1 | 88 | -20C | 5/sublib |
| Beads | Dynabeads MyOne Streptavidin C1 | Thermo | 65002 | 176 | 4C | 10/sublib |
| | SPRI beads | Beckman | B23318 | 1408 | 25C | 80/sublib |
| Chemicals | 7.5% BSA | Thermo | 15260037 | 1275.36 | 4C | 182.63/sample, 70 fixed |
| | 10% Tween-20 | Biorad | 1662404 | 141.4 | 25C | 11/sample, 0.5/ATAC rxn |
| | 10% NP-40 | Thermo | 85124 | 76.1 | 25C | 1/sample, 69.5 fixed |
| | 1% Digitonin | Promega | G9441 | 75.4 | -20C | Dilute to 1%, 1/sample +0.5/ATAC rxn |
| | 1M DTT | Sigma | 646563 | 7.3 | -20C | 1.1/sample |
| | 16% Formaldehyde (FA) | Thermo | 28906 | 66 | 25C | 10/sample |
| | 100% DMF | Thermo | 20673 | 1100 | 25C | 8/ATAC rxn |
| | PIC (protease inhibitor cocktail) | Sigma | P8340 | 27.5 | -20C | 0.2/ATAC rxn |
| | dNTPs (10mM each) | NEB | N0447L | 225.5 | -20C | 2.5/RT rxn, 5/sublib |
| | 50% PEG 6000 | | | 1089 | 25C | 15/RT rxn, 15/sublib |
| | Buffer EB | Qiagen | 19086 | lots | 25C | |
| | 10x T4 DNA Ligase Buffer | NEB | B0202S | 1002.4 | -20C | 1002.4 fixed |
| | PMSF 100mM | Thermo | 36978 | 88 | -20C | 5/sublib |
| | Ficoll PM-400 (20%) | Sigma | F5415 | 176 | 4C | 10/sublib |
| | 10x SYBRgreen | Thermo | S7563 | 15.8 | -20C | Dilute stock to 10x, 0.9/sublib |
| | 20x EVAgreen | Biotium | 31000 | 8.8 | -20C | 0.5/sublib |
| | 5M NaCl | Any | | | 25C | |
| | 1M MgCl2 | | | | 25C | |
| | 1M Tris HCl pH 7.5 | | | | 25C | |
| | 1M Tris HCl pH 8.0 | | | | 25C | |
| | 2.5M Glycine | | | | 25C | |
| | 0.2M Tris-acetate | | | | 4C | |
| | 5M K-acetate | | | | 25C | |
| | 1M Mg-acetate | | | | 25C | |
| | 1x PBS | | | | 25C | |
| | 20% SDS | | | | 25C | |
| | 0.5M EDTA | | | | 25C | |
| Oligos | RT primer 100µM | See Sai Ma et al Cell 2020 Supp Table for all oligo sequences | | 275 | -20C | 5/RT rxn |
| | RNA PCR Primer 25µM | | | 17.6 | -20C | 1/sublib |
| | P7 primer 25µM | | | 56.3 | -20C | 3.2/sublib |
| | Nextera i5 index primers Ad 1.xx 25µM | | | 32 x 1 | -20C | 1/atac or rna sublib, all unique |
| | TSO 100µM | | | 22 | -20C | 1.25/sublib |
| | Round 1 blocking 1mM | | | 25.3 | -20C | 25.3 fixed |
| | Round 2 blocking 1mM | | | 30.4 | -20C | 30.4 fixed |
| | Round 3 blocking 1mM | | | 26.5 | -20C | 26.5 fixed |
| Kits | DNA clean and concentrate kit | Zymo | D4014 | 48 assays | 25C | |
| | Qubit dsDNA HS assay | Thermo | Q32854 | 32 assays | 25C /4C | |
| | D5000 ScreenTape | Agilent | 5067-5588 | 17 assays | 4C | |
| | D5000 Reagents | Agilent | 5067-5589 | 17 assays | 4C | |
| | D1000 ScreenTape | Agilent | 5067-5582 | 17 assays | 4C | |
| | D1000 Reagents | Agilent | 5067-5583 | 17 assays | 4C | |
| Equipment | Centrifuge | Any | | | | |
| | FlowMi 40um filter | | | | | |
| | Wide bore p200 and p1000 tips | | | | | |
| | Tube rotator | | | | | |
