## Supplementary Note 2 for "Dissecting regulatory syntax in human development with scalable multiomics and deep learning"

#### Supplementary Note 2: Quality control per organ.

**Quality control for Adrenal (AG), Brain (BR), Eye (EY), Heart (HT), Liver (LI), Lung (LU), Muscle (MU), Skin (SK), Spleen (SP), StomachEsophagus (ST), Thymus (TM), and Thyroid (TR). a)** UMAP of organ and gestational age per cluster in post-conception weeks. **b)** RNA marker gene dot plot for each cell cluster. **c)** RNA QC metrics (# UMIs, # Genes) and ATAC QC metrics (# Fragments, TSS enrichment ratio) for each cluster. **d)** ATAC marker peaks (left) and motif enrichment within marker peaks (right), columns sorted by hierarchical clustering of motif enrichments. **e)** Top TF motifs with the highest variability across clusters from chromVAR deviation calculations. **f)** ChromBPNet per-cluster summary statistics (total # fragments and # cells per cluster) and QC metrics (Pearson R correlation between observed and predicted log counts in peaks, and Jensen-Shannon distance between predicted profiles).

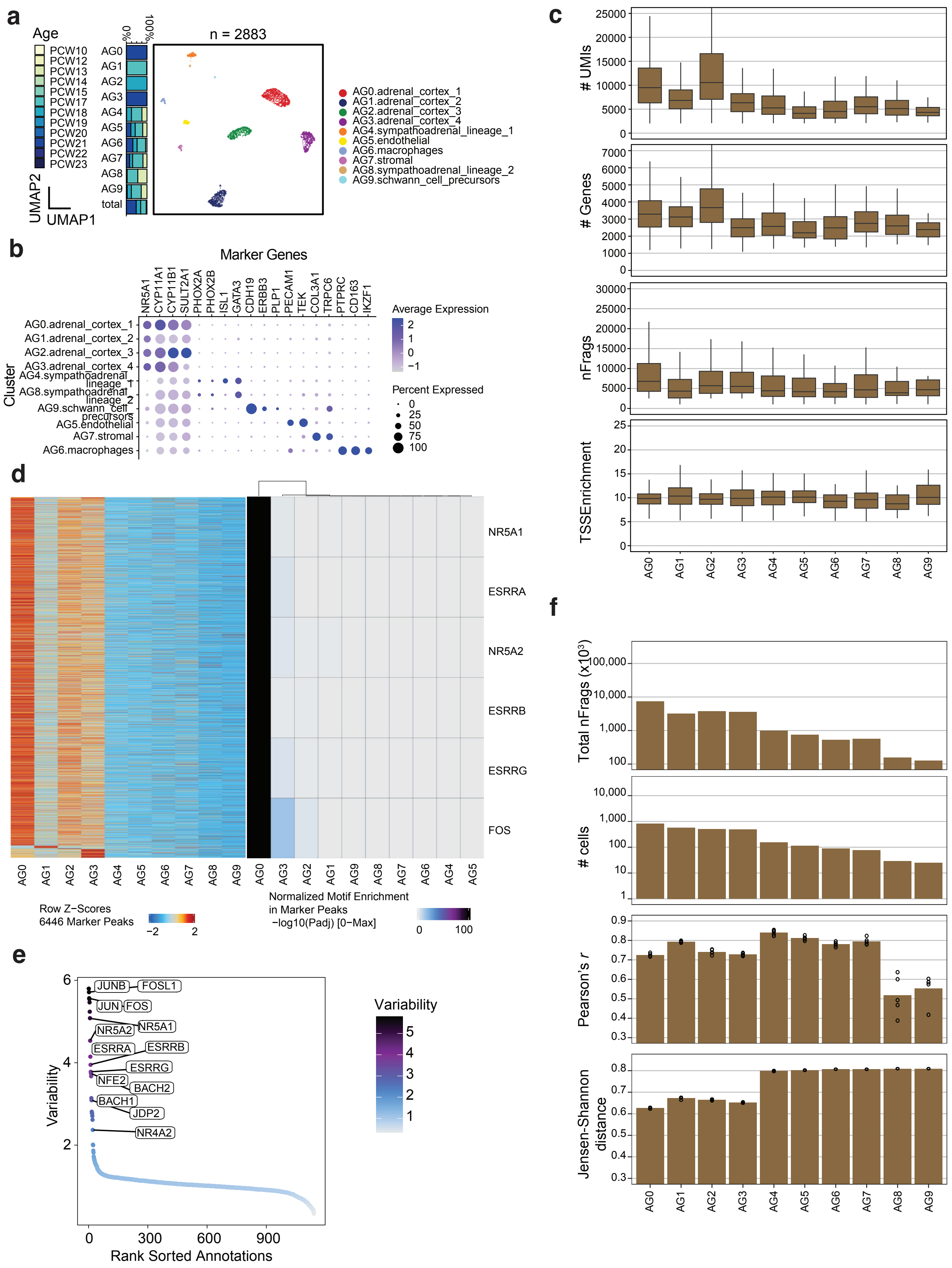
a
Age
PCW10
PCW12
PCW13
PCW14
PCW15
PCW17
PCW18
PCW19
PCW20
PCW21
PCW22
PCW23
0%
100%
AG0
AG1
AG2
AG3
AG4
AG5
AG6
AG7
AG8
AG9
total
UMAP2
UMAP1
n = 2883
AG0.adrenal_cortex_1
AG1.adrenal_cortex_2
AG2.adrenal_cortex_3
AG3.adrenal_cortex_4
AG4.sympathoadrenal_lineage_1
AG5.endothelial
AG6.macrophages
AG7.stromal
AG8.sympathoadrenal_lineage_2
AG9.schwann_cell_precursors
b
Marker Genes
NR5A1
CYP11A1
CYP11B1
SULT2A1
PHOX2A
PHOX2B
ISL1
GATA3
CDH19
ERBB3
PLP1
PECAM1
TEK
COL3A1
TRPC6
PTPRC
CD163
IKZF1
Cluster
AG0.adrenal_cortex_1
AG1.adrenal_cortex_2
AG2.adrenal_cortex_3
AG3.adrenal_cortex_4
AG4.sympathoadrenal_lineage_1
AG8.sympathoadrenal_lineage_2
AG9.schwann_cell_precursors
AG5.endothelial
AG7.stromal
AG6.macrophages
Average Expression
2
1
0
−1
Percent Expressed
0
25
50
75
100
c
### UMIs
25000
20000
15000
10000
5000
0
### Genes
7000
6000
5000
4000
3000
2000
1000
0
nFrags
30000
25000
20000
15000
10000
5000
0
TSSEnrichment
25
20
15
10
5
0
AG0
AG1
AG2
AG3
AG4
AG5
AG6
AG7
AG8
AG9
d
NR5A1
ESRRA
NR5A2
ESRRB
ESRRG
FOS
AG0
AG1
AG2
AG3
AG4
AG5
AG6
AG7
AG8
AG9
AG0
AG3
AG2
AG1
AG9
AG8
AG7
AG6
AG4
AG5
Row Z-Scores
6446 Marker Peaks
−2
2
Normalized Motif Enrichment
in Marker Peaks
−log10(Padj) [0−Max]
0
100
e
Variability
6
4
2
JUNB
FOSL1
JUN
FOS
NR5A2
NR5A1
ESRRA
ESRRB
ESRRG
NFE2
BACH2
BACH1
JDP2
NR4A2
0
300
600
900
Rank Sorted Annotations
Variability
5
4
3
2
1
f
Total nFrags (x10^3)
100,000
10,000
1,000
100
### cells
10,000
1,000
100
10
1
Pearson's r
0.9
0.8
0.7
0.6
0.5
0.4
0.3
Jensen-Shannon distance
0.8
0.7
0.6
0.5
0.4
0.3
AG0
AG1
AG2
AG3
AG4
AG5
AG6
AG7
AG8
AG9

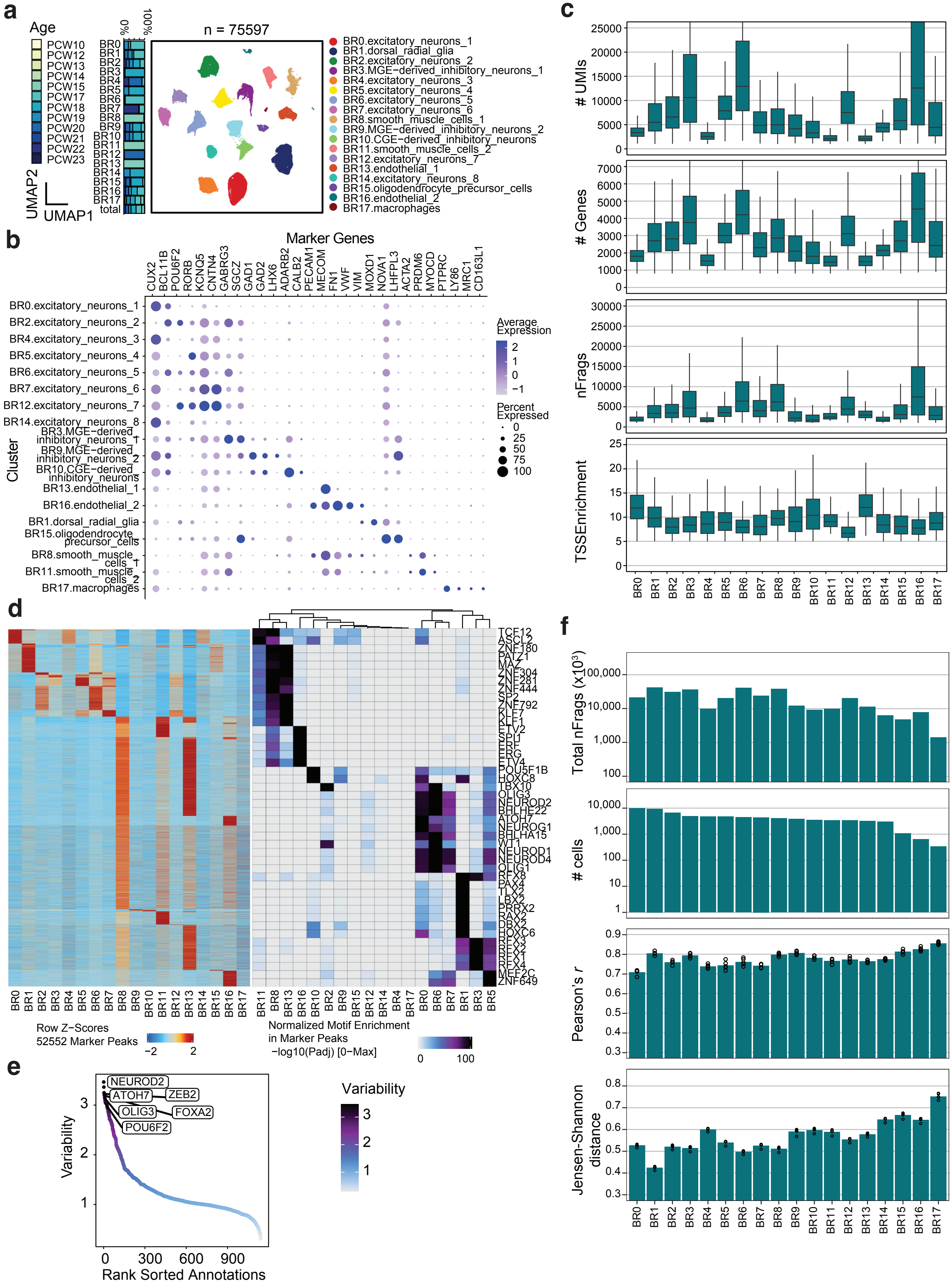
a
Age
PCW10
PCW12
PCW13
PCW14
PCW15
PCW17
PCW18
PCW19
PCW20
PCW21
PCW22
PCW23
0%
100%
UMAP2
UMAP1
total
n = 75597
BR0.excitatory_neurons_1
BR1.dorsal_radial_glia
BR2.excitatory_neurons_2
BR3.MGE−derived_inhibitory_neurons_1
BR4.excitatory_neurons_3
BR5.excitatory_neurons_4
BR6.excitatory_neurons_5
BR7.excitatory_neurons_6
BR8.smooth_muscle_cells_1
BR9.MGE−derived_inhibitory_neurons_2
BR10.CGE−derived_inhibitory_neurons
BR11.smooth_muscle_cells_2
BR12.excitatory_neurons_7
BR13.endothelial_1
BR14.excitatory_neurons_8
BR15.oligodendrocyte_precursor_cells
BR16.endothelial_2
BR17.macrophages
b
Marker Genes
CUX2 BCL11B POU6F2 RORB KCNQ5 CNTN4 GABRG3 SGCZ GAD1 GAD2 LHX6 ADARB2 CALB2 PECAM1 MECOM FN1 VWF VIM MOXD1 NOVA1 LHFPL3 ACTA2 PRDM6 MYOCD PTPRC LY86 MRC1 CD163L1
Cluster
Average Expression
Percent Expressed
c
### UMIs
### Genes
nFrags
TSSEnrichment
d
Row Z−Scores
52552 Marker Peaks
Normalized Motif Enrichment in Marker Peaks −log10(Padj) [0−Max]
TCF12 ASCL2 ZNF180 PATZ1 MAZ ZNF304 ZNF281 ZNF444 SP2 ZNF792 KLF7 KLF1 ETV2 SPI1 ERF ERG ETV4 POU5F1B HOXC8 TBX10 OLIG3 NEUROD2 BHLHE22 ATOH7 NEUROG1 BHLHA15 WT1 NEUROD1 NEUROD4 OLIG1 RFX8 PAX4 TLX2 LBX2 PRRX2 RAX2 DBX2 HOXC6 RFX3 RFX2 RFX1 RFX4 MEF2C ZNF649
e
Variability
Rank Sorted Annotations
NEUROD2
ATOH7
ZEB2
OLIG3
FOXA2
POU6F2
f
Total nFrags (x10³)
### cells
Pearson's r
Jensen-Shannon distance

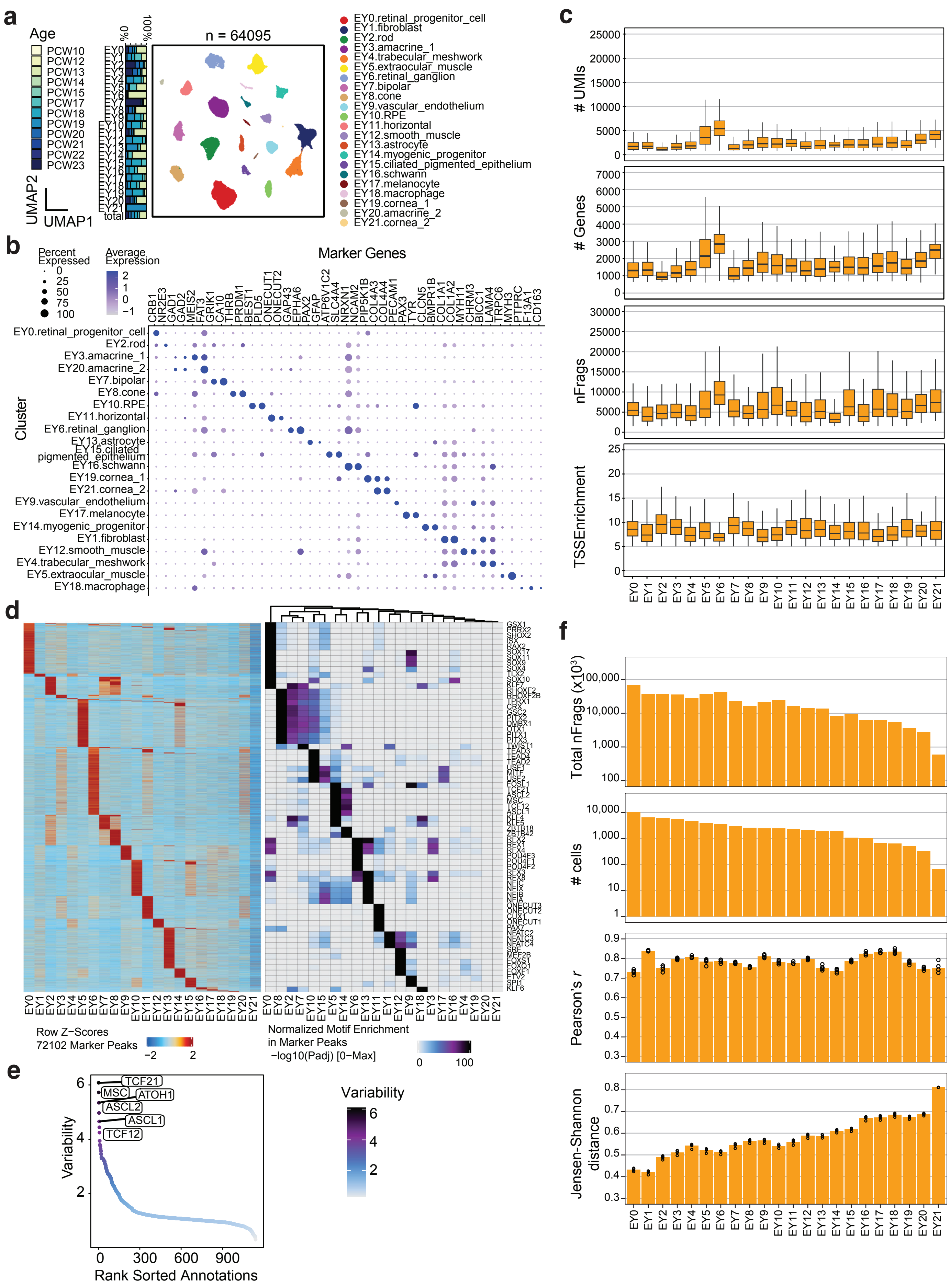
a
Age
PCW10
PCW12
PCW13
PCW14
PCW15
PCW17
PCW18
PCW19
PCW20
PCW21
PCW22
PCW23
0%
100%
n = 64095
UMAP2
UMAP1
EY0.retinal_progenitor_cell
EY1.fibroblast
EY2.rod
EY3.amacrine_1
EY4.trabecular_meshwork
EY5.extraocular_muscle
EY6.retinal_ganglion
EY7.bipolar
EY8.cone
EY9.vascular_endothelium
EY10.RPE
EY11.horizontal
EY12.smooth_muscle
EY13.astrocyte
EY14.myogenic_progenitor
EY15.ciliated_pigmented_epithelium
EY16.schwann
EY17.melanocyte
EY18.macrophage
EY19.cornea_1
EY20.amacrine_2
EY21.cornea_2
b
Percent Expressed
Average Expression
Marker Genes
Cluster
c
### UMIs
### Genes
nFrags
TSSEnrichment
d
Row Z-Scores
72102 Marker Peaks
Normalized Motif Enrichment in Marker Peaks
−log10(Padj) [0−Max]
e
Variability
Rank Sorted Annotations
TCF21
MSC
ATOH1
ASCL2
ASCL1
TCF12
f
Total nFrags (x10³)
### cells
Pearson's r
Jensen-Shannon distance

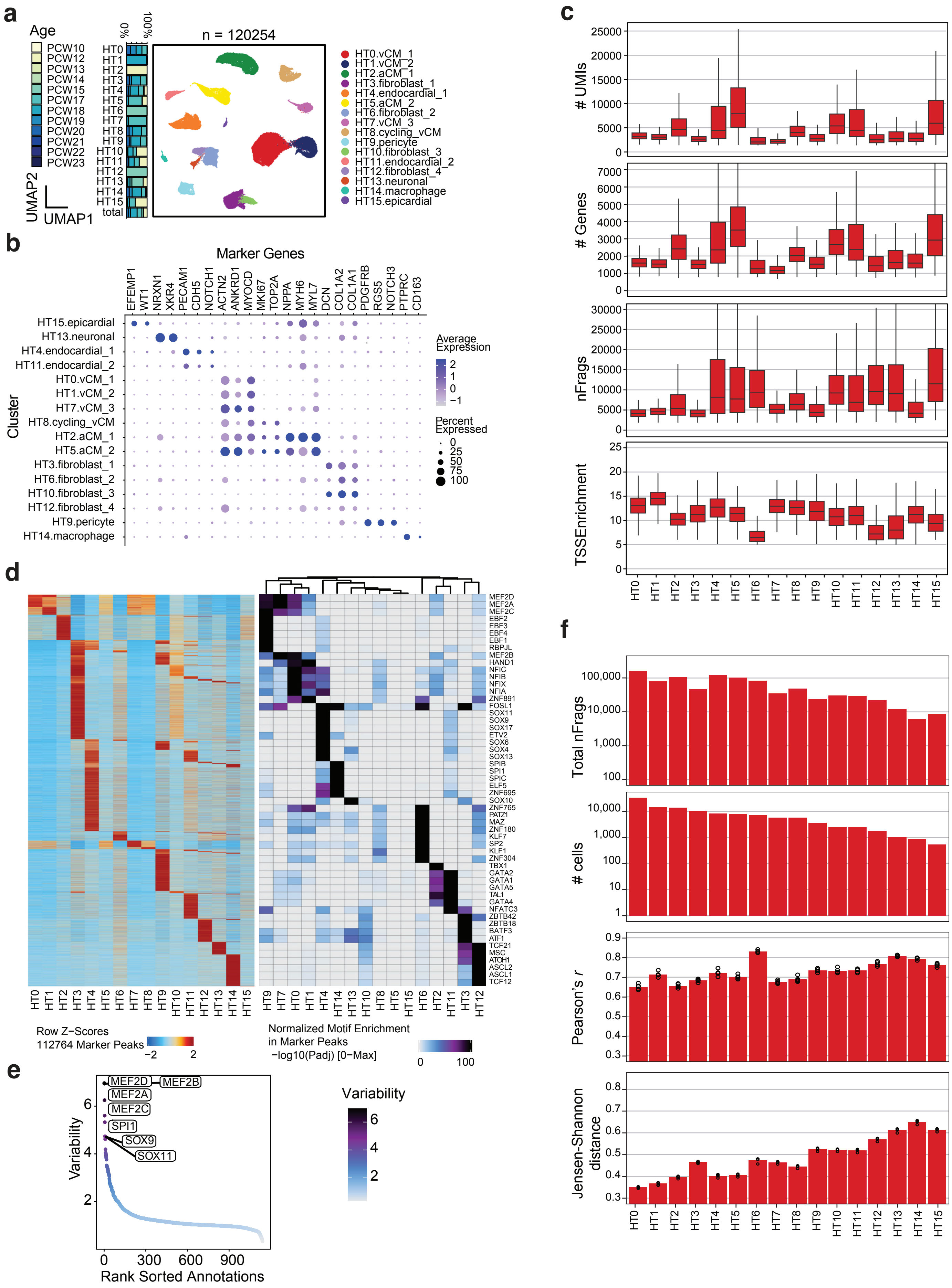
a
Age
PCW10
PCW12
PCW13
PCW14
PCW15
PCW17
PCW18
PCW19
PCW20
PCW21
PCW22
PCW23
0%
100%
n = 120254
UMAP2
UMAP1
HT0.vCM_1
HT1.vCM_2
HT2.aCM_1
HT3.fibroblast_1
HT4.endocardial_1
HT5.aCM_2
HT6.fibroblast_2
HT7.vCM_3
HT8.cycling_vCM
HT9.pericyte
HT10.fibroblast_3
HT11.endocardial_2
HT12.fibroblast_4
HT13.neuronal
HT14.macrophage
HT15.epicardial
total
b
Marker Genes
EFEMP1
WT1
NRXN1
XKR4
PECAM1
CDH5
NOTCH1
ACTN2
ANKRD1
MYOCD
MKI67
TOP2A
NPPA
MYH6
MYL7
DCN
COL1A2
COL1A1
PDGFRB
RGS5
NOTCH3
PTPRC
CD163
Cluster
Average Expression
Percent Expressed
c
### UMIs
### Genes
nFrags
TSSEnrichment
d
Row Z−Scores
112764 Marker Peaks
Normalized Motif Enrichment in Marker Peaks
−log10(Padj) [0−Max]
MEF2D
MEF2A
MEF2C
EBF2
EBF3
EBF4
EBF1
RBPJL
MEF2B
HAND1
NFIC
NFIB
NFIX
NFIA
ZNF891
FOSL1
SOX11
SOX9
SOX17
ETV2
SOX6
SOX4
SOX13
SPIB
SPI1
SPIC
ELF5
ZNF695
SOX10
ZNF765
PATZ1
MAZ
ZNF180
KLF7
SP2
KLF1
ZNF304
TBX1
GATA2
GATA1
GATA5
TAL1
GATA4
NFATC3
ZBTB42
ZBTB18
BATF3
ATF1
TCF21
MSC
ATOH1
ASCL2
ASCL1
TCF12
e
Variability
Rank Sorted Annotations
MEF2D
MEF2B
MEF2A
MEF2C
SPI1
SOX9
SOX11
f
Total nFrags
### cells
Pearson's r
Jensen-Shannon distance

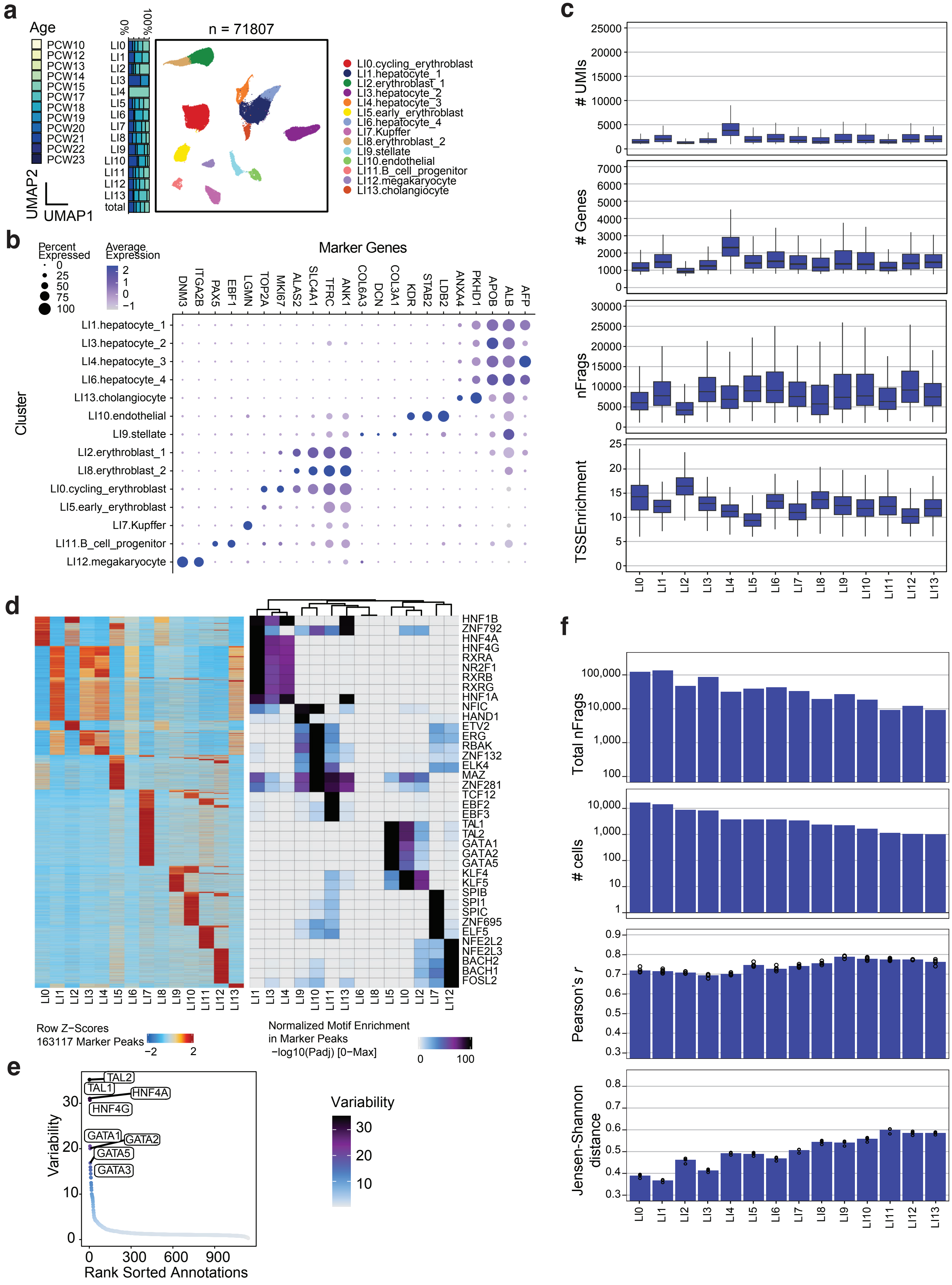
a
Age
PCW10
PCW12
PCW13
PCW14
PCW15
PCW17
PCW18
PCW19
PCW20
PCW21
PCW22
PCW23
UMAP2
UMAP1
0%
100%
LI0
LI1
LI2
LI3
LI4
LI5
LI6
LI7
LI8
LI9
LI10
LI11
LI12
LI13
total
n = 71807
LI0.cycling_erythroblast
LI1.hepatocyte_1
LI2.erythroblast_1
LI3.hepatocyte_2
LI4.hepatocyte_3
LI5.early_erythroblast
LI6.hepatocyte_4
LI7.Kupffer
LI8.erythroblast_2
LI9.stellate
LI10.endothelial
LI11.B_cell_progenitor
LI12.megakaryocyte
LI13.cholangiocyte
b
Percent Expressed
0
25
50
75
100
Average Expression
2
1
0
−1
Marker Genes
DNM3
ITGA2B
PAX5
EBF1
LGMN
TOP2A
MKI67
ALAS2
SLC4A1
TFRC
ANK1
COL6A3
DCN
COL3A1
KDR
STAB2
LDB2
ANXA4
PKHD1
APOB
ALB
AFP
Cluster
LI1.hepatocyte_1
LI3.hepatocyte_2
LI4.hepatocyte_3
LI6.hepatocyte_4
LI13.cholangiocyte
LI10.endothelial
LI9.stellate
LI2.erythroblast_1
LI8.erythroblast_2
LI0.cycling_erythroblast
LI5.early_erythroblast
LI7.Kupffer
LI11.B_cell_progenitor
LI12.megakaryocyte
c
### UMIs
25000
20000
15000
10000
5000
0
### Genes
7000
6000
5000
4000
3000
2000
1000
0
nFrags
30000
25000
20000
15000
10000
5000
0
TSSEnrichment
25
20
15
10
5
0
LI0
LI1
LI2
LI3
LI4
LI5
LI6
LI7
LI8
LI9
LI10
LI11
LI12
LI13
d
HNF1B
ZNF792
HNF4A
HNF4G
RXRA
NR2F1
RXRB
RXRG
HNF1A
NFIC
HAND1
ETV2
ERG
RBAK
ZNF132
ELK4
MAZ
ZNF281
TCF12
EBF2
EBF3
TAL1
TAL2
GATA1
GATA2
GATA5
KLF4
KLF5
SPIB
SPI1
SPIC
ZNF695
ELF5
NFE2L2
NFE2L3
BACH2
BACH1
FOSL2
LI0
LI1
LI2
LI3
LI4
LI5
LI6
LI7
LI8
LI9
LI10
LI11
LI12
LI13
LI1
LI3
LI4
LI9
LI10
LI11
LI13
LI6
LI8
LI5
LI0
LI2
LI7
LI12
Row Z−Scores
163117 Marker Peaks
−2
2
Normalized Motif Enrichment
in Marker Peaks
−log10(Padj) [0−Max]
0
100
e
TAL2
TAL1
HNF4A
HNF4G
GATA1
GATA2
GATA5
GATA3
Variability
30
20
10
0
0
300
600
900
Rank Sorted Annotations
Variability
30
20
10
f
Total nFrags
100,000
10,000
1,000
100
### cells
10,000
1,000
100
10
1
Pearson's r
0.9
0.8
0.7
0.6
0.5
0.4
0.3
Jensen-Shannon distance
0.8
0.7
0.6
0.5
0.4
0.3
LI0
LI1
LI2
LI3
LI4
LI5
LI6
LI7
LI8
LI9
LI10
LI11
LI12
LI13

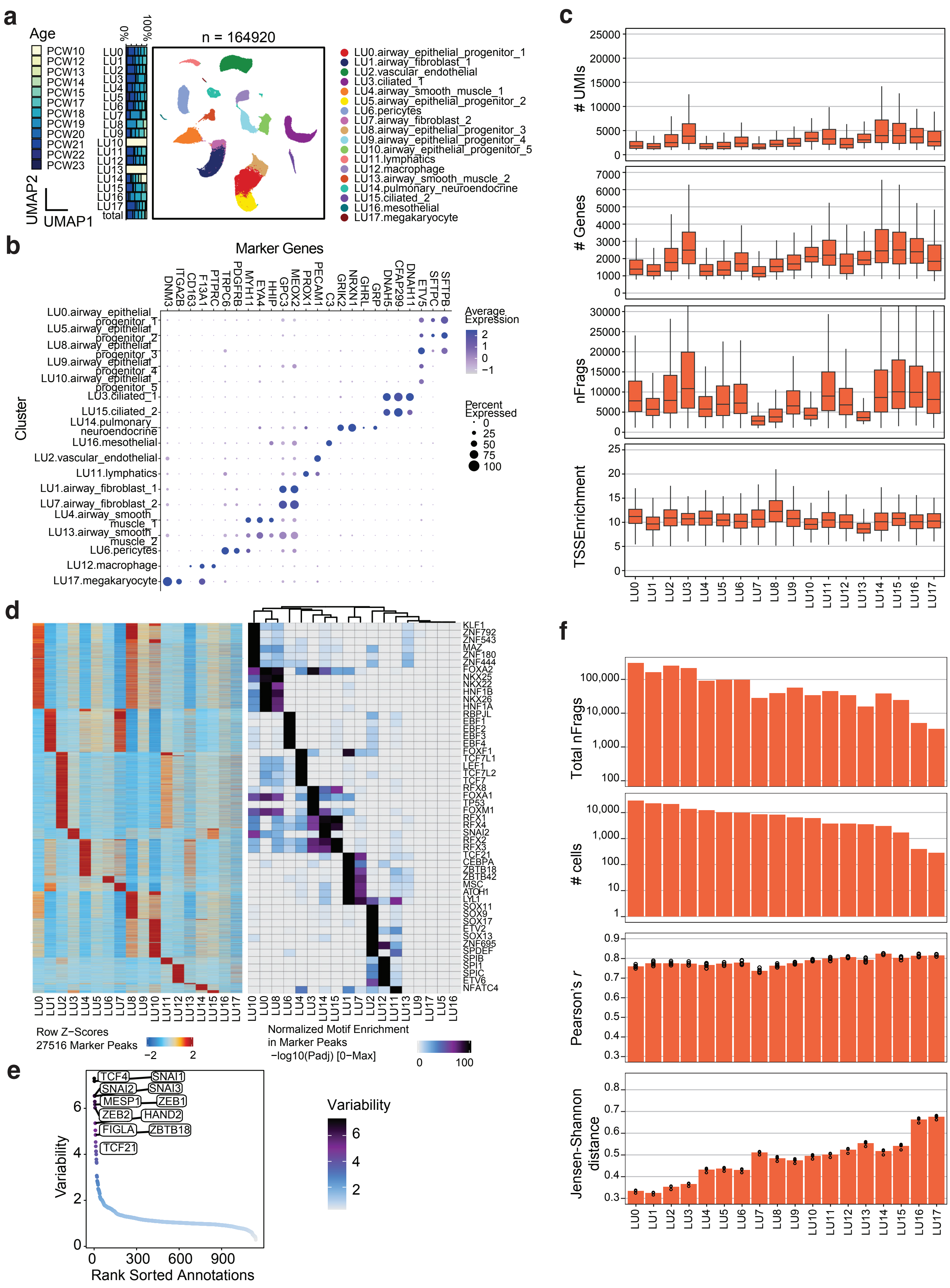
a
Age
PCW10
PCW12
PCW13
PCW14
PCW15
PCW17
PCW18
PCW19
PCW20
PCW21
PCW22
PCW23
0%
100%
LU0
LU1
LU2
LU3
LU4
LU5
LU6
LU7
LU8
LU9
LU10
LU11
LU12
LU13
LU14
LU15
LU16
LU17
total
UMAP2
UMAP1
n = 164920
LU0.airway_epithelial_progenitor_1
LU1.airway_fibroblast_1
LU2.vascular_endothelial
LU3.ciliated_1
LU4.airway_smooth_muscle_1
LU5.airway_epithelial_progenitor_2
LU6.pericytes
LU7.airway_fibroblast_2
LU8.airway_epithelial_progenitor_3
LU9.airway_epithelial_progenitor_4
LU10.airway_epithelial_progenitor_5
LU11.lymphatics
LU12.macrophage
LU13.airway_smooth_muscle_2
LU14.pulmonary_neuroendocrine
LU15.ciliated_2
LU16.mesothelial
LU17.megakaryocyte
b
Marker Genes
DNM3
ITGA2B
CD163
F13A1
PTPRC
TRPC6
PDGFRB
MYH11
EYA4
HHIP
GPC3
MEOX2
PROX1
PECAM1
C3
GRIK2
NRXN1
GHRL
GRP
DNAH5
CFAP299
DNAH11
ETV5
SFTPC
SFTPB
Cluster
LU0.airway_epithelial_progenitor_1
LU5.airway_epithelial_progenitor_2
LU8.airway_epithelial_progenitor_3
LU9.airway_epithelial_progenitor_4
LU10.airway_epithelial_progenitor_5
LU3.ciliated_1
LU15.ciliated_2
LU14.pulmonary_neuroendocrine
LU16.mesothelial
LU2.vascular_endothelial
LU11.lymphatics
LU1.airway_fibroblast_1
LU7.airway_fibroblast_2
LU4.airway_smooth_muscle_1
LU13.airway_smooth_muscle_2
LU6.pericytes
LU12.macrophage
LU17.megakaryocyte
Average Expression
2
1
0
−1
Percent Expressed
0
25
50
75
100
c
### UMIs
25000
20000
15000
10000
5000
0
### Genes
7000
6000
5000
4000
3000
2000
1000
0
nFrags
30000
25000
20000
15000
10000
5000
0
TSSEnrichment
25
20
15
10
5
0
LU0 LU1 LU2 LU3 LU4 LU5 LU6 LU7 LU8 LU9 LU10 LU11 LU12 LU13 LU14 LU15 LU16 LU17
d
KLF1
ZNF792
ZNF543
MAZ
ZNF180
ZNF444
FOXA2
NKX25
NKX22
HNF1B
NKX26
HNF1A
RBPJL
EBF1
EBF2
EBF3
EBF4
FOXF1
TCF7L1
LEF1
TCF7L2
TCF7
RFX8
FOXA1
TP53
FOXM1
RFX1
RFX4
SNAI2
RFX2
RFX3
TCF21
CEBPA
ZBTB18
ZBTB42
MSC
ATOH1
LYL1
SOX11
SOX9
SOX17
ETV2
SOX13
ZNF695
SPDEF
SPIB
SPI1
SPIC
ETV6
NFATC4
LU0 LU1 LU2 LU3 LU4 LU5 LU6 LU7 LU8 LU9 LU10 LU11 LU12 LU13 LU14 LU15 LU16 LU17
LU10 LU0 LU8 LU6 LU4 LU3 LU14 LU15 LU1 LU7 LU2 LU12 LU11 LU13 LU9 LU17 LU5 LU16
Row Z−Scores
27516 Marker Peaks
−2
2
Normalized Motif Enrichment
in Marker Peaks
−log10(Padj) [0−Max]
0
100
e
TCF4
SNAI1
SNAI2
SNAI3
MESP1
ZEB1
ZEB2
HAND2
FIGLA
ZBTB18
TCF21
Variability
6
4
2
0
0
300
600
900
Rank Sorted Annotations
Variability
6
4
2
f
Total nFrags
100,000
10,000
1,000
100
### cells
10,000
1,000
100
10
1
Pearson's r
0.9
0.8
0.7
0.6
0.5
0.4
0.3
Jensen-Shannon distance
0.8
0.7
0.6
0.5
0.4
0.3
LU0 LU1 LU2 LU3 LU4 LU5 LU6 LU7 LU8 LU9 LU10 LU11 LU12 LU13 LU14 LU15 LU16 LU17

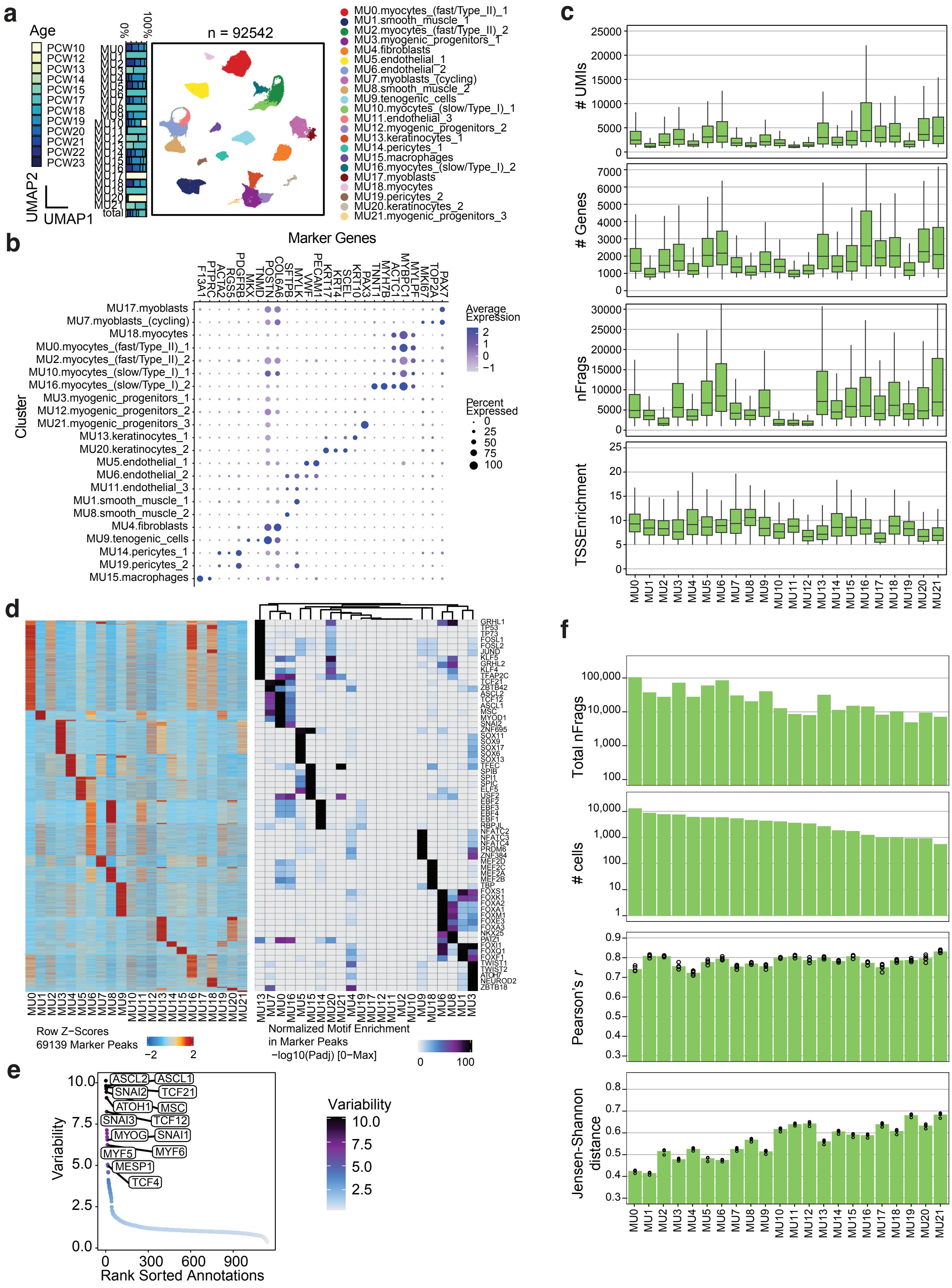
a
Age
PCW10
PCW12
PCW13
PCW14
PCW15
PCW17
PCW18
PCW19
PCW20
PCW21
PCW22
PCW23
0%
100%
total
n = 92542
UMAP2
UMAP1
MU0.myocytes_(fast/Type_II)_1
MU1.smooth_muscle_1
MU2.myocytes_(fast/Type_II)_2
MU3.myogenic_progenitors_1
MU4.fibroblasts
MU5.endothelial_1
MU6.endothelial_2
MU7.myoblasts_(cycling)
MU8.smooth_muscle_2
MU9.tenogenic_cells
MU10.myocytes_(slow/Type_I)_1
MU11.endothelial_3
MU12.myogenic_progenitors_2
MU13.keratinocytes_1
MU14.pericytes_1
MU15.macrophages
MU16.myocytes_(slow/Type_I)_2
MU17.myoblasts
MU18.myocytes
MU19.pericytes_2
MU20.keratinocytes_2
MU21.myogenic_progenitors_3
b
Marker Genes
F13A1
PTPRC
ACTA2
RGS5
PDGFRB
MKX
TNMD
POSTN
COL6A6
SFTPB
MYLK
VWF
PECAM1
KRT17
KRT4
SCEL
KRT10
PAX3
TNNT1
MYH7B
MYBPC1
ACTC1
MYLPF
MKI67
TOP2A
PAX7
Cluster
Average Expression
2
1
0
−1
Percent Expressed
0
25
50
75
100
c
### UMIs
### Genes
nFrags
TSSEnrichment
d
Row Z−Scores
69139 Marker Peaks
−2
2
Normalized Motif Enrichment in Marker Peaks
−log10(Padj) [0−Max]
0
100
GRHL1
TP53
TP73
FOSL1
FOSL2
JUND
KLF5
GRHL2
KLF4
TFAP2C
TCF21
ZBTB42
ASCL2
TCF12
ASCL1
MSC
MYOD1
SNAI2
ZNF695
SOX11
SOX9
SOX17
SOX6
SOX13
TFEC
SPIB
SPI1
SPIC
ELF5
USF2
EBF2
EBF3
EBF4
EBF1
RBPJL
NFATC2
NFATC3
NFATC4
PRDM6
ZNF384
MEF2D
MEF2C
MEF2A
MEF2B
TBP
FOXS1
FOXK1
FOXA2
FOXA1
FOXM1
FOXE3
FOXA3
NKX25
PATZ1
FOXI1
FOXO1
FOXF1
TWIST1
TWIST2
ATOH7
NEUROD2
ZBTB18
e
Variability
Rank Sorted Annotations
ASCL2
ASCL1
SNAI2
TCF21
ATOH1
MSC
SNAI3
TCF12
MYOG
SNAI1
MYF5
MYF6
MESP1
TCF4
10.0
7.5
5.0
2.5
0.0
0
300
600
900
f
Total nFrags
### cells
Pearson's r
Jensen-Shannon distance

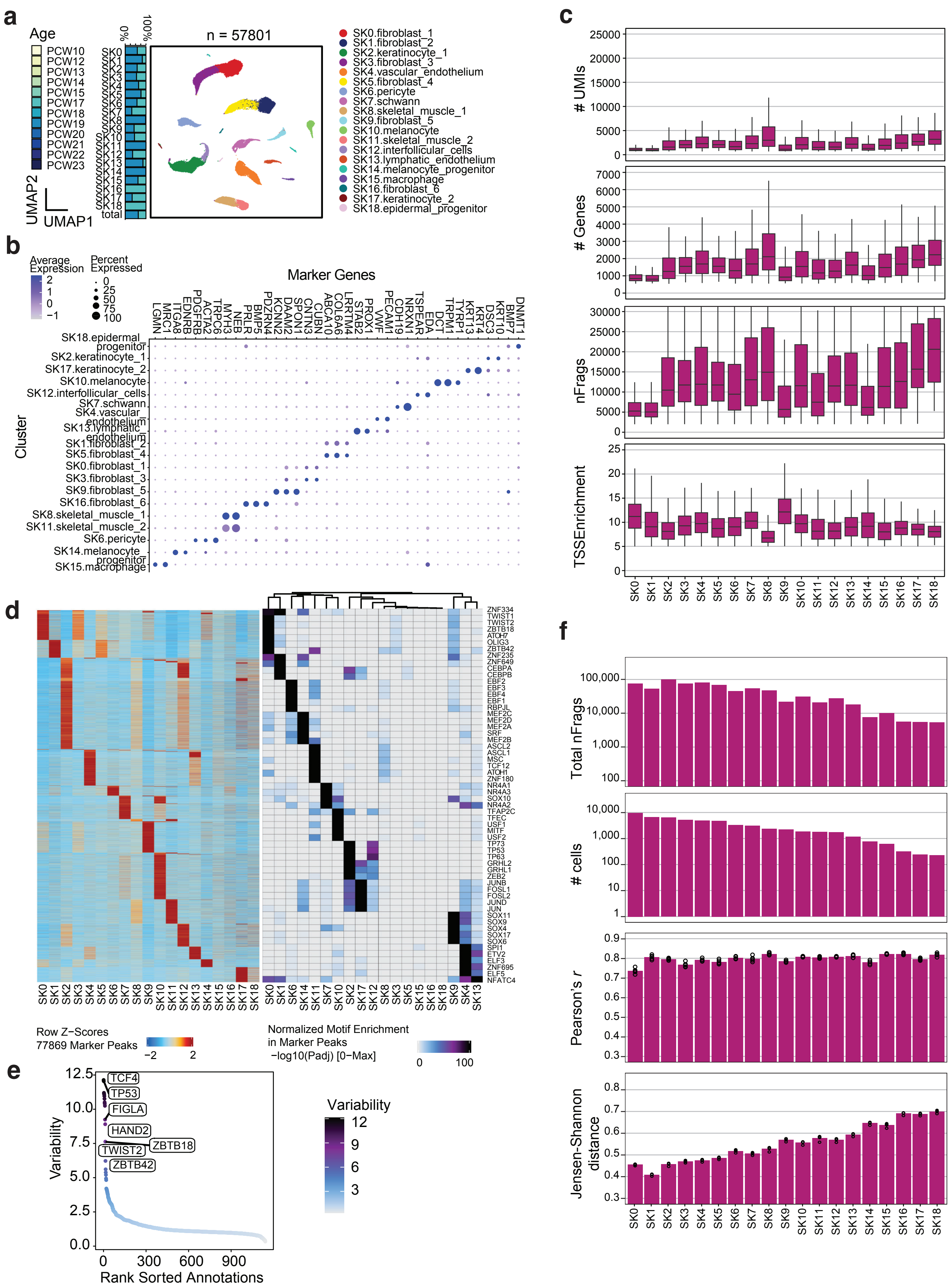

a
Age
PCW10
PCW12
PCW13
PCW14
PCW15
PCW17
PCW18
PCW19
PCW20
PCW21
PCW22
PCW23
0%
100%
n = 57801
UMAP2
UMAP1
SK0.fibroblast_1
SK1.fibroblast_2
SK2.keratinocyte_1
SK3.fibroblast_3
SK4.vascular_endothelium
SK5.fibroblast_4
SK6.pericyte
SK7.schwann
SK8.skeletal_muscle_1
SK9.fibroblast_5
SK10.melanocyte
SK11.skeletal_muscle_2
SK12.interfollicular_cells
SK13.lymphatic_endothelium
SK14.melanocyte_progenitor
SK15.macrophage
SK16.fibroblast_6
SK17.keratinocyte_2
SK18.epidermal_progenitor
b
Average Expression
Percent Expressed
Marker Genes
Cluster
LGMN MRC1 ITGA8 EDNRB PDGFRB ACTA2 TRPC6 MYH3 NEB PRLR BMP5 PDZRN4 KCNN2 DAAM2 SPON1 CNTN3 CUBN ABCA10 COL6A6 LRRTM4 STAB2 PROX1 VWF PECAM1 CDH19 NRXN1 TSPEAR EDA DCT TRPM1 TYRP1 KRT13 KRT4 DSC3 KRT10 BMP7 DNMT1
c
### UMIs
### Genes
nFrags
TSSEnrichment
d
Row Z-Scores
77869 Marker Peaks
Normalized Motif Enrichment
in Marker Peaks
−log10(Padj) [0−Max]
e
Variability
TCF4
TP53
FIGLA
HAND2
ZBTB18
TWIST2
ZBTB42
Rank Sorted Annotations
f
Total nFrags
### cells
Pearson's r
Jensen-Shannon distance

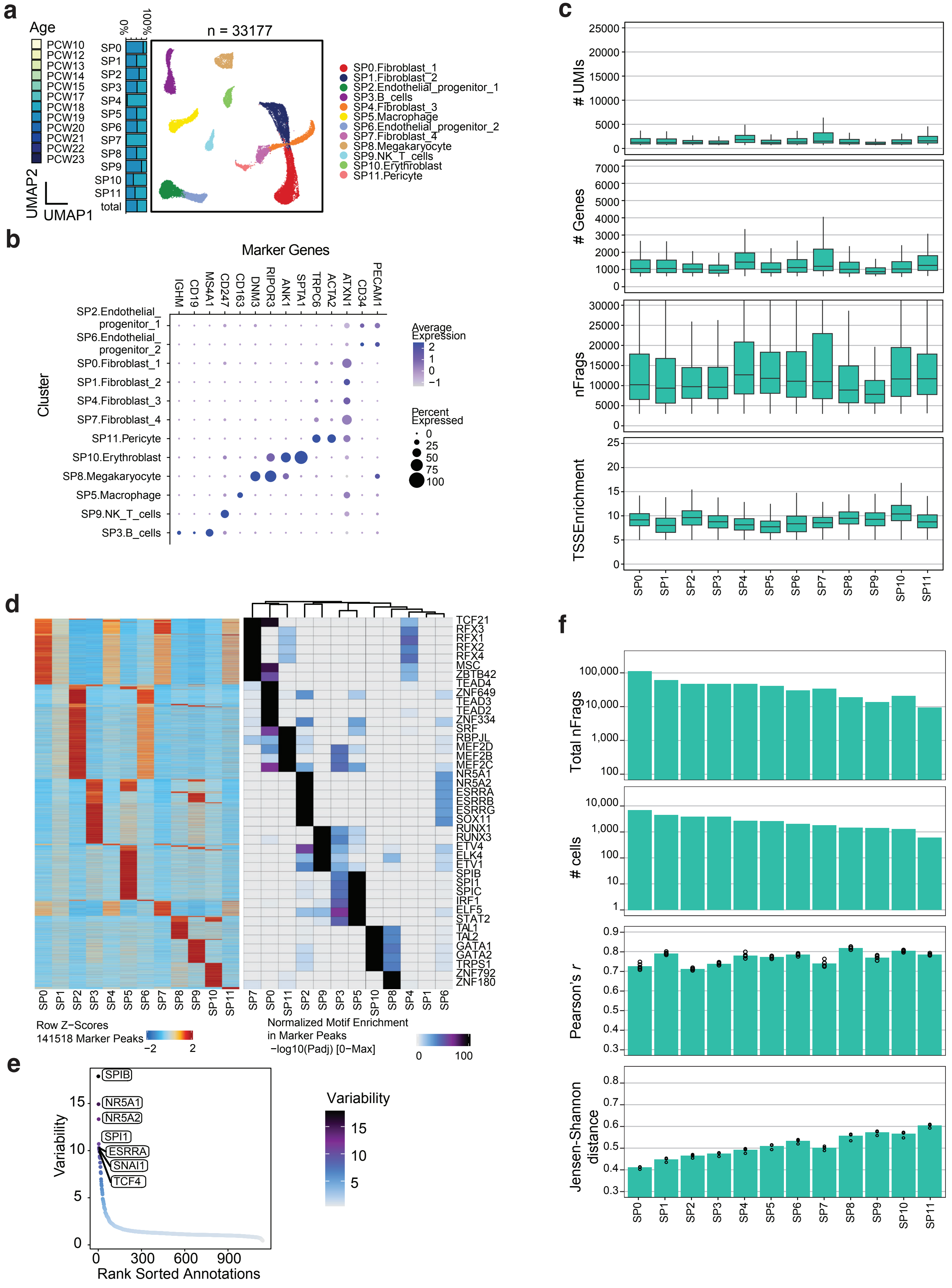

a
Age
PCW10
PCW12
PCW13
PCW14
PCW15
PCW17
PCW18
PCW19
PCW20
PCW21
PCW22
PCW23
0%
100%
SP0
SP1
SP2
SP3
SP4
SP5
SP6
SP7
SP8
SP9
SP10
SP11
total
n = 33177
UMAP2
UMAP1
SP0.Fibroblast_1
SP1.Fibroblast_2
SP2.Endothelial_progenitor_1
SP3.B_cells
SP4.Fibroblast_3
SP5.Macrophage
SP6.Endothelial_progenitor_2
SP7.Fibroblast_4
SP8.Megakaryocyte
SP9.NK_T_cells
SP10.Erythroblast
SP11.Pericyte
b
Marker Genes
IGHM
CD19
MS4A1
CD247
CD163
DNM3
RIPOR3
ANK1
SPTA1
TRPC6
ACTA2
ATXN1
CD34
PECAM1
Cluster
SP2.Endothelial_progenitor_1
SP6.Endothelial_progenitor_2
SP0.Fibroblast_1
SP1.Fibroblast_2
SP4.Fibroblast_3
SP7.Fibroblast_4
SP11.Pericyte
SP10.Erythroblast
SP8.Megakaryocyte
SP5.Macrophage
SP9.NK_T_cells
SP3.B_cells
Average Expression
2
1
0
−1
Percent Expressed
0
25
50
75
100
c
### UMIs
25000
20000
15000
10000
5000
0
### Genes
7000
6000
5000
4000
3000
2000
1000
0
nFrags
30000
25000
20000
15000
10000
5000
0
TSSEnrichment
25
20
15
10
5
0
SP0 SP1 SP2 SP3 SP4 SP5 SP6 SP7 SP8 SP9 SP10 SP11
d
TCF21
RFX3
RFX1
RFX2
RFX4
MSC
ZBTB42
TEAD4
ZNF649
TEAD3
TEAD2
ZNF334
SRF
RBPJL
MEF2D
MEF2B
MEF2C
NR5A1
NR5A2
ESRRA
ESRRB
ESRRG
SOX11
RUNX1
RUNX3
ETV4
ELK4
ETV1
SPIB
SPI1
SPIC
IRF1
ELF5
STAT2
TAL1
TAL2
GATA1
GATA2
TRPS1
ZNF792
ZNF180
SP0 SP1 SP2 SP3 SP4 SP5 SP6 SP7 SP8 SP9 SP10 SP11
SP7 SP0 SP11 SP2 SP9 SP3 SP5 SP10 SP8 SP4 SP1 SP6
Row Z−Scores
141518 Marker Peaks
−2
2
Normalized Motif Enrichment
in Marker Peaks
−log10(Padj) [0−Max]
0
100
e
Variability
SPIB
NR5A1
NR5A2
SPI1
ESRRA
SNAI1
TCF4
15
10
5
0
0
300
600
900
Rank Sorted Annotations
Variability
15
10
5
f
Total nFrags
100,000
10,000
1,000
100
### cells
10,000
1,000
100
10
1
Pearson's r
0.9
0.8
0.7
0.6
0.5
0.4
0.3
Jensen-Shannon distance
0.8
0.7
0.6
0.5
0.4
0.3
SP0 SP1 SP2 SP3 SP4 SP5 SP6 SP7 SP8 SP9 SP10 SP11

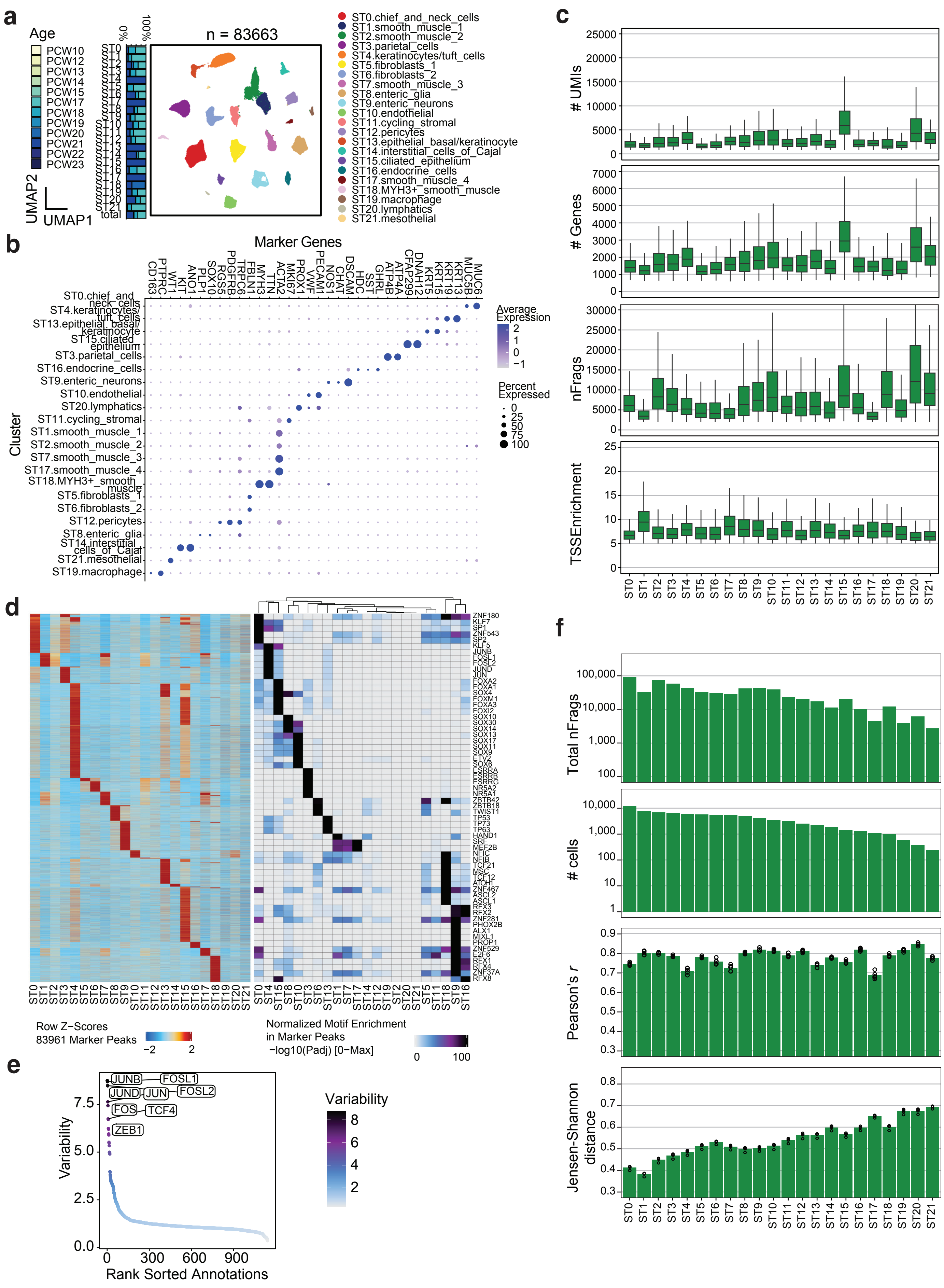

a
Age
PCW10
PCW12
PCW13
PCW14
PCW15
PCW17
PCW18
PCW19
PCW20
PCW21
PCW22
PCW23
0%
100%
n = 83663
UMAP2
UMAP1
ST0.chief_and_neck_cells
ST1.smooth_muscle_1
ST2.smooth_muscle_2
ST3.parietal_cells
ST4.keratinocytes/tuft_cells
ST5.fibroblasts_1
ST6.fibroblasts_2
ST7.smooth_muscle_3
ST8.enteric_glia
ST9.enteric_neurons
ST10.endothelial
ST11.cycling_stromal
ST12.pericytes
ST13.epithelial_basal/keratinocyte
ST14.interstitial_cells_of_Cajal
ST15.ciliated_epithelium
ST16.endocrine_cells
ST17.smooth_muscle_4
ST18.MYH3+_smooth_muscle
ST19.macrophage
ST20.lymphatics
ST21.mesothelial
b
Marker Genes
Cluster
Average Expression
Percent Expressed
c
### UMIs
### Genes
nFrags
TSSEnrichment
d
Row Z-Scores
83961 Marker Peaks
Normalized Motif Enrichment in Marker Peaks
−log10(Padj) [0−Max]
e
Variability
Rank Sorted Annotations
JUNB
FOSL1
JUND
JUN
FOSL2
FOS
TCF4
ZEB1
f
Total nFrags
### cells
Pearson's r
Jensen-Shannon distance

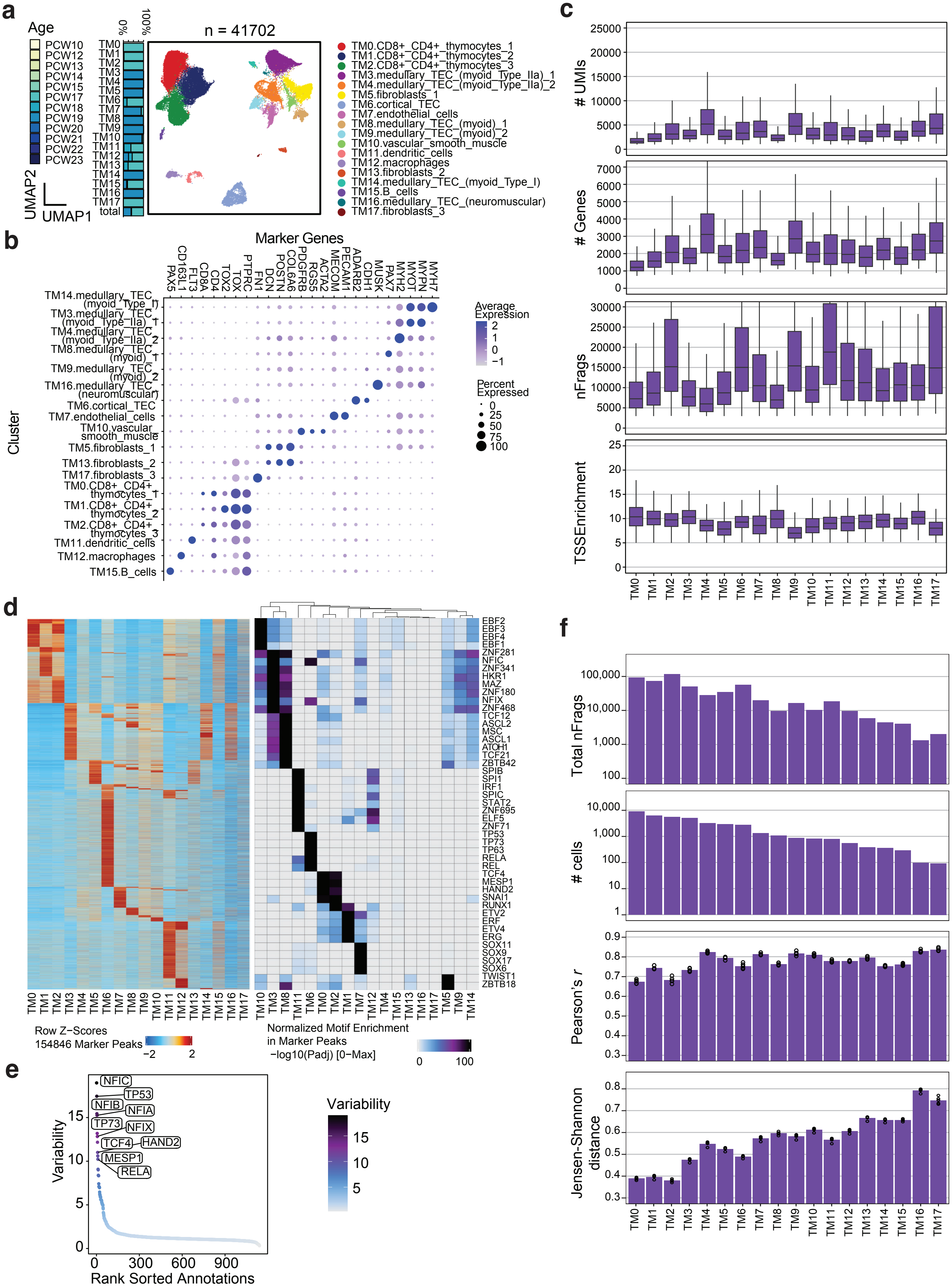
a
Age
PCW10
PCW12
PCW13
PCW14
PCW15
PCW17
PCW18
PCW19
PCW20
PCW21
PCW22
PCW23
0%
100%
TM0
TM1
TM2
TM3
TM4
TM5
TM6
TM7
TM8
TM9
TM10
TM11
TM12
TM13
TM14
TM15
TM16
TM17
total
UMAP2
UMAP1
n = 41702
TM0.CD8+_CD4+_thymocytes_1
TM1.CD8+_CD4+_thymocytes_2
TM2.CD8+_CD4+_thymocytes_3
TM3.medullary_TEC_(myoid_Type_IIa)_1
TM4.medullary_TEC_(myoid_Type_IIa)_2
TM5.fibroblasts_1
TM6.cortical_TEC
TM7.endothelial_cells
TM8.medullary_TEC_(myoid)_1
TM9.medullary_TEC_(myoid)_2
TM10.vascular_smooth_muscle
TM11.dendritic_cells
TM12.macrophages
TM13.fibroblasts_2
TM14.medullary_TEC_(myoid_Type_I)
TM15.B_cells
TM16.medullary_TEC_(neuromuscular)
TM17.fibroblasts_3
b
Marker Genes
PAX5
CD163L1
FLT3
CD8A
CD4
TOX2
TOX
PTPRC
FN1
DCN
POSTN
COL6A6
PDGFRB
RGS5
ACTA2
MECOM
PECAM1
ADARB2
CDH1
MUSK
PAX7
MYH2
MYOT
MYPN
MYH7
Cluster
TM14.medullary_TEC (myoid_Type_I)
TM3.medullary_TEC (myoid_Type_IIa)_1
TM4.medullary_TEC (myoid_Type_IIa)_2
TM8.medullary_TEC (myoid)_1
TM9.medullary_TEC (myoid)_2
TM16.medullary_TEC (neuromuscular)
TM6.cortical_TEC
TM7.endothelial_cells
TM10.vascular_smooth_muscle
TM5.fibroblasts_1
TM13.fibroblasts_2
TM17.fibroblasts_3
TM0.CD8+_CD4+_thymocytes_1
TM1.CD8+_CD4+_thymocytes_2
TM2.CD8+_CD4+_thymocytes_3
TM11.dendritic_cells
TM12.macrophages
TM15.B_cells
Average Expression
2
1
0
−1
Percent Expressed
0
25
50
75
100
c
### UMIs
### Genes
nFrags
TSSEnrichment
d
EBF2
EBF3
EBF4
EBF1
ZNF281
NFIC
ZNF341
HKR1
MAZ
ZNF180
NFIX
ZNF468
TCF12
ASCL2
MSC
ASCL1
ATOH1
TCF21
ZBTB42
SPIB
SPI1
IRF1
SPIC
STAT2
ZNF695
ELF5
ZNF71
TP53
TP73
TP63
RELA
REL
TCF4
MESP1
HAND2
SNAI1
RUNX1
ETV2
ERF
ETV4
ERG
SOX11
SOX9
SOX17
SOX6
TWIST1
ZBTB18
Row Z−Scores
154846 Marker Peaks
−2
2
Normalized Motif Enrichment in Marker Peaks
−log10(Padj) [0−Max]
0
100
e
NFIC
TP53
NFIB
NFIA
TP73
NFIX
TCF4
HAND2
MESP1
RELA
Variability
Rank Sorted Annotations
f
Total nFrags
### cells
Pearson's r
Jensen-Shannon distance

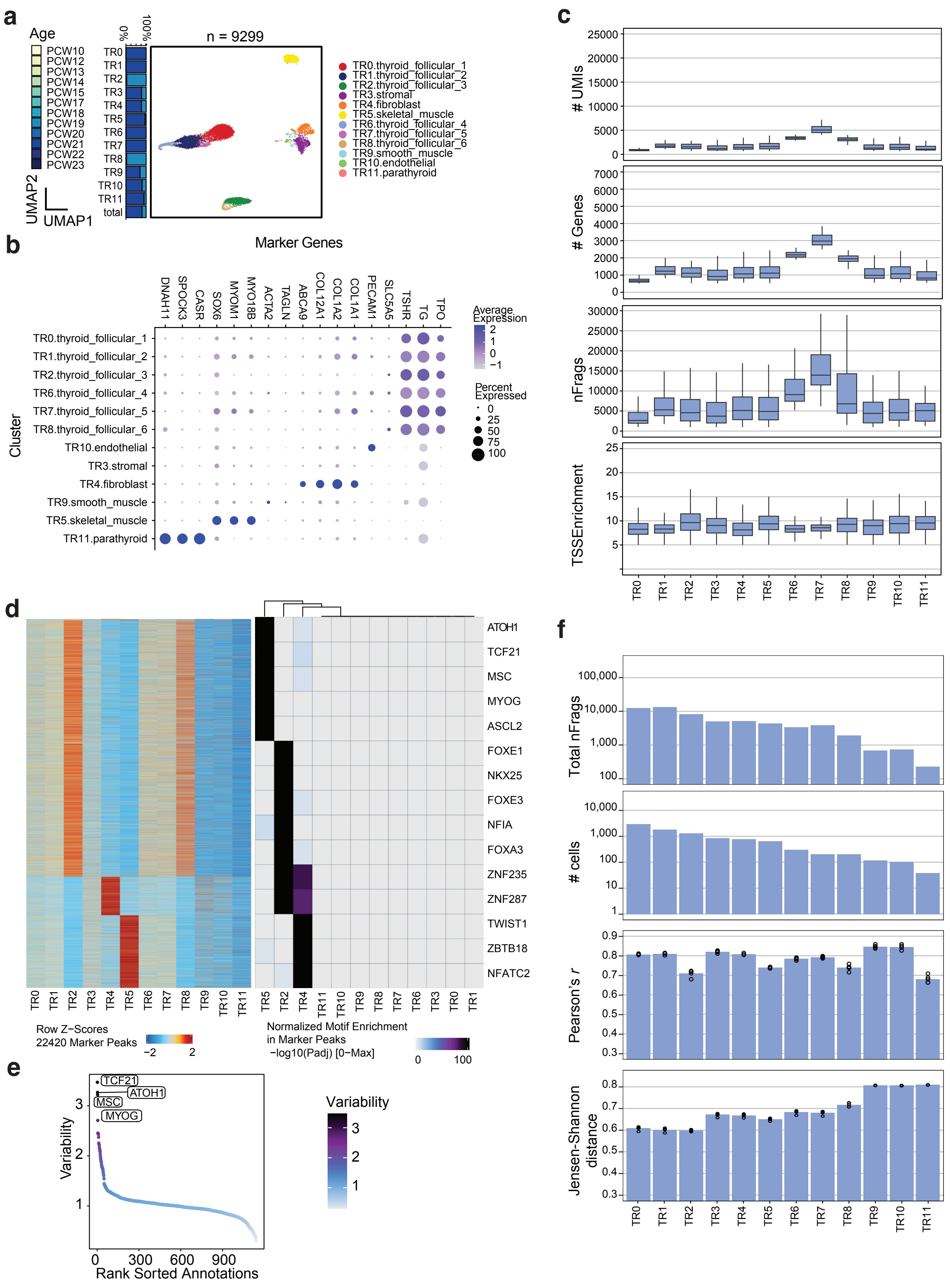
a
Age
PCW10
PCW12
PCW13
PCW14
PCW15
PCW17
PCW18
PCW19
PCW20
PCW21
PCW22
PCW23
0%
100%
TR0
TR1
TR2
TR3
TR4
TR5
TR6
TR7
TR8
TR9
TR10
TR11
total
UMAP2
UMAP1
n = 9299
TR0.thyroid_follicular_1
TR1.thyroid_follicular_2
TR2.thyroid_follicular_3
TR3.stromal
TR4.fibroblast
TR5.skeletal_muscle
TR6.thyroid_follicular_4
TR7.thyroid_follicular_5
TR8.thyroid_follicular_6
TR9.smooth_muscle
TR10.endothelial
TR11.parathyroid
b
Marker Genes
DNAH11
SPOCK3
CASR
SOX6
MYOM1
MYO18B
ACTA2
TAGLN
ABCA9
COL12A1
COL1A2
COL1A1
PECAM1
SLC5A5
TSHR
TG
TPO
Cluster
TR0.thyroid_follicular_1
TR1.thyroid_follicular_2
TR2.thyroid_follicular_3
TR6.thyroid_follicular_4
TR7.thyroid_follicular_5
TR8.thyroid_follicular_6
TR10.endothelial
TR3.stromal
TR4.fibroblast
TR9.smooth_muscle
TR5.skeletal_muscle
TR11.parathyroid
Average Expression
2
1
0
−1
Percent Expressed
0
25
50
75
100
c
### UMIs
25000
20000
15000
10000
5000
0
### Genes
7000
6000
5000
4000
3000
2000
1000
0
nFrags
30000
25000
20000
15000
10000
5000
0
TSSEnrichment
25
20
15
10
5
TR0
TR1
TR2
TR3
TR4
TR5
TR6
TR7
TR8
TR9
TR10
TR11
d
ATOH1
TCF21
MSC
MYOG
ASCL2
FOXE1
NKX25
FOXE3
NFIA
FOXA3
ZNF235
ZNF287
TWIST1
ZBTB18
NFATC2
TR0
TR1
TR2
TR3
TR4
TR5
TR6
TR7
TR8
TR9
TR10
TR11
TR5
TR2
TR4
TR11
TR10
TR9
TR8
TR7
TR6
TR3
TR0
TR1
Row Z−Scores
22420 Marker Peaks
−2
2
Normalized Motif Enrichment
in Marker Peaks
−log10(Padj) [0−Max]
0
100
e
TCF21
ATOH1
MSC
MYOG
Variability
3
2
1
0
300
600
900
Rank Sorted Annotations
Variability
3
2
1
f
Total nFrags
100,000
10,000
1,000
100
### cells
10,000
1,000
100
10
1
Pearson's r
0.9
0.8
0.7
0.6
0.5
0.4
0.3
Jensen-Shannon distance
0.8
0.7
0.6
0.5
0.4
0.3
TR0
TR1
TR2
TR3
TR4
TR5
TR6
TR7
TR8
TR9
TR10
TR11
